# Competing Inter-Domain Quorum-Sensing Systems Control Prophage Lysis-Lysogeny Decisions

**DOI:** 10.64898/2026.07.02.736140

**Authors:** Michael L. Bunsick, Gabriel D. D’Agostino, Thu Vu Phuc Nguyen, Francis J. Santoriello, Emilee E. Shine, Bonnie L. Bassler

**Affiliations:** Department of Molecular Biology, Princeton University, Princeton, New Jersey, United States of America; Howard Hughes Medical Institute, Chevy Chase, Maryland, United States of America

**Keywords:** bacteriophage, communication, quorum sensing, lysogeny

## Abstract

*Phaeobacter inhibens* T5^T^ harbors three LuxI/LuxR quorum-sensing systems: one encoded by the host and one in each of two prophages. Each prophage quorum-sensing autoinducer activates its own lysogeny program and that of the co-resident prophage. By driving competitors toward lysogeny, prophages deny their rivals access to susceptible hosts. One prophage engages in a bidirectional interaction with the host: the prophage activates host quorum sensing, while the host represses prophage quorum sensing. Because the host and prophage autoinducers exert opposing effects on the same prophage promoter, the prophage’s lysis-lysogeny decision depends on the ratio of the two autoinducers rather than their absolute concentrations. Ratiometric sensing allows the prophage to infer the relative abundances of infected and uninfected hosts and to commit to lysis only when uninfected hosts predominate. Phage quorum-sensing modules are widespread and undergo diversification and exchange. These findings reveal how prophages surveil and manipulate competitors that share chemical environments.

## Introduction

Bacterial viruses (phages) face a critical decision when infecting a host cell. They can either reproduce immediately and lyse their host or persist as prophages in a dormant state known as lysogeny^1,2,3^. The reproductive payoff of this choice depends on both the conditions within the host and the composition of the surrounding bacterial population. At the level of a single host cell, the number of progeny virions a phage can produce is capped by the infected cell’s metabolic capacity and constrained by competition from co-resident prophages^4,5,6^. At the bacterial population level, any further phage reproduction requires the released virions to encounter new, susceptible hosts. Because phages usually cannot infect cells already carrying a homologous prophage, the number of uninfected host cells in the surrounding population dictates the payoff from lytic activity^7,8^. Premature lysis releases virions into a bacterial population with limited host availability, while prolonged lysogeny in a host-rich environment squanders reproductive opportunity.

Phages navigate this reproductive trade-off using signaling pathways that link sensory cues to their lysis-lysogeny decision making machinery. Commonly, stresses such as DNA damage activate the bacterial SOS response, components of which inactivate prophage repressors of lytic gene expression^2,3^. This mechanism provides a reliable means for phages to escape from compromised hosts, thus preserving the phage’s reproductive opportunities. Because the SOS response is conserved across bacteria, it serves as a general mechanism driving prophage lytic induction^2,3^. Although this mechanism reliably reports on host viability, it does not provide information to the phage about host availability, host susceptibility, or phage-phage competition.

Beyond host-intrinsic stress signals, phages can exploit social cues in bacterial communities by surveilling host quorum-sensing (QS) signals. QS is a bacterial chemical communication process that relies on the production, release, accumulation, and group-wide detection of extracellular signal molecules called autoinducers^9,10^. QS enables bacteria to orchestrate collective behaviors. Some prophages can monitor QS autoinducers and use the information they garner to regulate their lysis-lysogeny transitions. For example, a receptor from the vibriophage VP882 binds a host-produced autoinducer and launches the phage lytic program at high host cell density^11^. This mechanism enables phage VP882 to preferentially produce virions when surrounded by other host cells, presumably maximizing chances of successful transmission. Similarly, *Bacillus* phages can also gauge population density using arbitrium peptides^12,13^. These peptides reflect prophage density and bias phages toward lysogeny when already infected hosts dominate the population.

Although recent work has revealed mechanisms that enable phages to produce and respond to individual autoinducers, how such processes operate in environments with multiple coexisting QS systems remains unexplored. These environments become especially intriguing when the QS systems are harbored by organisms with competing evolutionary interests. Such cases include two co-resident prophages or a phage and its bacterial host. In these situations, QS signals become public resources that different players can monitor and manipulate to overcome their competitors. To explore whether phages can exploit this richer signaling landscape, we examined a naturally occurring polylysogen in which multiple bacterial and prophage QS systems coexist. This polylysogen, *Phaeobacter inhibens* T5^T^, carries three LuxI/LuxR QS circuits: one encoded by the host and one encoded by each of two chromosomally-integrated prophages^14^. This configuration provides a minimal ecosystem in which we can probe signaling interactions between the host and both co-resident phages.

We show that in *P. inhibens* T5^T^, each prophage produces a homoserine lactone QS autoinducer that promotes its own lysogeny. This mechanism reinforces dormancy when phage genomes are abundant. Each prophage can also activate the other prophage’s QS system, biasing its competitor toward lysogeny. By suppressing rival prophage replication, the autoinducer-producing prophage should gain a competitive advantage. Beyond phage-to-phage cross-activation, we show that one of the prophages engages in a bidirectional interaction with the host QS system: the prophage autoinducer activates host QS, and the host autoinducer suppresses QS in the prophage. This interaction drives the infected host to initiate QS behaviors at lower cell densities than do uninfected cells. For the prophage, the host autoinducer promotes lytic induction, while its own autoinducer promotes lysogeny maintenance. The opposing effects of the two autoinducers render the prophage’s lysis-lysogeny switch sensitive to the ratio of the two chemicals. We propose that the phage uses this ratiometric information to infer the relative abundances of infected and uninfected host cells in the population. When the host:prophage autoinducer ratio is high, the phage intuits that uninfected hosts dominate the population and it triggers the lytic cascade. When the host:prophage autoinducer ratio is low, the phage deduces that prophage-infected host cells predominate and it maintains lysogeny. This strategy could enable the phage to maximize transmission opportunities while limiting costly, unproductive competition with other lysogens. Together, our findings demonstrate how QS enables phages to collect and manipulate sensory information to outmaneuver both hosts and competing prophages.

## Results

### The *Phaeobacter inhibens* T5^T^ polylysogen possesses one host and two prophage QS systems

To learn how phages interpret QS signals in authentic environments, we sought a natural system in which multiple QS circuits coexist. We focused on LuxI/LuxR QS systems because they produce well-defined acyl-homoserine lactone (HSL) autoinducers and because their component genes are readily identifiable in bacterial genomes^8^. By these criteria, the marine bacterium *P. inhibens* T5^T^ provides an ideal system (Fig. 1A)^14^. The host chromosome harbors a single *luxI/luxR* pair, previously named *pgaI/pgaR*^15^. The *P. inhibens* T5^T^ genome also contains two unrelated prophages, each harboring a *luxI/luxR* gene pair. One prophage, which we designate Phage A, belongs to the Mu family and encodes the transposition machinery characteristic of this group (Fig. 1B). The second phage, called Phage B, is a myovirus-like prophage related to *Escherichia coli* phage vB_EcoM-ep3 (Fig. 1B).

**Figure 1.**
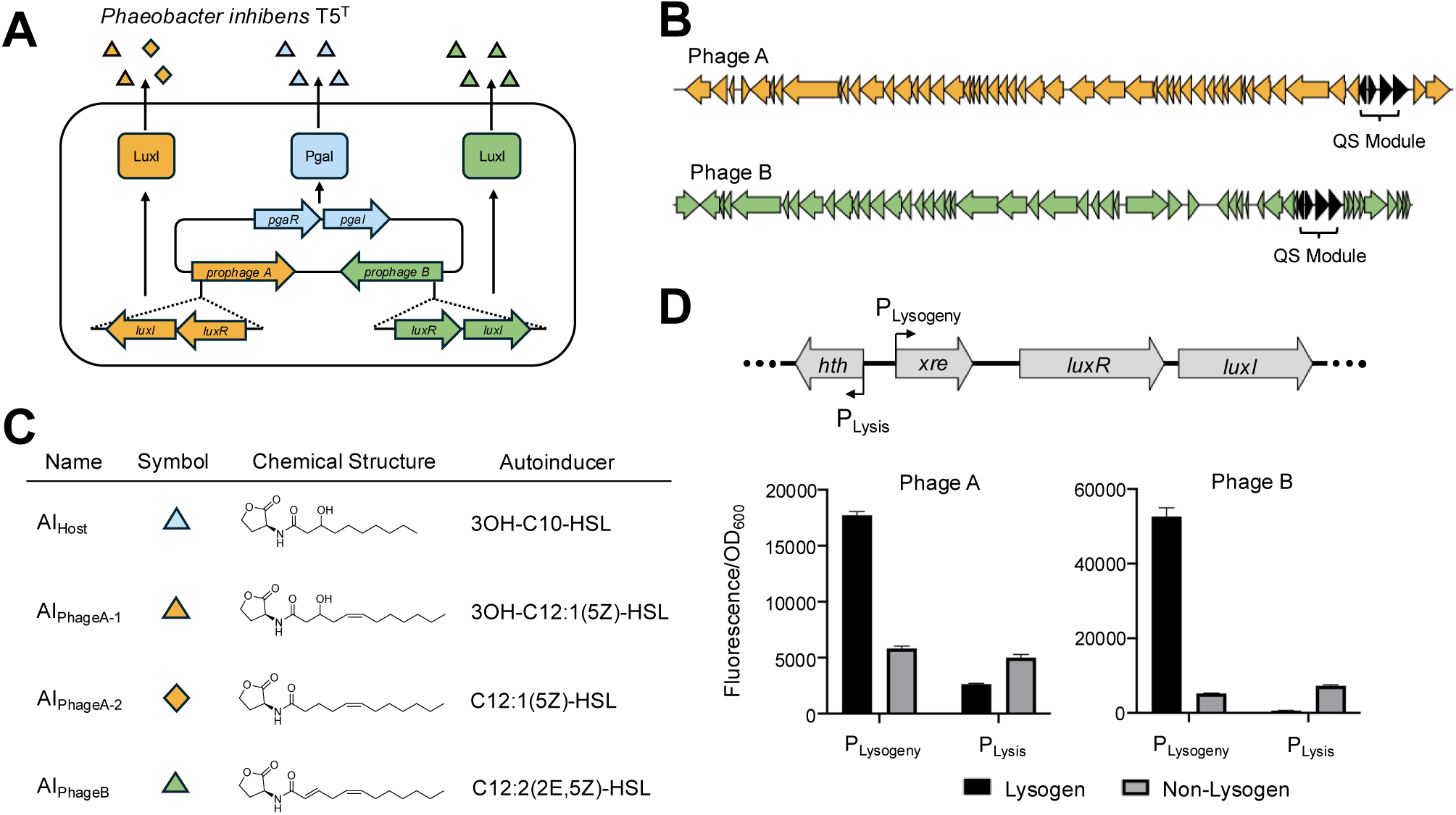
*P. inhibens* T5^T^ harbors three QS systems that produce unique autoinducers. **(A)** Schematic of the *P. inhibens* T5^T^ QS network. The host (blue), Phage A (orange), and Phage B (green) each encode a LuxI/LuxR-type QS system. The three LuxI synthases produce distinct autoinducers (triangles and diamonds). **(B)** Genetic organization of Phage A (orange) and Phage B (green). Arrows indicate the directions of transcription. The four gene QS modules (black arrows), containing the *luxR* and *luxI* genes, are bracketed. **(C)** Structures and identities of the four autoinducers produced by the three *P. inhibens* T5^T^ QS systems. Symbols and color codes are used throughout this work. **(D)** Top: Organization of *P. inhibens* T5^T^ prophage QS modules. Bottom: Phage A (left) and Phage B (right) P_Lysogeny_ and P_Lysis_ *mScarlet-I* outputs, measured in lysogenic (black) and non-lysogenic (gray) *P. inhibens* T5^T^ strains. mScarlet-I fluorescence is normalized to OD_600_. Data represent means ± SDs of four biological replicates.

The three *P. inhibens* T5^T^ LuxI/LuxR pairs share less than 30% amino-acid identity. This level of sequence divergence suggested that each QS circuit might produce a distinct HSL, a precondition for selective cross-regulation. To test this possibility, we individually overexpressed each *luxI* gene in a *P. inhibens* strain lacking the two QS prophages. We analyzed cell-free culture fluids using UPLC-MS. Consistent with previous reports, the host enzyme, PgaI, produced 3OH-C10-HSL (AI_Host_) (Fig. 1C)^15,16^. We identified two HSLs associated with the Phage A-encoded LuxI and one HSL made by the Phage B-encoded LuxI (see Methods). Structural analyses combining MS/MS and NMR, corroborated by chemical derivatization and comparison to synthetic standards, established their identities. Phage A LuxI produces 3OH-C12:1(5Z)-HSL (AI_PhageA-1_) and C12:1(5Z)-HSL (AI_PhageA-2_) (Fig. 1C). Phage B LuxI produces C12:2(2E,5Z)-HSL (AI_PhageB_) (Fig. 1C). We quantified the individual autoinducers by UPLC-MS. At stationary phase (growth for 16 h at 28°C) AI_Host_ = 579 nM; AI_PhageA-1_ = 73 nM; AI_PhageA-2_ = 14 nM; AI_PhageB_ = 142 nM (Fig. S1A). These results show that all three LuxI proteins are active autoinducer synthases and that the host and each prophage produce non-identical autoinducers.

Given their divergence in autoinducer production, we wondered whether the *P. inhibens* T5^T^ QS systems also differ in their genomic organization. We found that the host *luxI/luxR* genes are arranged in a simple two-gene operon. By contrast, both phage *luxI/luxR* pairs reside within four-gene modules that have identical synteny (Fig. 1B, 1D (top)). In both prophages, the *luxR* and *luxI* genes form a three-gene operon together with an upstream gene encoding an XRE-family transcription factor. 5’ to this operon lies a counter-oriented gene encoding a predicted helix-turn-helix DNA-binding protein, which we designate *hth*. Notably, although the overall architectures of the Phage A and Phage B QS modules are conserved between the two prophages, the encoded proteins show low sequence identity.

Neither prophage genome contains recognizable genes specifying homologs of the canonical regulatory proteins typically associated with lysis–lysogeny control, such as the cI/Cro proteins in Phage lambda or Ner/C in Mu-like phages. The absence of these typical lysis-lysogeny regulators suggests that the components encoded in the four-gene QS modules may themselves serve as the switches controlling the phages’ lifestyle transitions. A structural feature of these module supports this interpretation: divergent promoters drive the expression of the *hth* genes and the *xre–luxR–luxI* operons. This arrangement is identical to that underlying the genetic switch in Phage lambda. In the lambda system, divergent promoters drive reciprocal repression of *cI* and *cro*: cI maintains lysogeny by repressing *cro*, whereas Cro promotes lysis by repressing *cI*^2^. Based on this identical architecture, we predicted that each phage’s *xre–luxR–luxI* operon would be expressed during lysogeny, while its *hth* gene would be expressed during lysis. To test this supposition, we fused each promoter to *mScarlet-I* and measured reporter activity in strains carrying or lacking the cognate prophage. In both cases, the promoter driving *xre–luxR–luxI* was strongly expressed in lysogenic cells but inactive in cells lacking the prophage encoding it. In contrast, the counter-oriented *hth* promoter showed the opposite behavior, being expressed in the absence of the prophage and strongly repressed in lysogens (Fig. 1D (bottom)). We therefore designate the prophage promoters upstream of the *xre-luxR-luxI* operons P_Lysogeny_ and the counter-oriented promoters upstream of the *hth* genes P_Lysis_. Together, these results suggest the conserved four-gene modules function as QS-responsive genetic switches. Because the prophage *luxR* genes sit within their P_Lysogeny_ operons, autoinducer activation of LuxR should generate a positive feedback loop that drives commitment to lysogeny.

### Identification of the targets and regulatory logic of the three *P. inhibens* T5^T^ QS systems

Our model predicts that LuxR-dependent activation of P_Lysogeny_ maintains expression of the *xre-luxR-luxI* operons in lysogens. To test this assertion, we constructed reporter plasmids in which the P_Lysogeny_ promoter from Phage A or Phage B drives expression of *mScarlet-I* inserted upstream of the *xre–luxR–luxI* operon. As controls, we built constructs containing the P_Lysogeny_ promoters upstream of only *mScarlet-I*. We individually introduced the reporter plasmids into a *P. inhibens* strain lacking all the QS *luxI* and *luxR* genes. We call this strain the *P. inhibens* Δ3*luxI* Δ3*luxR* strain. This design allowed us to assess whether each phage LuxR activates its own P_Lysogeny_ promoter. Strong *mScarlet-I* fluorescence occurred in the strains harboring the plasmids carrying the full *xre–luxR–luxI* operons but not in the controls (Fig. 2A, B). These data indicate that the *xre-luxR-luxI* operons are sufficient to activate their cognate P_Lysogeny_ promoters.

**Figure 2.**
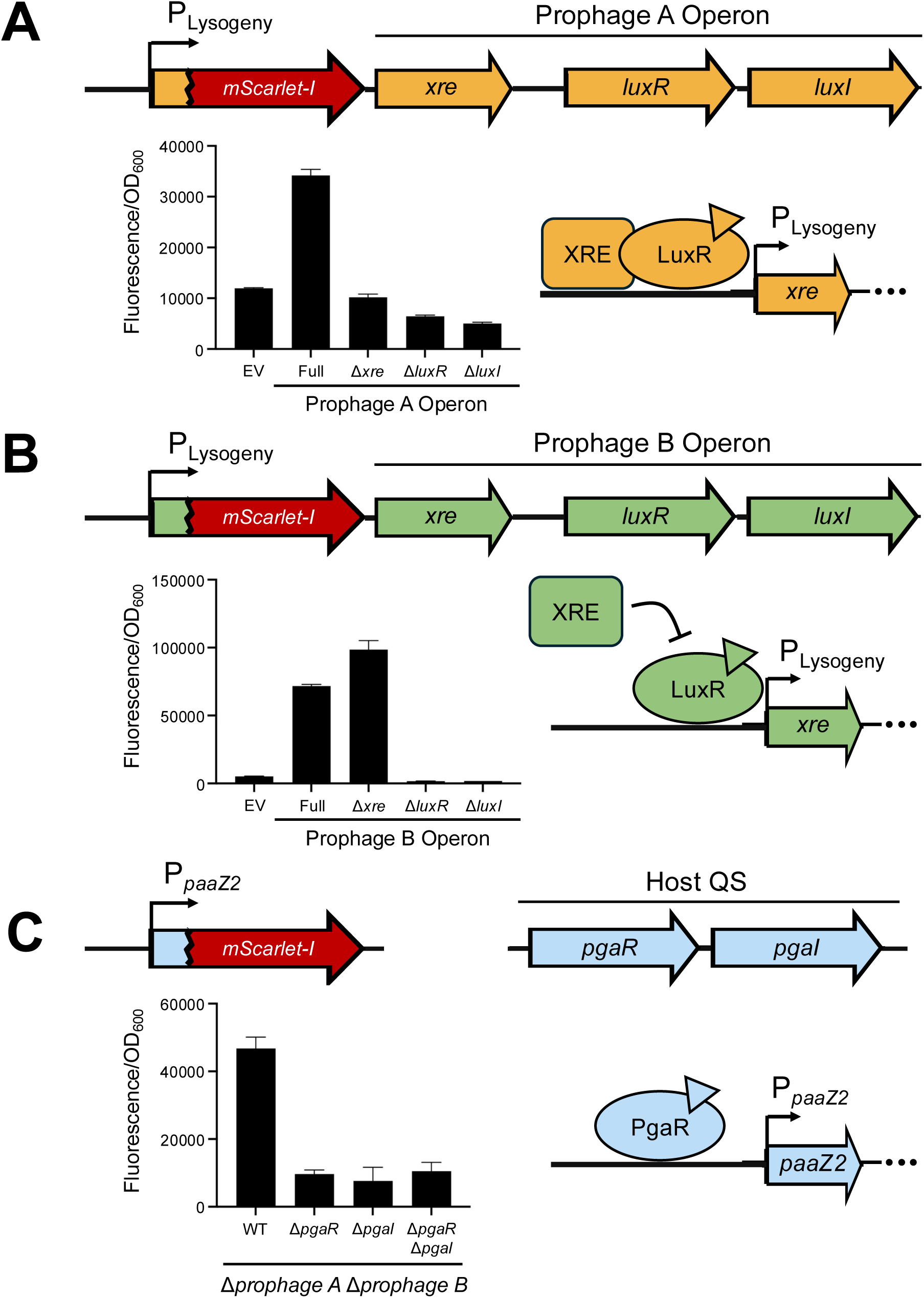
The *P. inhibens* T5^T^ QS systems control their target promoters using distinct regulatory mechanisms. Schematics of reporter constructs, their outputs, and the corresponding regulatory arrangements are shown. (**A**) Phage A P_Lysogeny_ reporter activity was measured in the Δ*3luxI* Δ*3luxR P. inhibens* strain. EV denotes the P_Lysogeny_–*mScarlet-I* reporter lacking the downstream prophage genes. Full denotes the entire *xre–luxR–luxI* operon in the reporter downstream of P_Lysogeny_–*mScarlet* as shown in the schematic. (**B**) As in panel A for the Phage B. (**C**) P*_paaZ2_*-*mScarlet-I* reporter activity was measured in the Δ*prophage A* Δ*prophage B P. inhibens* strain (WT) and the strain containing the genomic deletions indicated on the x-axis. (A-C) mScarlet-I fluorescence is normalized to OD_600_. Data represent the means ± SDs of four biological replicates.

To define the minimal requirements for P_Lysogeny_ activation, we assessed the contributions of the individual genes within the *xre-luxR-luxI* operons. To this end, we individually deleted *xre*, *luxR*, or *luxI* in our reporter plasmids and measured the resulting fluorescence output for each construct. For Phage A and Phage B, expression from P_Lysogeny_ required both *luxI* and *luxR*, consistent with the canonical LuxI-LuxR signaling mechanism (Fig. 2A, B). In line with this result, we could restore expression from P_Lysogeny_ in the Δ*luxI* reporter for Phage A by adding either of its cognate autoinducers AI_PhageA-1_ (EC_50_ = 0.71 nM) or AI_PhageA-2_ (EC_50_ = 3.2 μM) (Fig. S1B). Likewise, we restored expression from the Phage B P_Lysogeny_ promoter in the construct lacking *luxI* by adding AI_PhageB_ (EC_50_ = 1.1 nM) (Fig. S1B).

Despite these similarities, the two prophage QS systems differed in their dependence on the XRE proteins. In Phage A, deletion of *xre* abolished expression from P_Lysogeny_ indicating that the Phage A *xre-luxR-luxI* operon requires XRE for activation (Fig. 2A). This regulatory architecture mirrors that recently reported for phage QS circuits in which a LuxR-type receptor functions in conjunction with an XRE-family transcription factor to activate target promoters^6,17^. In contrast, deletion of *xre* from the Phage B construct led to a modest increase in P_Lyosgeny_ expression (Fig. 2B). Thus, the Phage B XRE protein suppresses P_Lysogeny_ transcription. Together, these results show that the two prophage QS systems share the same components but assemble them into different regulatory architectures.

To complete our analysis of the three coexisting *P. inhibens* T5^T^ QS systems, we examined the host-encoded PgaI/PgaR circuit. To probe function, we constructed a plasmid that reports on host QS activity using the *paaZ2* promoter. PaaZ2 encodes an enzyme in the tropodithietic acid (TDA) antibiotic biosynthetic pathway and was previously shown to be regulated by PgaI/PgaR^15,18,19^. We fused the *paaZ2* promoter to *mScarlet-I* and measured fluorescence in strains lacking both prophages and containing (designated WT for wildtype) or lacking *pgaI* or *pgaR*. Deleting either *pgaI* or *pgaR* abolished *mScarlet-I* expression, confirming that activation of the *paaZ2* promoter requires both the PgaR receptor and its cognate PgaI-produced autoinducer (Fig. 2C, Fig. S1B demonstrates restoration with AI_Host_, EC_50_ = 0.8 nM). Beyond showing that the host PgaI/PgaR QS system functions, these results establish the *paaZ2* promoter as a reliable target to read out host QS activity.

### Cross-regulation between host and prophage QS systems

Having established that all three *P. inhibens* T5^T^ LuxI/LuxR QS systems function as expected, we tested possible interactions. To do this, we exploited the reporter constructs described above that lack the *luxI* genes in our Δ3*luxI* Δ3*luxR* strain. Because the constructs produce the LuxR receptors but not their cognate LuxI synthases, reporter activation can only occur in response to externally supplied autoinducers. This arrangement allows us to measure the response of individual LuxR proteins to non-cognate autoinducers.

#### Phage to phage cross-regulation

To assess cross-regulation between the two prophage QS systems, we supplied each prophage autoinducer to Δ3*luxI* Δ3*luxR* strains carrying the opposing prophage’s Δ*luxI* reporter construct. AI_PhageB_ activated transcription from the Phage A P_Lysogeny_ reporter (EC_50_ = 119.4 nM) and, likewise, AI_PhageA-2_ induced the Phage B P_Lysogeny_ reporter (EC_50_ = 75.8 nM), however, AI_PhageA-1_ did not (Fig. 3A, Fig. S2). Thus, each prophage can detect an autoinducer produced by the other.

**Figure 3.**
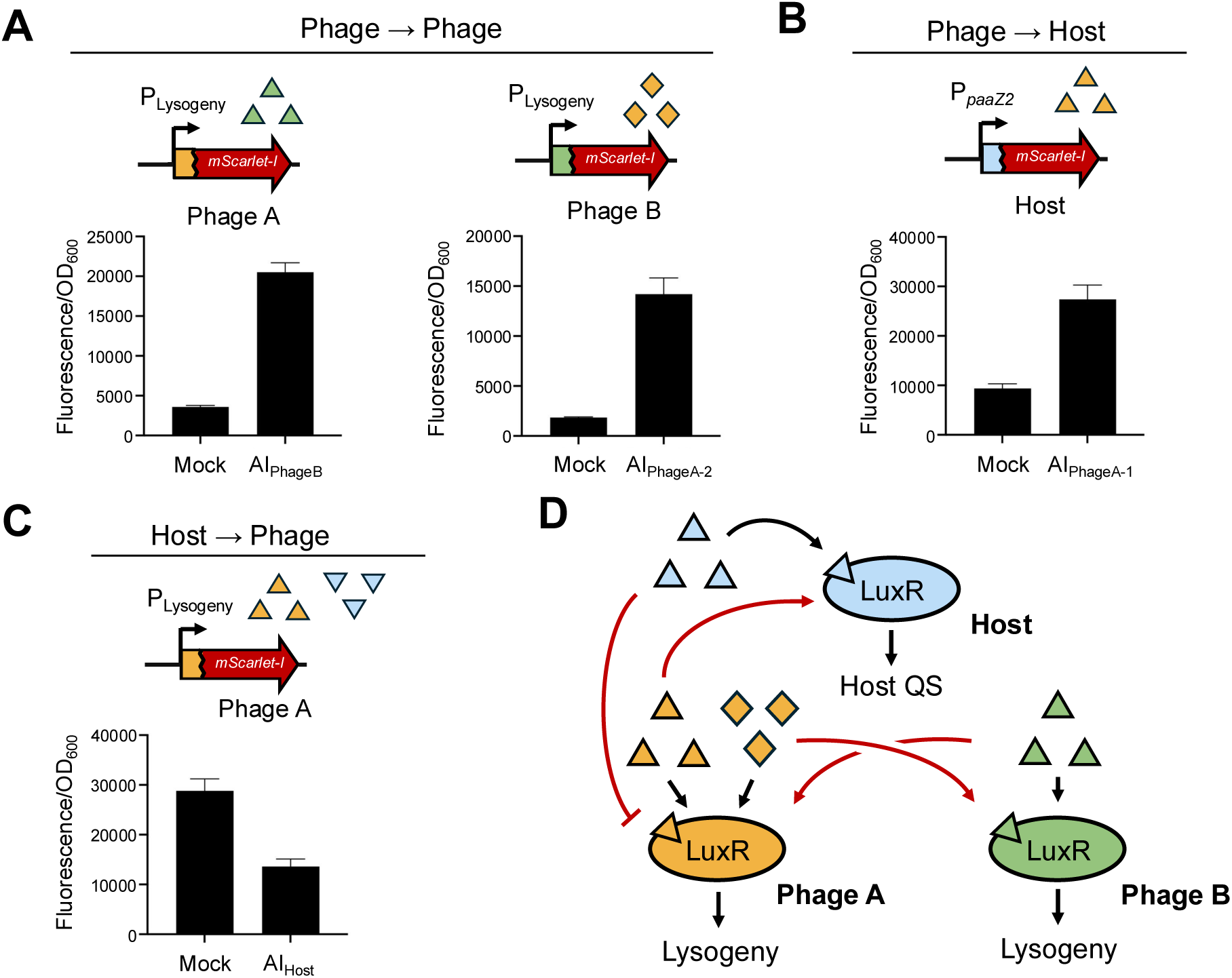
Phage and host autoinducers cross-regulate one another’s QS systems. Shown are schematics of the reporter construct and autoinducer supplementation used in each graph. (**A**) Phage-phage QS cross-regulation. Activity of Phage A (left) and Phage B (right) P_Lysogeny_ reporters in the Δ*3luxI* Δ*3luxR P. inhibens* T5^T^ strain. Each reporter construct contained P_Lysogeny_–*mScarlet-I-xre-luxR* but lacked the *luxI* gene. (**B**) Phage-to-host cross-regulation. Activity of the host QS reporter P*_paaZ2_*-*mScarlet-I* in the Δ*prophage A* Δ*prophage B* Δ*pgaI* strain. (**C**) Host-to-phage cross-regulation. Activity of the Phage A P_Lysogeny_–*mScarlet-I-xre-luxR-luxI* reporter in the Δ*prophage A* Δ*prophage B* Δ*pgaI* strain. In A-C, cultures were treated with 10 µM of the indicated autoinducer or DMSO (Mock). (**D**) Model depicting cross-regulatory interactions between the *P. inhibens* T5^T^ QS systems. Blue triangles represent AI_Host_. Orange triangles and diamonds represent the Phage A autoinducers AI_PhageA-1_ and AI_PhageA-2_, respectively. Green triangles represent the Phage B autoinducer AI_PhageB_. Black arrows indicate endogenous activation. Red arrows indicate cross-regulatory interactions. (A-C) mScarlet-I fluorescence was normalized to OD_600_. Data represent the means ± SDs of four biological replicates.

#### Phage to host cross-regulation

We investigated whether the prophage-produced autoinducers can be detected by the host PgaR receptor. We used the reporter construct carrying the P*_paaZ2_*–*mScarlet-I* fusion in a Δ*pgaI* host. AI_PhageA-1_ stimulated expression (EC_50_ = 4.2 μM) whereas neither AI_PhageA-2_ nor AI_PhageB_ showed activity (Fig. 3B, Fig. S2). Thus, one of the Phage A autoinducers can activate the *P. inhibens* T5^T^ host QS pathway.

#### Host to phage cross-regulation

To complete the analysis of possible cross-regulation, we examined whether the host autoinducer influences the prophage QS circuits. AI_Host_ did not activate either prophage reporter (Fig. S2A). We therefore tested whether AI_Host_ inhibits prophage QS. To do so, we measured the effect of exogenous AI_Host_ on our phage P_Lysogeny_ reporter constructs carrying the full prophage *xre–luxR–luxI* operons. In these strains, the prophage autoinducers produced from the constructs induce expression from their cognate P_Lysogeny_ promoters, allowing us to assess the effect of exogenously supplied AI_Host_. AI_Host_ suppressed Phage A P_Lysogeny_ expression (Fig. 3C) but had no effect on P_Lysogeny_ from Phage B (Fig. S2C).

Above, by relying on exogenously supplied autoinducers, we demonstrate that the three QS systems exhibit positive and negative cross-regulation (Fig. 3D). Our next goal was to test whether cross-regulation occurs under native conditions, i.e., with autoinducers produced by the bacteria/prophages themselves. Accordingly, we introduced the Phage A or Phage B Δ*luxI* reporter constructs into a *P. inhibens* strain carrying or lacking the non-cognate prophage. As a result, reporter activation can only occur in response to autoinducer produced by the non-cognate resident prophage. Both reporters exhibited decreased expression in the absence of the non-cognate prophage (Fig. S3A-B). These results indicate that each prophage naturally produces sufficient autoinducer to cross-activate the other’s LuxR receptor.

Using a similar strategy, we also tested whether Phage A can cross-activate the host QS system *in vivo*. Indeed, Phage A autoinducer produced by the chromosomally-integrated Phage A QS module activated expression from the host reporter construct (Fig. S3C). Moreover, as mentioned above, in *P. inhibens* T5^T^, the host QS system controls the production of the pigmented antibiotic called TDA. This pigment confers a brown color to the bacterial culture and limits the maximum cell density achieved because it is toxic. Deletion of the host autoinducer synthase (*pgaI*) eliminates pigment production and alleviates growth suppression^15,18,20^. To measure *in vivo* cross-regulation, we integrated the Phage A *xre-luxR-luxI* QS module into an ectopic locus on the chromosome of the Δ3*luxI* Δ3*luxR* strain. We introduced *pgaR* under its native promoter on a plasmid into both the parental Δ3*luxI* Δ3*luxR* strain and the derivative carrying the integrated Phage A QS module. We compared TDA pigment production and the final OD_600_ values of the two strains. Pigment production and growth suppression occurred in the strain harboring the Phage A QS module but not the parental Δ3*luxI* Δ3*luxR* strain (Fig. S3D). These results indicate that the Phage A autoinducer can substitute for the host autoinducer *in vivo*.

The prevalence of cross-regulation in the *P. inhibens* T5^T^ system raises the possibility that the QS receptors respond promiscuously to many related autoinducers. To explore whether or not this is the case, we tested each of our QS reporter constructs lacking *luxI* genes against a panel of purified HSLs varying in acyl chain length (C4-C12) and oxidation state (unmodified, 3-OH, and 3-O). The Phage A reporter responded to the non-cognate HSLs C6-HSL, C8-HSL, and 3O-C8-HSL. By contrast, the Phage B reporter did not respond to any non-cognate HSL in our panel. The host reporter responded to its cognate autoinducer 3OH-C10-HSL and the non-cognate HSLs 3OH-C8-HSL, 3O-C10-HSL, and 3OH-C12-HSL (Fig. S4). These results indicate that the QS receptors display relatively narrow ligand specificities, with the Phage B LuxR exhibiting particularly strict selectivity.

### The host PgaR receptor together with its autoinducer represses Phage A QS

Figure 3C shows that AI_Host_ suppresses the Phage A QS response. Two mechanisms could explain this result: (1) the host autoinducer could function through PgaR to represses Phage A P_Lysogeny_ or (2) the host autoinducer could inhibit the Phage A LuxR directly. To distinguish between these possibilities, we introduced the Phage A reporter carrying the full *xre–luxR–luxI* operon downstream of P_Lysogeny_ into two *P. inhibens* T5^T^ strains: the Δ3*luxI* Δ3*luxR* strain and a derivative carrying the *pgaR* receptor gene. We supplied both strains with AI_Host_ and monitored reporter activity. AI_Host_ had no effect in the strain lacking *pgaR*. By contrast, AI_Host_ strongly reduced mScarlet-I fluorescence emission from Phage A P_Lysogeny_ in the Δ3*luxI* Δ3*luxR* strain carrying *pgaR* (Fig. 4A). Thus, the host suppresses Phage A QS via a PgaR-dependent mechanism.

**Figure 4.**
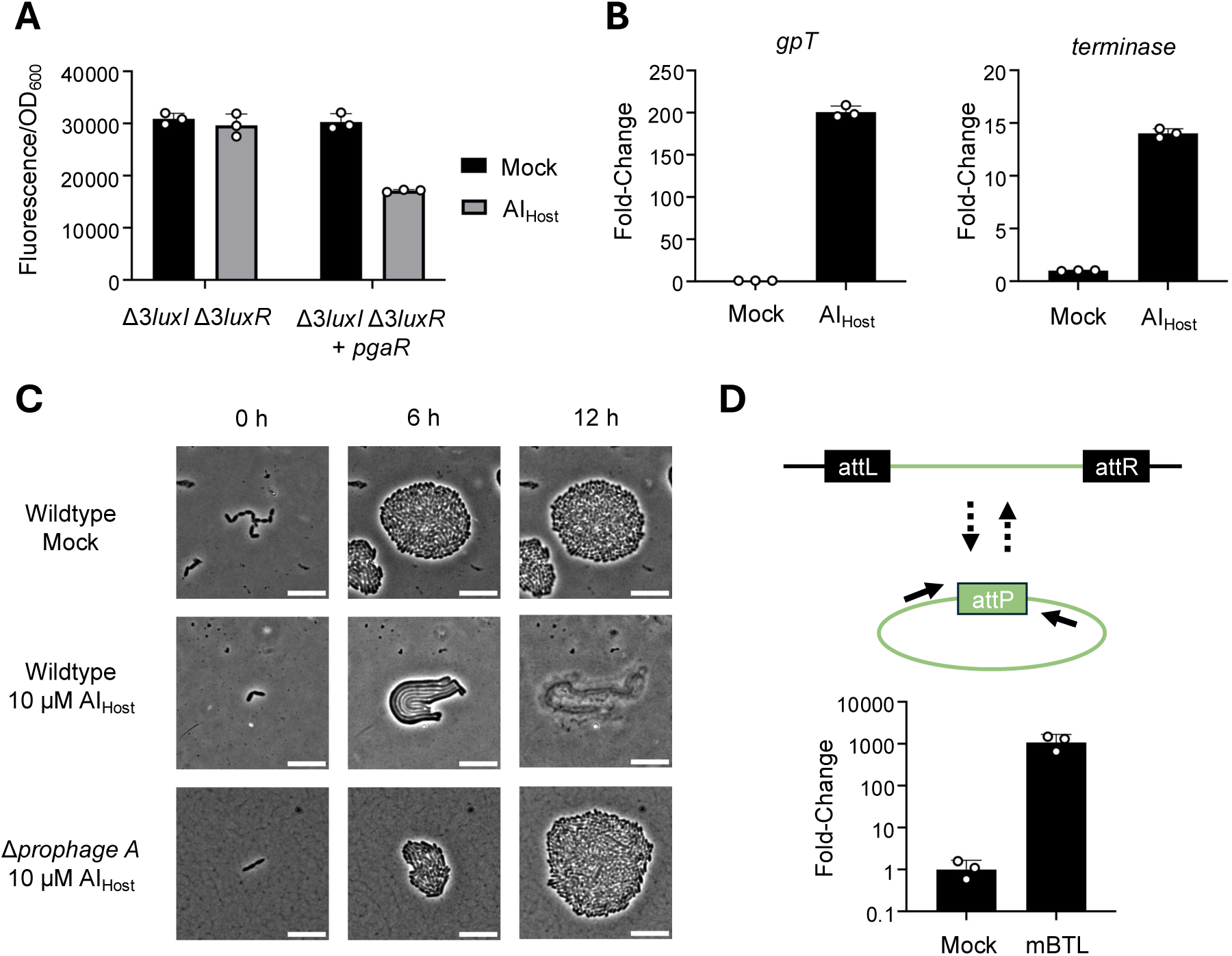
Inhibition of Phage QS triggers lytic induction. (**A**) Phage A P_Lysogeny_–*mScarlet-I-xre-luxR* activity was measured in the Δ*3luxI* Δ*3luxR P. inhibens* T5^T^ strain and in the strain carrying *pgaR* (denoted + *pgaR*). Cultures were administered DMSO (Mock) or 10 μM AI_Host_. (**B**) Expression of representative Phage A structural genes following administration of 10 μM AI_Host_. Transcript levels of the Phage A major head protein (*gpT*) and *terminase* genes were quantified by RT-qPCR in WT *P. inhibens* T5^T^ and are shown as the fold-change relative to mock-treated cultures. (**C**) Represenatative time-lapse microscopy images of the designated *P. inhibens* T5^T^ strains at the times following the indicated treatment. Each treatment was independently replicated three times; in each replicate, approximately 10 microcolonies were imaged over time. Scale bar, 10 μm. (**D**) (Top) Schematic of the assay used to monitor Prophage B excision to generate a circular genome containing the *attP* junction. Primer binding sites are indicated as solid arrows. (Bottom) qPCR quantitation of the Phage B circular *attP* junction following treatment of WT *P. inhibens* T5^T^ with DMSO (Mock) or with 100 μM mBTL for 5 h. Data are reported as fold-change relative to the Mock treatement. (A, B, D) Data represent means ± SDs of three biological replicates.

Unlike Phage A, the Phage B P_Lysogeny_ promoter does not respond to AI_Host_ (Fig. S2C). We therefore required an alternative method to modulate Phage B QS. In earlier work, we developed the synthetic QS antagonist mBTL^21^. Here, we show that mBTL inhibits the Phage B LuxR (Fig. S5A). To determine whether inhibition requires host PgaR, we introduced the Phage B reporter into the Δ3*luxI* Δ3*luxR P. inhibens* strain or its derivative carrying host *pgaR*. mBTL inhibited reporter expression in both strains (Fig. S5A). Thus, mBTL does not require the host PgaR receptor to repress QS in Phage B.

### Inhibition of phage QS triggers prophage lytic induction

Above, we showed that the *xre–luxR–luxI* operons of both prophages activate transcription from their cognate P_Lysogeny_ promoters. In both cases, if P_Lysogeny_ maintains the lysogenic state, then inhibition should trigger P_Lysis_ expression and drive lytic induction. AI_Host_ and mBTL provide the means to inhibit, respectively, the Phage A and Phage B P_Lysogeny_ promoters, allowing us to test this prediction.

We first assessed the consequence of inhibition of Phage A QS. Following treatment with AI_Host_, induction of the Phage A structural genes, including those encoding the major head protein (*gpT*) and the terminase occurred (Fig. 4B). Companion time-lapse microscopy showed a striking host response. Within 3 h of treatment, the *P. inhibens* T5^T^ cells ceased dividing, then began elongating, before lysing at 8 h (Fig. 4C). Consistent with our above results, AI_Host_ driven lysis did not occur in *P. inhibens* T5^T^ cured of Phage A (Fig. 4C). These results show that growth arrest and lysis are due to activation of prophage activity, not toxicity of the AI_Host_ compound. Consistent with these data, DNaseI protection assays show Phage A produces virions (Fig. S5B).

Because apparently no naturally produced inhibitor of Phage B P_Lysogeny_ exists in our system, we cannot perform experiemnts analogous to those we show for Phage A in Figure 4C. Nonetheless, we can use mBTL to assess Phage B lytic induction. Unlike Phage A, Phage B contains recognizable chromosomal attachment sites that enable us to monitor prophage excision. Following mBTL treatment, we detected the circularized Phage B genome and the restored *attP* site (Fig. 4D). Thus, inhibition of Phage B QS triggers excision from the host chromosome and release of DNaseI protected phage particles (Fig. S5B). Together, these results show that inhibition of QS induced both prophage lytic cascades.

### Host:prophage autoinducer ratios govern Prophage A lysogeny-lysis transitions

Because the Prophage A and host autoinducers exert opposing effects on Prophage A P_Lysogeny_, we hypothesized that P_Lysogeny_ could respond to their relative abundances (Fig. 5A). If so, Phage A could tie its lysis-lysogeny decision making to the proportion of uninfected *P. inhibens* T5^T^ hosts in its environment. In a population dominated by hosts lysogenized by Prophage A, AI_PhageA-1_ should accumulate relative to AI_Host_ and enhance expression from Phage A P_Lysogeny_. By contrast, in a population of primarily uninfected hosts, AI_Host_ should predominate, suppress expression from P_Lysogeny_, and trigger Prophage A induction.

**Figure 5.**
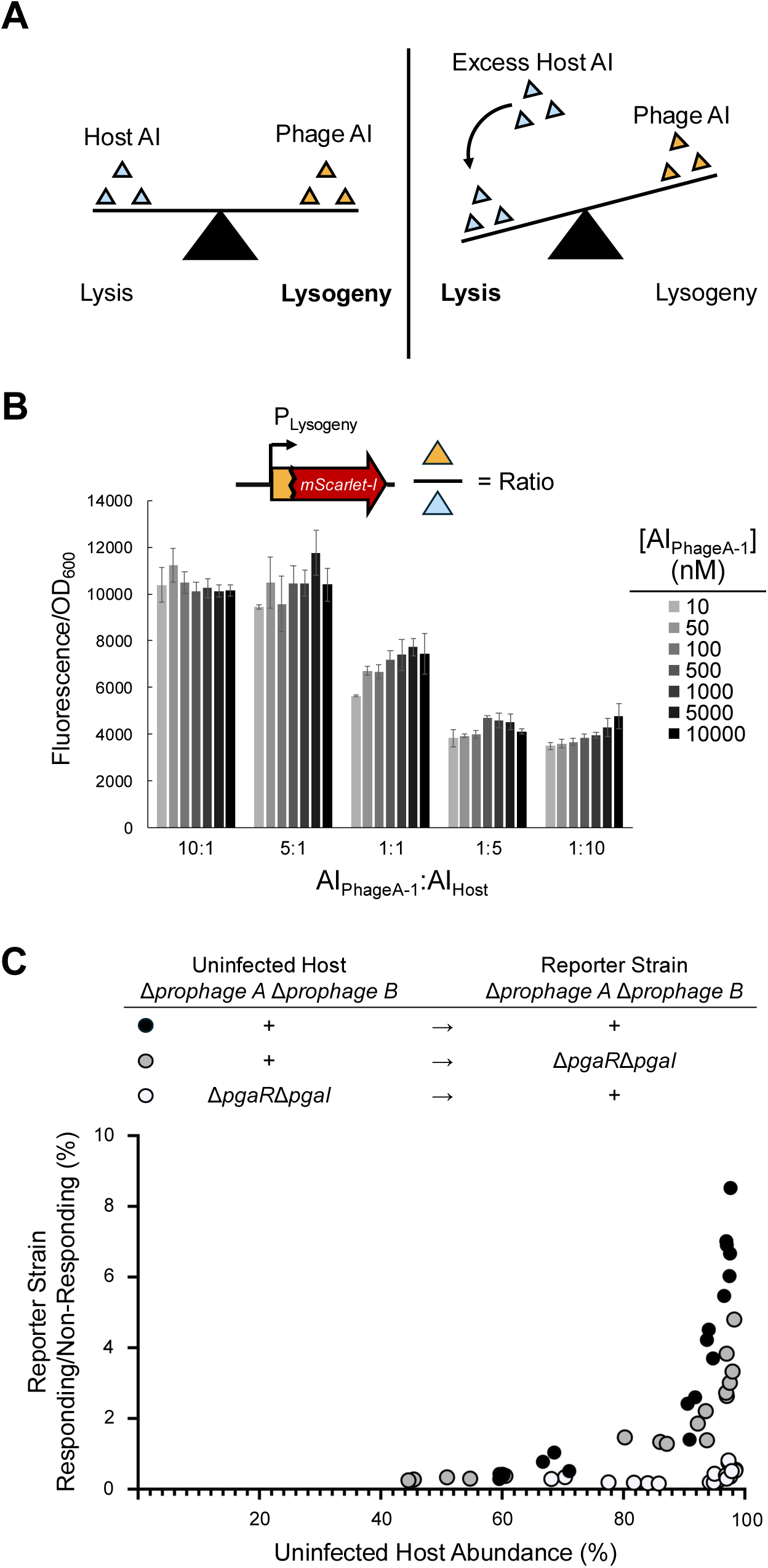
Opposing host and phage autoinducer inputs enable Prophage A to detect AI ratios. (**A**) Model of host and phage autoinducer integration by prophage A. See text for details. (**B**) Activity of the Phage A P_Lysogeny_–*mScarlet-I-xre-luxR* reporter in the Δ*prophage A* Δ*prophage B* Δ*pgaI P. inhibens* T5^T^ strain. Cultures were supplemented with the indicated concentrations of AI_PhageA-1_ and appropriate corresponding amounts of AI_Host_ to generate the ratios designated on the x-axis. mScarlet-I fluorescence was normalized to OD_600_. Data represent the means ± SDs of four biological replicates. (**C**) Single-cell flow cytometry analyses of Prophage A reporter activity as a function of Uninfected Host abundance. DAPI was added prior to flow cytometry to distinguish cells from debris. The fraction of Uninfected Host cells in each population was calculated from the ratio of sfGFP^−^ (Uninfected Host) to sfGFP^+^ (Reporter Strain) cells. Reporter Strain cells were classified as responding or non-responding based on their positions within predefined fluorescence gates, and the percentage of responding Reporter Strain cells was calculated as the number of Reporter Strain cells in the responding gate divided by the total number of Reporter Strain cells. Black circles indicate mixtures of the Reporter Strain and the Uninfected Host. Gray circles indicate mixtures of the Δ*pgaR* Δ*pgaI* Reporter Strain and the Uninfected Host. White circles indicate mixtures of the Reporter Strain and the Δ*pgaR* Δ*pgaI* Uninfected Host. Each data point represents an independent co-culture analyzed by flow cytometry and includes at least 10,000 viable Reporter Strain cells.

To test this idea, we measured Phage A P_Lysogeny_ expression across a range of AI_Host_ and AI_PhageA-1_ concentrations while holding the ratio of the two autoinducers constant. We used the Phage A P_Lysogeny_ reporter construct lacking *luxI* in our derivative of the *P. inhibens* Δ3*luxI* Δ3*luxR* strain carrying the host *pgaR* gene. This strain, therefore, does not produce, but can respond to both AI_Host_ and AI_PhageA-1_. Figure 5B shows that P_Lysogeny_ expression converged to similar values across a 1,000-fold range of absolute autoinducer concentrations when AI_Host_:AI_PhageA-1_ remained constant. This result indicates that the relative abundances of the AI_Host_ and the AI_PhageA-1_ control P_Lysogeny_ expression.

To determine whether AI_Host_ and AI_PhageA-1_ autoinducer ratios control the Phage A lysis-lysogeny decision *in vivo*, we used microscopy to follow the fates of WT *P. inhibens* T5^T^ cells while synthetically altering the balance between AI_Host_ and AI_PhageA-1_. In the absence of added autoinducers, *P. inhibens* grew normally and exhibited its characteristic oval morphology (Fig. S5C). As in Fig. 4C, addition of 10 μM AI_Host_ triggered Phage A-directed cell lysis, consistent with inhibition of Phage A QS and activation of its lytic cascade (Fig. S5C). If the Phage A regulatory switch indeed depends on the relative amounts of host and phage autoinducers, we reasoned that increasing the concentration of the Phage A autoinducer should counteract the effect of the added AI_Host_. Consistent with this model, administration of 10 μM AI_PhageA-1_ and 10 μM AI_Host_ restored normal *P. inhibens* T5^T^ growth and prevented cell lysis (Fig. S5C). Thus, the relative abundances of AI_Host_ and AI_PhageA-1_, not their absolute concentrations, control the Phage A lysis-lysogeny transition.

We also explored whether Phage A detection of both its own and its host’s autoinducer enables it to restrict its lysogeny-to-lysis transition to periods when the concentration of host autoinducer far exceeds the concentration of its own autoinducer. Presumably, in natural habitats, this scenario would occur when most of the host cells in the vicinity are not already lysogenized by Phage A. Lysing a current host only in environments containing primarily uninfected host cells could maximize Phage A transmission as Phage A progeny would not suffer losses from superinfection exclusion mechanisms.

To test this notion, we constructed a dual-fluorescence reporter plasmid in which the Phage A P_Lysogeny_ promoter drives expression of *mScarlet-I* and the PhageA *xre–luxR–luxI* operon. This plasmid also expresses *sfGFP* from a different, constitutive promoter, providing both an internal fluorescence normalization control for plasmid copy-number and cell-tracking marker. We introduced this plasmid into a *P. inhibens* T5^T^ Δ*prophage A* Δ*prophage B* strain, hereafter referred to as the Reporter Strain. We mixed this strain at defined ratios with a second Δ*prophage A* Δ*prophage B* strain carrying an empty vector, referred to as the Uninfected Host. We used single-cell flow cytometry to measure the number of cells of each strain and the expression of *mScarlet-*I in the Reporter Strain. Because only the Reporter Strain can produce AI_PhageA-1_, whereas both strains produce AI_Host_, increasing the fraction of Uninfected Host cells increases the AI_Host_:AI_PhageA-1_ ratio. We used the ratio of sfGFP-positive to sfGFP-negative events to determine the ratio of Reporter Stain cells to Uninfected Host cells in each co-culture. We also classified Reporter Strain cells as responding or non-responding to AI_Host_ based on predefined mScarlet-I fluorescent gates (see Methods).

As the fraction of Uninfected Host cells increased, a larger fraction of Reporter Strain cells repressed P_Lysogeny_ expression, i.e., transitioned from the non-responding to the responding gate (Fig. 5C, black symbols). This effect required host QS as it was diminished when the Uninfected Host or Reporter Strain lacked PgaR and PgaI (Fig. 5C, gray and white symbols, respectively). Together, these results indicate that Phage A exploits the opposing effects that AI_Host_ and AI_PhageA-1_ have on its P_Lysogeny_ promoter to infer the relative abundances of infected and uninfected hosts in the population. This information then directs when Phage A makes lysogeny-lysis transitions. This capability enables Phage A to preferentially induce its lytic cascade under conditions when cells in the vicinal community are primarily uninfected by Phage A.

### QS regulatory modules are widespread among mobile genetic elements in Alphaproteobacteria

Our above findings show how two unrelated prophages employ distinct QS circuits to regulate their lysis-lysogeny decisions. Their co-residence in *P. inhibens* T5^T^ therefore captures two independent instances of this evolutionary solution within a single genome. Yet, this arrangement does not provide information on broader evolutionary questions concerning QS-mediated phage-phage and phage-host competition. To explore the scale of these interactions, we screened Alphaproteobacteria, the class to which *P. inhibens* belongs, for genomes containing two or more *luxI* genes. Our presumption is that most genomes would harbor a single host-associated *luxI* gene, so the presence of additional copies of *luxI* could indicate the presence of a phage. This approach identified 1,338 *luxI* genes distributed across 528 genomes containing multiple *luxI* genes. Using an 80% amino acid identity cutoff produced 94 clusters containing at least three members. To identify potential phage association, we first identified *luxI* genes displaying hallmarks of horizontal transfer. To do this, we constructed a core genome phylogenetic tree for the Alphaproteobacteria, onto which we mapped the distribution of the 94 LuxI protein clusters (Fig. S6). We detected horizontally transferred *luxI* genes in three orders: the Rhodobacterales, which encompasses the Roseobacter clade and *Phaeobacter* spp., the Sphingomonadales, and the Hyphomicrobiales. A substantial fraction of the Hyphomicrobiales hits corresponded to *traI/R* and *rhiABC* genes associated with known conjugative plasmids. These instances were excluded from further analysis.

Regarding the remaining *luxI* genes, we extracted 100 kb genomic regions flanking each locus and subjected them to analysis with VIBRANT to identify putative prophage regions^22^. We detected eight horizontally transferred *luxI* clusters associated with predicted prophages and that were located in *xre-luxR-luxI* operons. These modules were confined exclusively to the Roseobacter clade (Fig. 6A, Fig. S6). Analogous to the broader *luxI* distribution, the arrangement of these clusters does not reflect the phylogeny of the Roseobacter genomes, consistent with their acquisition through horizontal transfer. An additional four genomes harbored QS systems not initially predicted to be prophage associated. However, manual inspection revealed that in each case, the *luxI* gene resided within degraded prophage remnants (Fig. S7).

**Figure 6.**
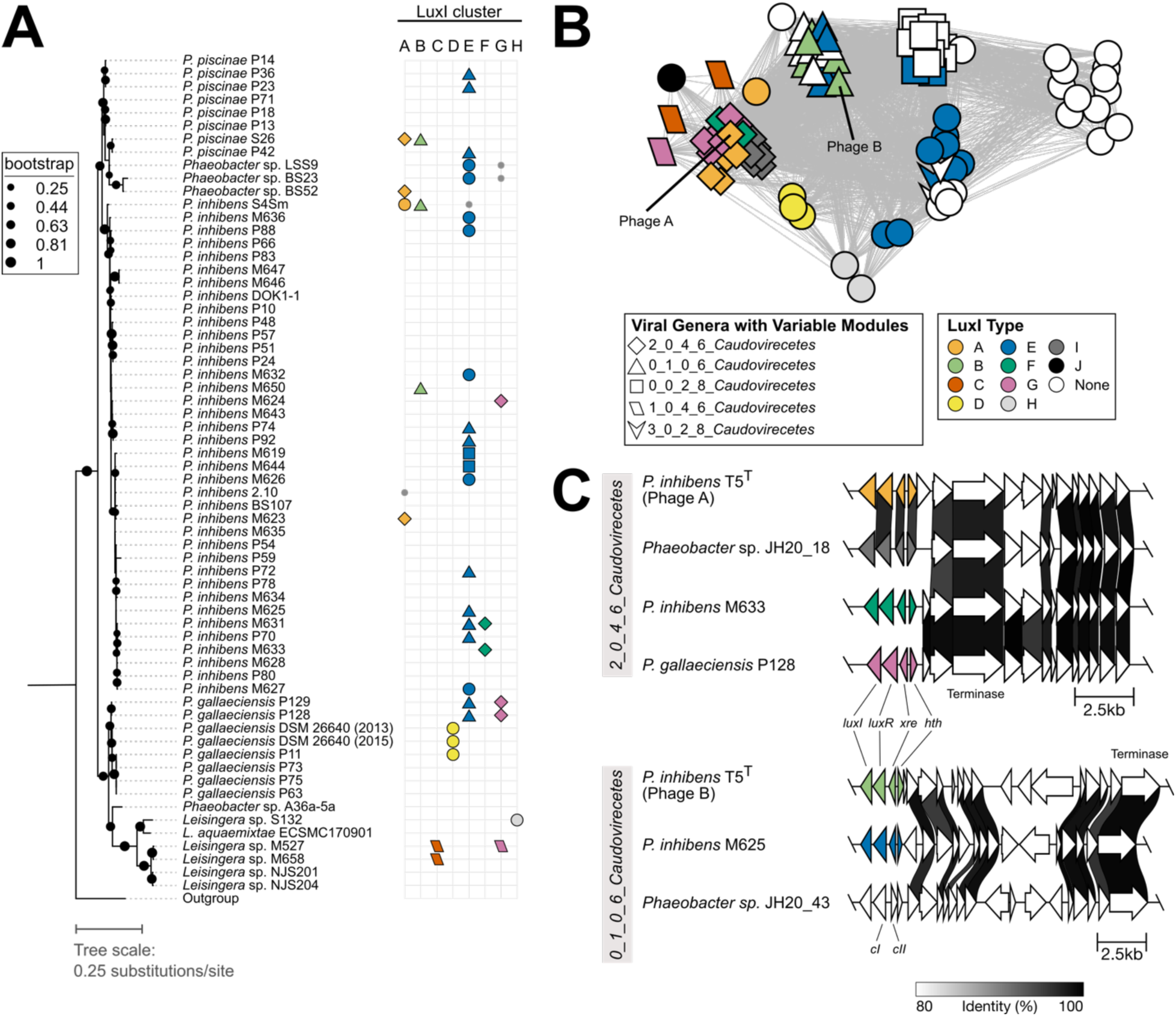
Variable phage-borne *luxI/R* modules are horizontally exchanged between *Phaeobacter* strains and their phages. (**A**) Core-genome phylogenetic tree of Roseobacter genomes with their corresponding LuxI proteins. Bootstrap support for each node is shown on the tree. Scale represents substitutions per amino acid site. LuxI proteins not predicted to be in a phage are shown as small gray points. LuxI proteins predicted to be harbored by a phage are shown as large symbols. Symbol colors and shapes correspond to the LuxI type and the predicted phage genus (shown in panel B), respectively. (**B**) Taxonomic network of *Phaeobacter* phages genomes. Nodes represent individual phages. Node colors represent the LuxI type harbored by each phage. For viral clusters with multiple LuxI types, node shapes represent vConTACT3-predicted genera. Edge weights were calculated by vConTACT3. (**C**) Genome synteny of *xre-luxR-luxI* operons and the neighboring sequences from select phages in viral genera containing Phage A and Phage B. Host species and strain are provided on the left. Arrows represent genes. *xre-luxR-luxI* operon genes are colored according to their assigned LuxI type. Gene homologs in neighboring sequences are connected by shaded links representing the percent identity between the amino acid sequences of the proteins encoded by the homologous genes. The absence of a link indicates less than 80% amino acid identity between proteins encoded by neighboring genes or the absence of a homolog in the neighbor.

Via an opposite strategy, we assessed the prevalence of QS systems among Roseobacter prophages. Using the identified prophage loci as database queries, we expanded our prophage dataset to 93 unique putative prophages and identified two additional *luxI* clusters that were missed in our initial analysis (Figure 6A,B; Fig. S8-S11). Of the 93 unique, predicted prophages, 53 harbored *luxI* QS systems, indicating that phage-encoded QS is a common feature among this group. To characterize the evolutionary relationships among these prophages and the distributions of their QS systems, we analyzed the predicted prophage genomes using vConTACT3, which employs a gene-sharing network approach to cluster related phages at the genus level (Fig. 6B)^23^. Excluding singletons, the 93 prophages resolved into 10 high-confidence novel phage genera with Phage A and Phage B studied here segregating into distinct genera, consistent with their divergent genomic architectures.

Several notable patterns emerged from this analysis. Within both the Phage A and Phage B genera, there exist distinct QS genes among closely related phages, implying that *luxI/R* modules undergo genetic exchange as discrete units within phage genera. This modularity is illustrated in Figure 5C (top), which shows four divergent *luxI/R* QS modules being shared across phage genomes with strong conservation across the remainder of the genome (Supplementary Table 1). Intriguingly, certain QS modules appear restricted to a particular phage genus, while others have transferred across genera, suggesting that these elements vary considerably in their mobilities. In some instances, the QS module has been replaced entirely by a non-QS regulatory region, as shown within the Phage B genus (Fig. 6C, bottom). Consistent with roles in governing lysis-lysogeny decisions, these alternative modules encode genes associated with DNA damage sensing, including cI repressor genes, and genes encoding proteins containing BRCA1 C-Terminus (BRCT) domains. Together, these data show how prophage QS gene modules behave as exchangeable regulatory cassettes, interchangeable with other QS modules and with modules encoding completely different regulators.

## Discussion

In natural microbial communities, organisms release autoinducers into shared chemical environments. In doing so, they may leak information about their states and abundances to neighboring organisms with complementary or competing evolutionary interests. The public nature of these signaling environments can influence who benefits from and who is harmed by the exchanged information and can impinge on how effectively organisms preserve or corrupt signaling fidelity. Here, we employed *P. inhibens* T5^T^ as a model system to investigate how phages exploit public chemical information to gauge population composition, manipulate competitors, and shape lysis-lysogeny decisions.

*P. inhibens* T5^T^ harbors three unique QS systems: one encoded by the host and one encoded by each of two chromosomally-integrated prophages, that we named Phage A and Phage B^14^. We demonstrated that each system responds to its cognate autoinducer. Moreover, the two prophages cross-activate one another’s QS systems. Phage A also engages in a bidirectional interaction with the *P. inhibens* T5^T^ host: one of the Phage A autoinducers activates host QS, while the *P. inhibens* T5^T^ host autoinducer represses the Phage A QS system.

In this system, the phage QS modules tie autoinducer concentration to their lysis-lysogeny transitions through a genetic switch. Specifically, in each case, the liganded LuxR receptor drives the expression of an operon encoding its cognate autoinducer synthase (*luxI*) and the *luxR* gene itself. This arrangement launches and reinforces lysogeny through a classic QS autoinduction mechanism. Because only lysogens produce autoinducer, its concentration reflects the abundance of prophages in the population. This regulatory logic closely parallels that of the *Bacillus* arbitrium system in which SPbeta phages use a diffusible peptide as a proxy for lysogen density^12,13^.

In isolation, each *P. inhibens* T5^T^ phage can robustly regulate its lysis-lysogeny transitions using the information encoded in the cognate autoinducer. However, this architecture suffers from an inherent vulnerability: any molecule capable of activating the phage LuxR will drive lysogeny, irrespective of the organism that produced it. Indeed, in *P. inhibens* T5^T^, the shared chemical environment permits autoinducers produced by one organism, host or phage, to influence the behavior of another. This coupling creates opportunities for information extraction and cross-organism control that are not possible with an individual QS system operating alone.

The interconnectedness of the *P. inhibens* T5^T^ QS circuits allows autoinducers to act as both QS information carriers and as tools for QS manipulation. Because both prophages produce autoinducers that cross-activate the other’s QS system, they bias their competitor toward lysogeny. This strategy may allow each prophage to suppress competitor virion production and thereby alter the outcome of phage-phage competition at both the single cell and population levels. A parallel phenomenon was recently described in arbitrium systems, in which peptides produced by one phage induced lysogeny in another phage^24,25^. In both cases, the prophage autoinducers play dual roles: they function as messengers of prophage population density and also as mediators of inter-phage competition. Additionally, in the *P. inhibens* T5^T^ system, one of Phage A’s autoinducers also activates the host QS system. Host QS, in turn, activates the production of TDA, an antibiotic known to kill competitor bacteria^26,27,28^. One potential consequence of Phage A stimulating population-wide TDA production is to allow Phage A to “clear the decks” of competitor bacteria. This capability presumably allows Phage A to engineer a bacterial community dominated by *P. inhibens* T5^T^. When host lysis occurs, elimination of non-*P. inhibens* T5^T^ bacteria in the vicinity would increase the probability that progeny virions exclusively encounter other *P. inhibens* T5^T^ cells, maximizing phage transmission.

The shared *P. inhibens* T5^T^ cellular and chemical environments also create opportunities for QS “eavesdropping”. In our system, Phage A surveils host QS through PgaR-mediated repression of its P_Lysogeny_ promoter. This feature allows Phage A to measure the concentrations of both AI_Host_ and AI_PhageA-1_. Because the two autoinducers exert opposing effects on Phage A P_Lysogeny_, the promoter effectively computes their ratio. The transcriptional response of Phage A P_Lysogeny_ therefore depends not on the absolute concentration of either autoinducer but, rather, on their relative abundances. Consistent with this assertion, we demonstrated that Phage A P_Lysogeny_ expression converges to one level across a wide range of absolute host and Phage A autoinducer concentrations when their ratio is held constant. Similarly, altering the ratios of infected and uninfected *P. inhibens* hosts in a population predictably shifts Phage A P_Lysogeny_ expression. When uninfected hosts dominate the population, AI_Host_ accumulates relative to AI_PhageA-1_, suppressing P_Lysogeny_ and committing Phage A to lytic induction. When lysogens dominate the population, the AI_PhageA-1_ to AI_Host_ ratio is high, driving Phage A into lysogeny, and preventing Phage A from releasing virions into a population of already infected hosts. Therefore, the ratiometric autoinducer detection mechanism we have discovered here enables Phage A to perform a community-level census from within a single cell.

Our results suggest that co-resident prophages can use autoinducers as barriers to competition. Deploying such chemical obstacles presumably also shapes which phages can successfully infect already lysogenized hosts. Consider an invading phage entering a host population replete with an autoinducer produced by a resident prophage. Upon infection, the invader’s QS receptor would detect the already present autoinducer and lock in the lysogeny gene expression program before the phage could launch its lytic cascade. This mechanism would allow the resident prophage to prevent the spread of competitors that rely on identical or cross-reactive QS systems. In this model, prophages partition their environments into restricted chemical niches, each defined by the autoinducers produced and detected. Notably in this context, our analysis shows some Roseobacter genomes maintain *luxI/R* modules within otherwise cryptic prophages. We speculate that hosts may exploit these retained QS modules to constrain the local signaling environment and interfere with incoming phages. If so, resident prophages and their hosts may effectively monopolize particular signaling niches, limiting invasion by interlopers that rely on structurally similar autoinducers. Such a scenario would favor co-existence of prophages harboring sufficiently distinct QS systems.

Based on this idea, we predict that competition within a shared chemical environment imposes opposing selective pressures on the *luxI* and *luxR* genes in prophage QS modules. In a polylysogen, phage LuxI synthases benefit from promiscuity. Specifically, autoinducers that cross-activate a competitor’s P_Lysogeny_ promoter suppress rival phage replication. We find a sign of this in *P. inhibens* T5^T^: Phage A produces two structurally distinct autoinducers, one of which selectively activates the Phage B LuxR receptor and the other of which selectively activates the host QS system. LuxR receptors face the opposite pressure. A receptor that responds to a competing organism’s autoinducer becomes vulnerable to manipulation^24,25^. Extreme receptor specificity therefore prevents interference. Consistent with this logic, both the Phage A and Phage B LuxR proteins exhibit display little response to most non-cognate HSLs. The result is an evolutionary wedge: synthases are selected to expand their influence over competing QS circuits, whereas receptors are selected to restrict that input^28^.

The competitive interactions we identify in *P. inhibens* T5^T^ suggest that phages face strong selective pressure to avoid manipulation by co-resident QS systems. Indeed, the patterns we observe across the Roseobacter metagenome suggest phages have three potential escape routes. First, the phages can escape manipulation through diversification of their existing QS modules. Mutations that produce new autoinducers or alter receptor specificity generate new signaling niches insulated from existing competitor QS components^29,30^. Consistent with this idea, the Roseobacter *luxI/R* genes display significant diversity and cluster into different groups. Over time, diversification produces a reservoir of functionally distinct modules that provide the raw material for the second escape strategy: the replacement of one QS module for another^31^. Module replacement could immediately alter a phage’s signaling interactions, allowing it to eliminate competition by organisms occupying the same chemical niche. Indeed, we identified closely related phages harboring distinct QS modules, indicating that *luxI/R* cassettes undergo exchange as discrete units. Finally, phages can avoid manipulation by abandoning QS regulation altogether. In this context, our genomic analyses reveal phages that appear to have exchanged their QS modules for modules encoding cI- or BRCT-based regulators which respond to DNA damage rather than to autoinducers. By adopting an orthogonal regulatory circuit, these phages become insensitive to manipulation through competitor QS signals. Although this strategy sacrifices access to information encoded in autoinducers, it eliminates the vulnerabilities inherent to operating within a shared signaling environment. Together, these patterns suggest that the remarkable diversity of lysis-lysogeny regulatory circuits present across Roseobacter prophages may reflect the cumulative outcomes of selection to evade manipulation within shared chemical environments.

The *P. inhibens* T5^T^ three-way QS interactions illustrate a broader principle in microbial QS: autoinducers are not merely communication cues, but, rather, constituents of shared chemical environments to which multiple organisms can contribute and extract information. The public nature of the chemical signals fashions opportunities for surveillance and manipulation, which in turn, propels selection for innovation in QS systems. We propose that connections between information, competition, and adaptation are likely general features of phage QS systems in naturally occurring microbial communities.

## Supporting information

Notice of HHMI Publication Policy

## Acknowledgements

We are grateful to all members of the Bassler laboratory for insightful discussions. We thank Dr. Julie Valastyan for her help with RT-qPCR experiments. We thank Dr. Christina DeCoste and the Princeton University Flow Cytometry Core Facility for assistance.

## Author Contributions

Outlined the study: M.L.B and B.L.B.

Performed experiments: M.L.B., G.D.D., T.V.P.N., F.J.S., and E.E.S.

Analyzed data: M.L.B., G.D.D., T.V.P.N., F.J.S., E.E.S, and B.L.B.

Interpreted data: M.L.B., G.D.D., T.V.P.N., F.J.S., E.E.S, and B.L.B.

Wrote manuscript: M.L.B. and B.L.B.

## Lead Contact

Requests for further information and reagents may be directed to and will be fulfilled by the lead contact, Bonnie L. Bassler.

## Materials Availability

All plasmids in this study are available from the lead contact upon request.

## Data and Code Availability

Custom code is provided in the supplementary material.

## Supplementary figure Legends

**Figure S1.**
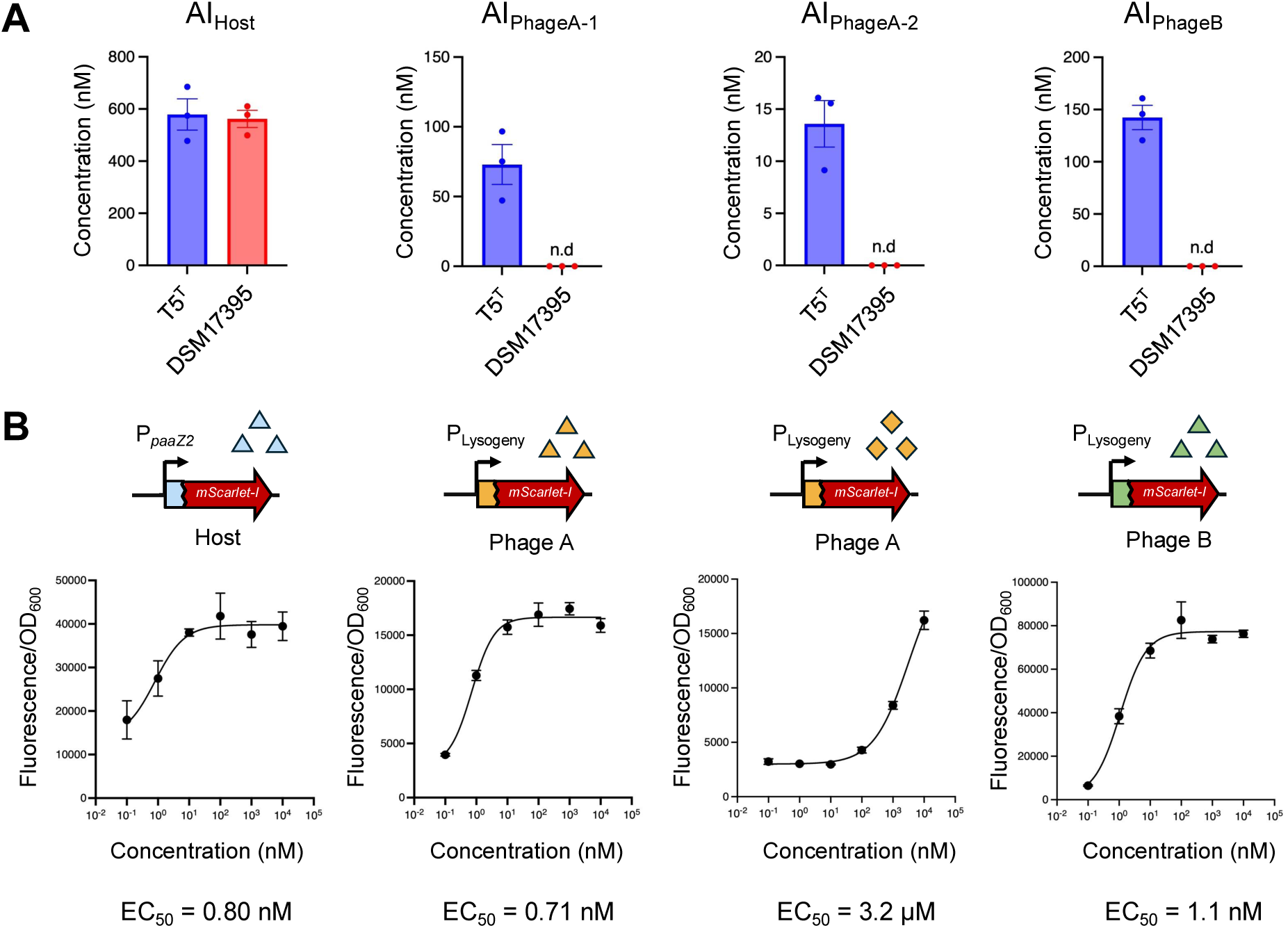
Quantitation of and receptor dose responses to *P. inhibens* T5^T^ HSLs. (**A**) Quantitation of HSLs produced by *P. inhibens* T5^T^ and the related strain DSM17395 that lacks the QS prophages. Data were acquired from stationary phase cultures that had been grown for 16 h at 28°C with shaking at 200 rpm. (**B**) Dose response curves for host and prophage LuxR receptors to their cognate HSLs. The Host curve was generated from P*_paaZ2_*-*mScarlet-I* in the *P. inhibens* Δ*3luxI* Δ*3luxR* strain carrying the *pgaR* gene. The Phage curves employed the respective P_Lysogeny_–*mScarlet-I-xre–luxR* constructs in the *P. inhibens* Δ*3luxI* Δ*3luxR* strain. Strains were incubated with the indicated concentrations of HSLs and fluorescence was normalized to the culture OD_600_. Curves were fit using a four-parameter logistic model and the resulting EC_50_ values are indicated below each graph. Data represent means ± SDs from three (panel A) and four (panel B) biological replicates. n.d., not detected.

**Figure S2.**
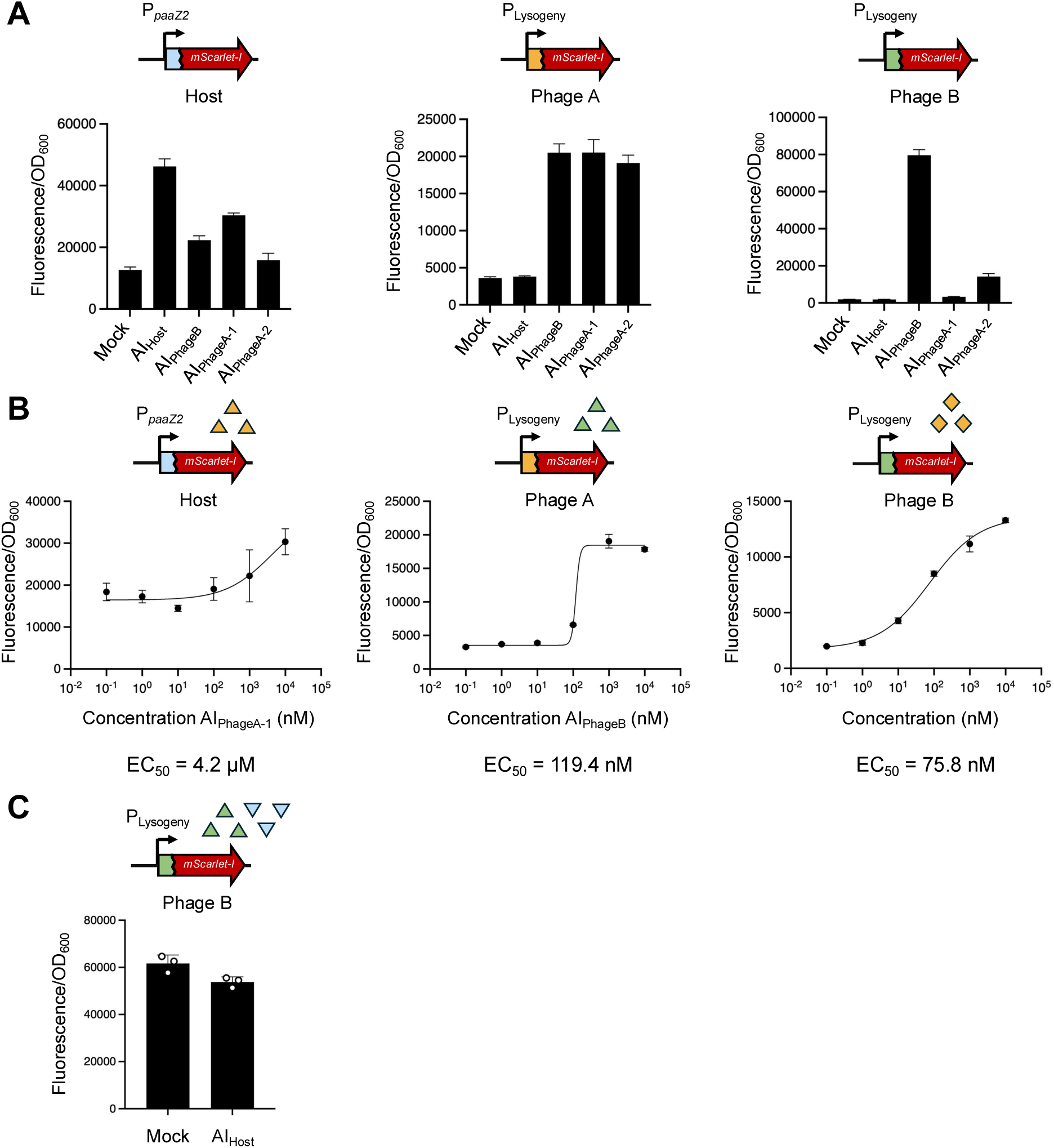
Cross-activation of host and prophage QS circuits. (**A**) Responses of the host and prophage QS reporters to non-cognate HSLs. The Host data were generated from P*_paaZ2_*-*mScarlet-I* in the *P. inhibens* Δ*3luxI* Δ*3luxR* strain carrying the *pgaR* gene. The Phage data were generated from the respective P_Lysogeny_–*mScarlet-I-xre–luxR–luxI* constructs in the *P. inhibens* Δ*3luxI* Δ*3luxR* strain. Strains were grown in the presence of 10 μM of the indicated HSL and fluorescence was normalized to the OD_600_. (**B**) Dose responses of Host and Phage LuxR receptors to non-cognate HSLs. Dose-response curves for the indicated reporter and HSL. Curves were fit using a four-parameter logistic model and the resulting EC_50_ values are indicated below each graph. In both panels, data represent means ± SDs from four biological replicates. (**C**) Activity of the Phage B P_Lysogeny_–*mScarlet-I-xre-luxR-luxI* reporter in the Δ*prophage A* Δ*prophage B* strain. Cultures were treated with DMSO (Mock) or 10 μM AI_Host_. Data represent means ± SDs from three biological replicates.

**Figure S3.**
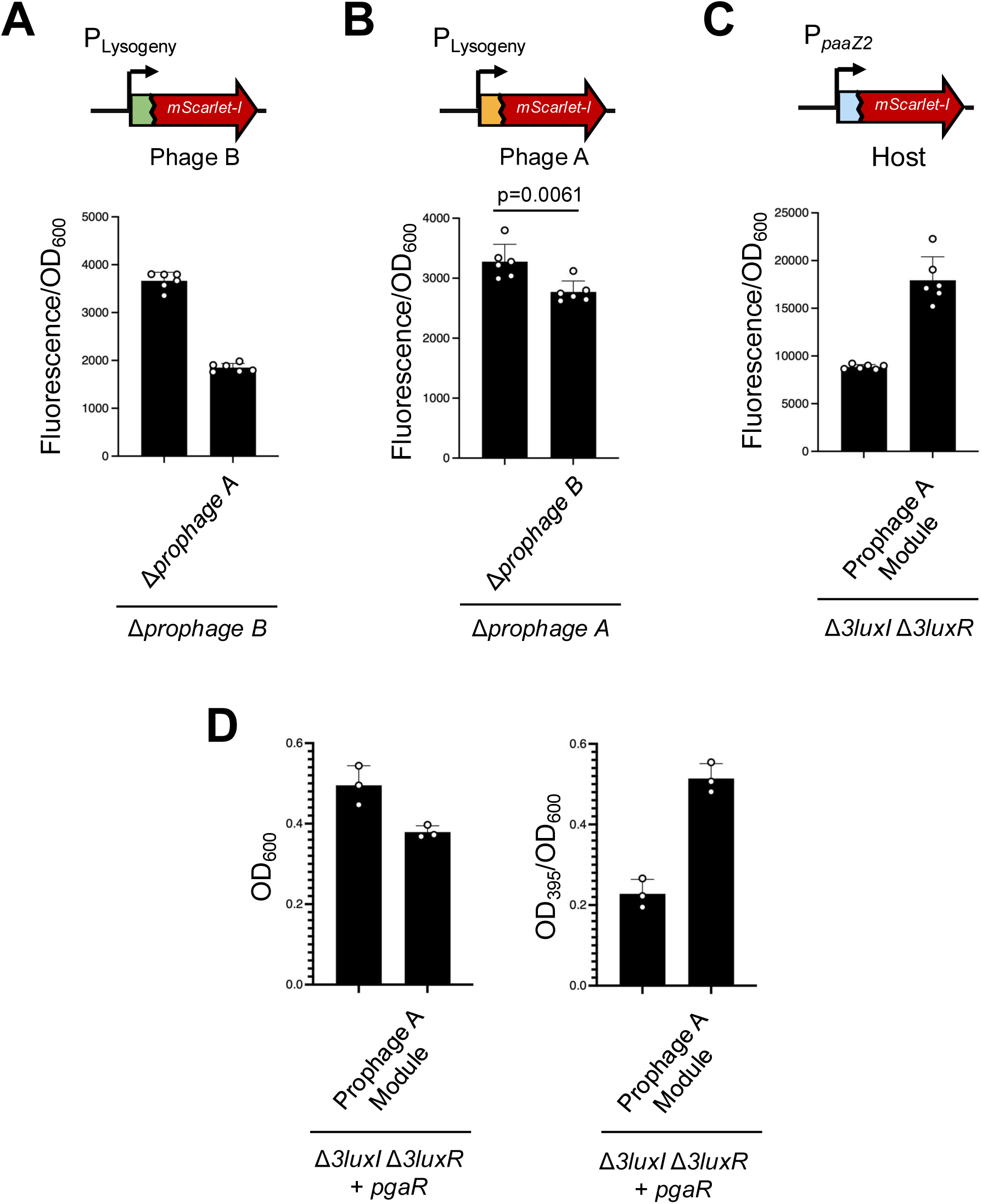
Cross-regulation between host and prophage QS systems *in vivo*. (**A**) *In vivo* cross-activation of the Prophage B QS system by Prophage A. The Prophage B P_Lysogeny_–*mScarlet-I-xre–luxR* reporter was introduced into either the Δ*prophage B* or Δ*prophage A* Δ*prophage B* strains. mScarlet-I fluorescence was normalized to OD_600_. (**B**) *In vivo* cross-activation of the Prophage A QS system by Prophage B. The Prophage A P_Lysogeny_–*mScarlet-I-xre–luxR* reporter was introduced into either the Δ*prophage A* or Δ*prophage A* Δ*prophage B* strains. mScarlet-I fluorescence was normalized to OD_600_. (**C**) Activation of the host QS system by the Prophage A QS module. The host P*_paaZ2_*–*mScarlet-I* reporter was introduced into either the Δ*3luxI* Δ*3luxR* strain or a derivative carrying the phage A QS module integrated into the chromosome. mScarlet-I fluorescence was normalized to OD_600_. (**D**) Phage A-mediated induction of TDA production. A plasmid expressing *pgaR* from its native promoter was introduced into either a Δ*3luxI* Δ*3luxR* strain or a derivative carrying the phage A QS module integrated into the chromosome. Left, final cell culture density (OD_600_). Right, production of the brown TDA-associated pigment quantified as absorbance at 395 nm to OD_600_.

**Figure S4.**
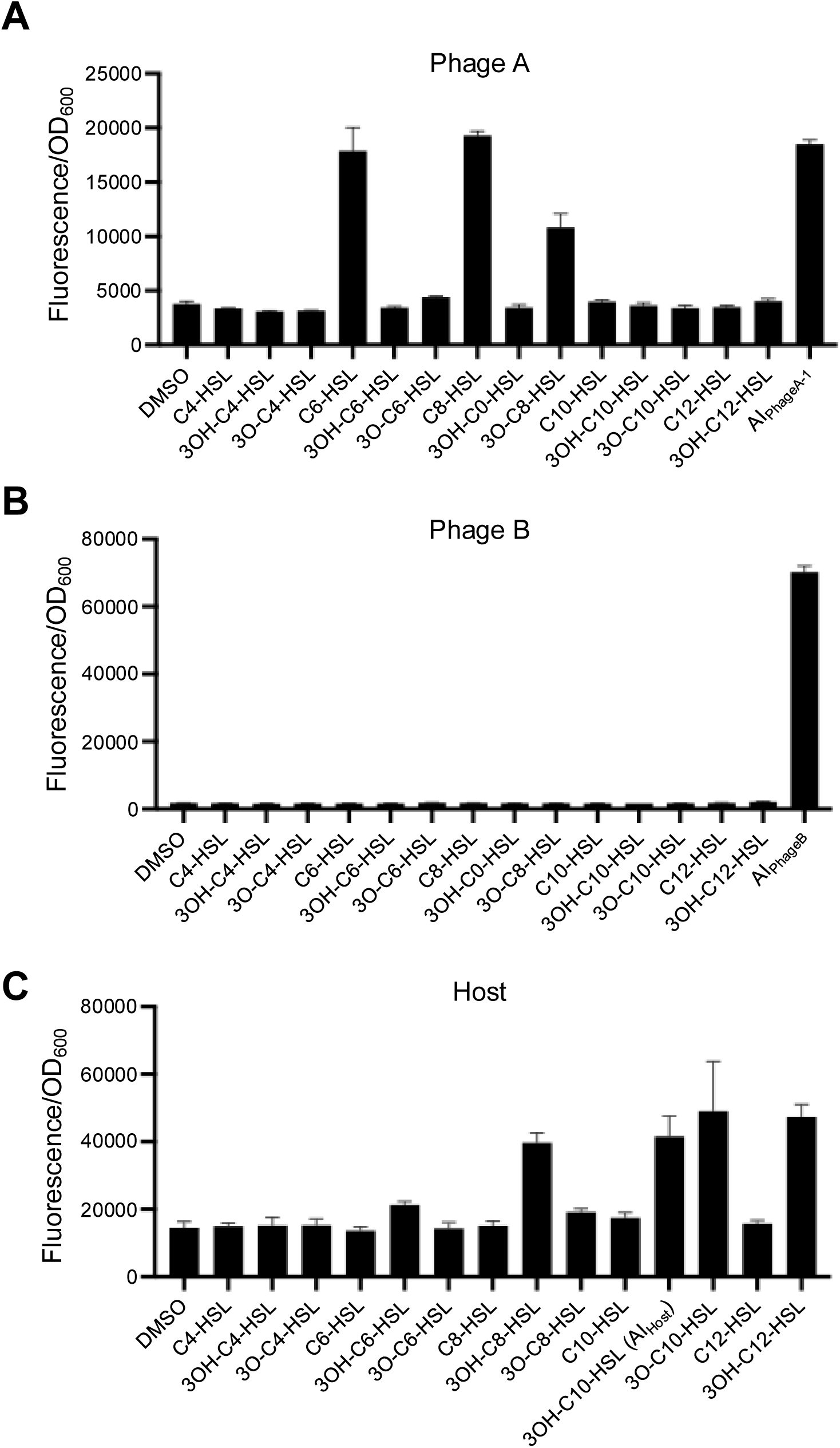
Ligand specifities of prophage and host LuxR receptors. (**A**-**C**) responses of the Prophage A (**A**), Prophage B (**B**), and host (**C**) LuxR receptors to a panel of chemically synthesized HSLs differing in chain length and oxidation state. The abbreviation 3O refers to 3-oxo. The prophage P_Lysogeny_–*mScarlet-I-xre–luxR* reporters were introduced into a Δ*3luxI* Δ*3luxR* strain. The host P*_paaZ2_*–*mScarlet-I* reporter was introduced into the Δ*3luxI* Δ*3luxR* strain carrying the host *pgaR* receptor. Cultures were exposed to 10 μM of the indicated HSL and fluorescence was normalized to culture density (OD_600_). In each panel, data represent means ± SDs from three biological replicates.

**Figure S5.**
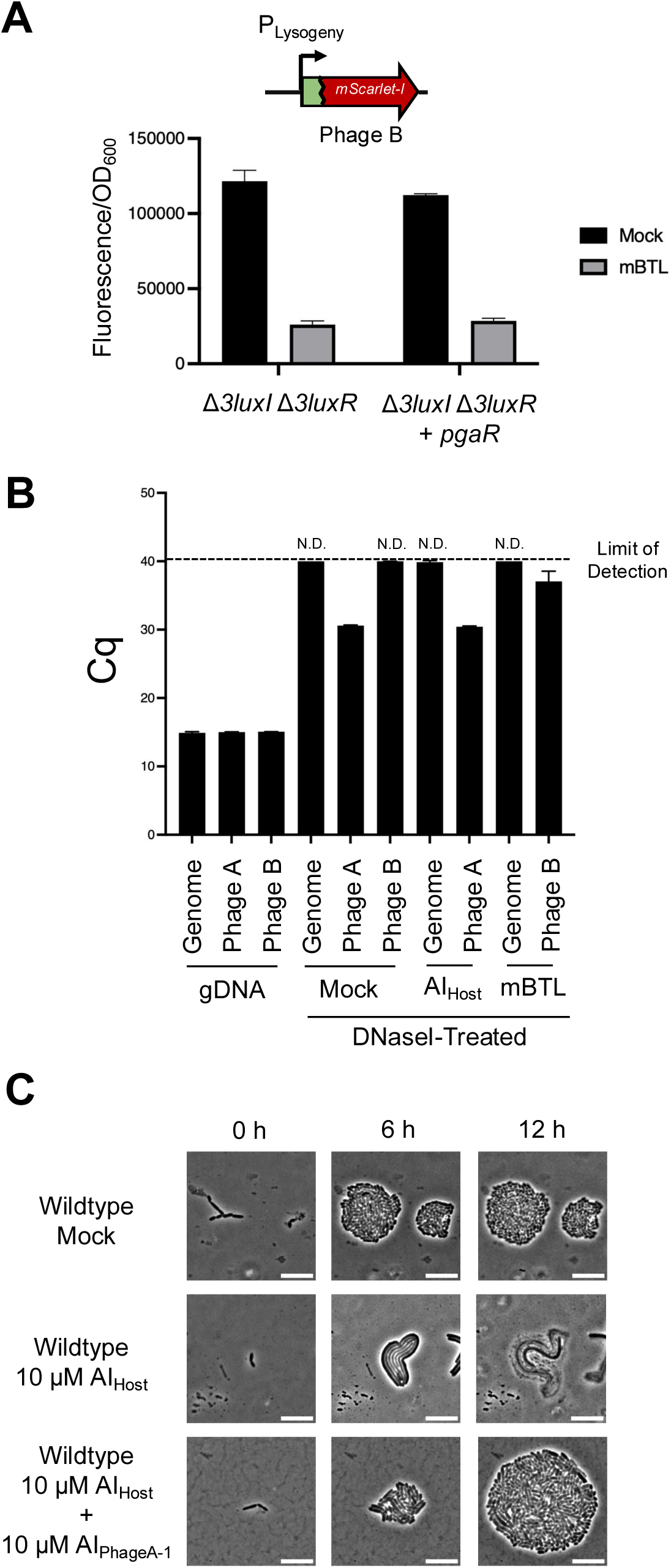
Inhibition of Prophage QS induces lytic growth. (**A**) Inhibition of the Prophage B QS system by the synthetic antagonist mBTL. The Prophage B P_Lysogeny_–*mScarlet-I-xre–luxR-luxI* reporter was introduced into either a Δ*3luxI* Δ*3luxR* strain or a derivative carrying the host *pgaR* receptor. Cultures were treated with DMSO (Mock) or 100 μM mBTL. mScarlet-I fluorescence was normalized to OD_600_. Data represent means ± SDs from four biological replicates. (**B**) DNase I protection assay for detecting extracellular phage particles of Phage A and Phage B. Purified genomic DNA (gDNA) or DNase I-treated cell-free fluids from cultures treated with DMSO (Mock), 10 μM AI_Host_, or 100 μM mBTL were analyzed by qPCR using primer sets targeting the bacterial chromosome (Genome), Prophage A (Phage A), or Prophage B (Phage B). Bars show Cq values for each qPCR. The dashed line indicates the assay limit of detection. Samples that failed to amplify were assigned a Cq value of 40 for plotting purposes and are designated as not detected (n.d.). (**C**) Representative time-lapse microscopy images of the designated *P. inhibens* T5^T^ strains at the times following the indicated treatment. Each treatment was independently replicated three times; in each replicate, approximately 10 microcolonies were imaged over time. Scale bar, 10 μm.

**Figure S6.**
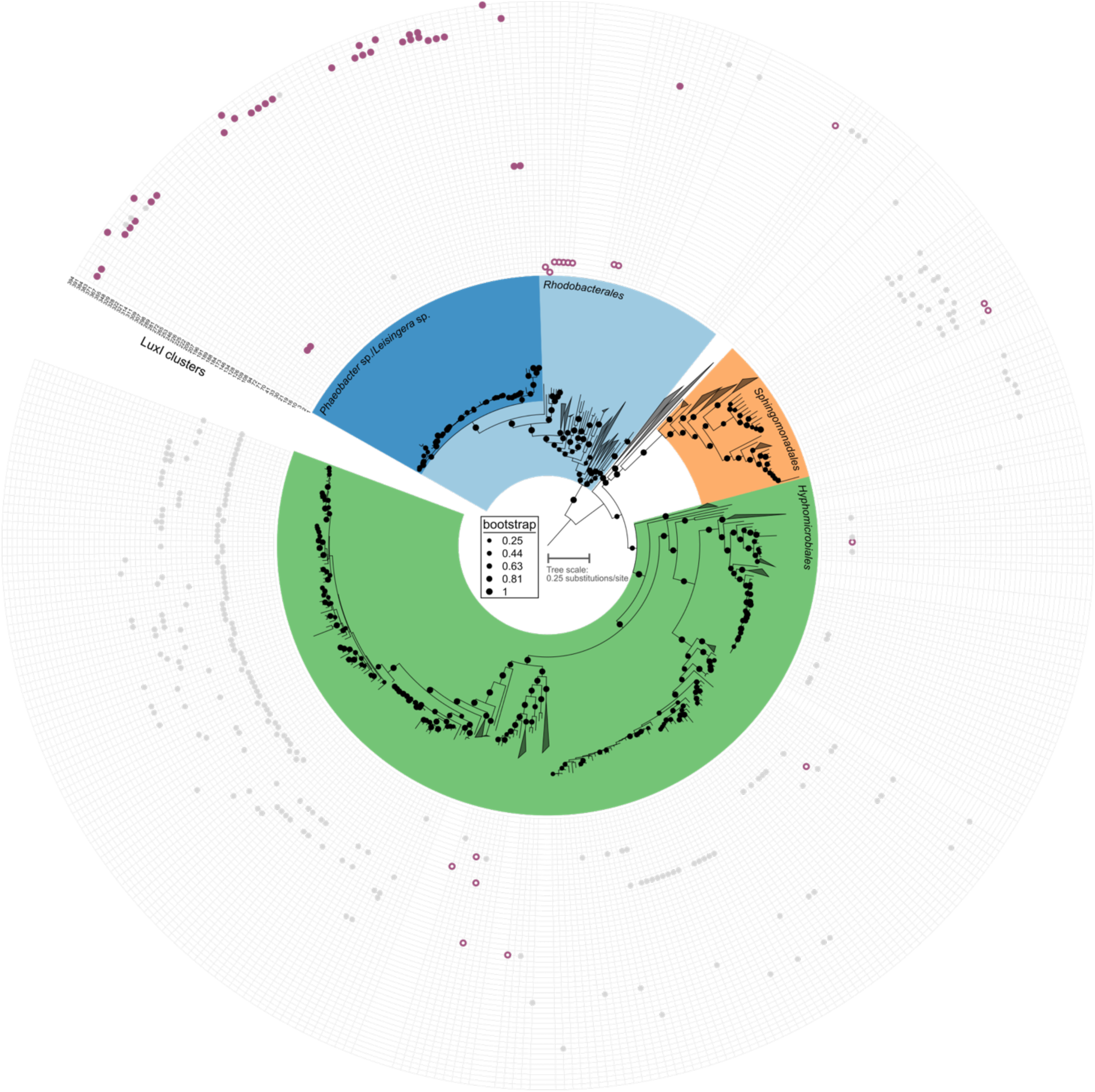
Within Alphaproteobacteria genomes that harbor multiple *luxI* genes, 54 *luxI* clusters display patterns of horizontal gene transfer. Core-genome phylogenetic tree of Alphaproteobacteria genomes that harbor multiple *luxI* genes overlayed with LuxI protein clusters that display a pattern of horizontal inheritance. Bootstrap support for each node is shown on the tree. Scale represents substitutions per amino acid site. LuxI proteins not predicted to be in a phage are shown as small gray points. LuxI proteins predicted to be harbored by a phage and found in an *xre-luxR-luxI* operon are shown as solid purple points. LuxI proteins predicted to be harbored by a phage but not found in an *xre-luxR-luxI* operon are shown as empty purple points.

**Figure S7.**
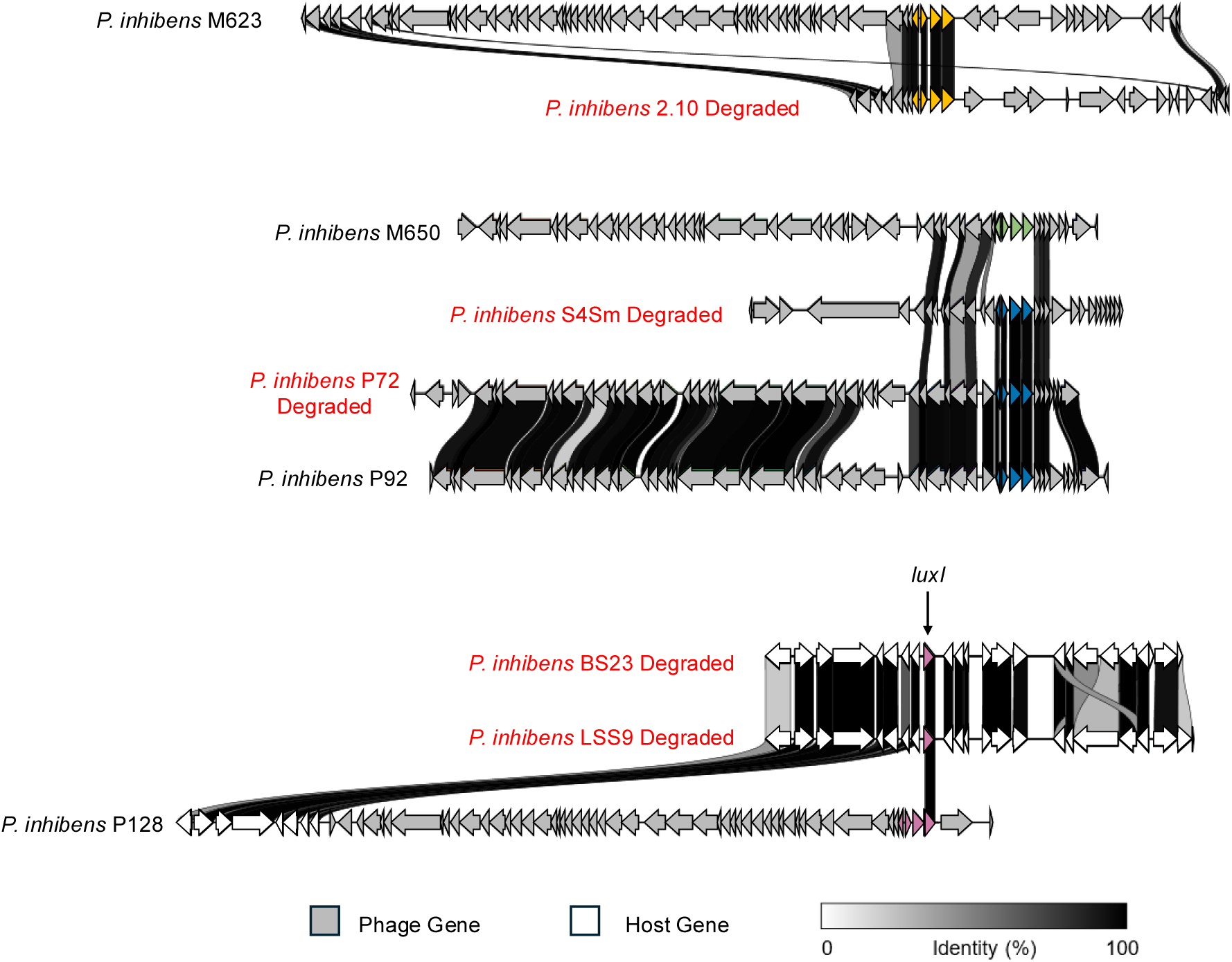
QS modules are retained within degraded prophages. Comparative genomic analysis of QS modules residing in degraded prophages. Gene neighborhoods surrounding each QS module were compared to their closest prophage relatives using Clinker. Colored genes denote QS modules and are shaded according to the color scheme used in Fig. 6. Gray arrows represent phage-derived genes. White arrows represent bacterial genes. Black connections indicate regions of sequence similarity with the degree of shading corresponding to amino acid sequence identity as shown in the scale bar.

**Figure S8.**
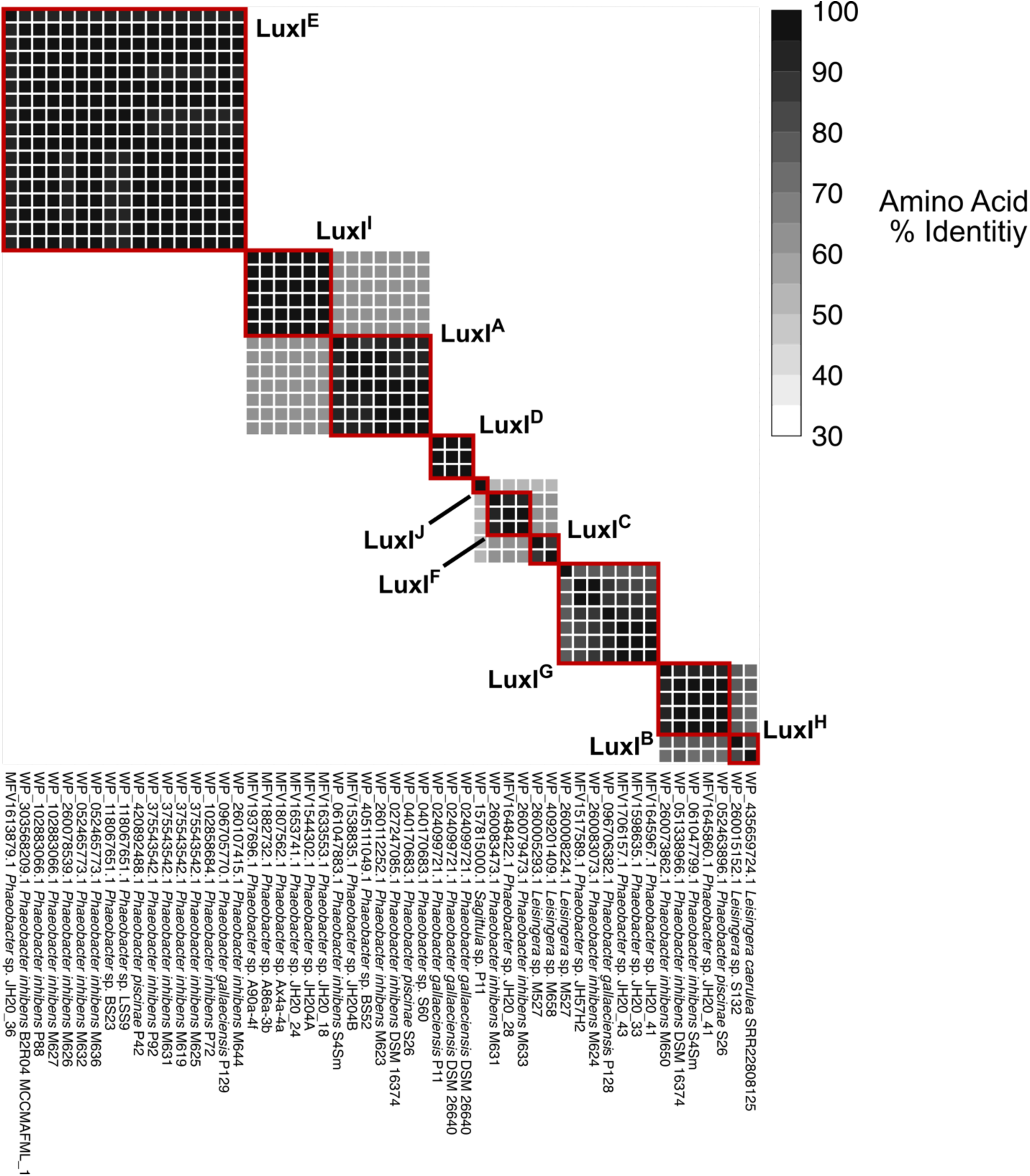
LuxI proteins encoded by phage *xre-luxR-luxI* operons cluster into distinct LuxI types. Pairwise heatmap of amino acid identities between the phage-encoded LuxI proteins. Protein identifiers, host species, and strain name are provided below each column. LuxI clusters determined with an 80% amino acid identity cutoff are indicated with red squares and labels.

**Figure S9.**
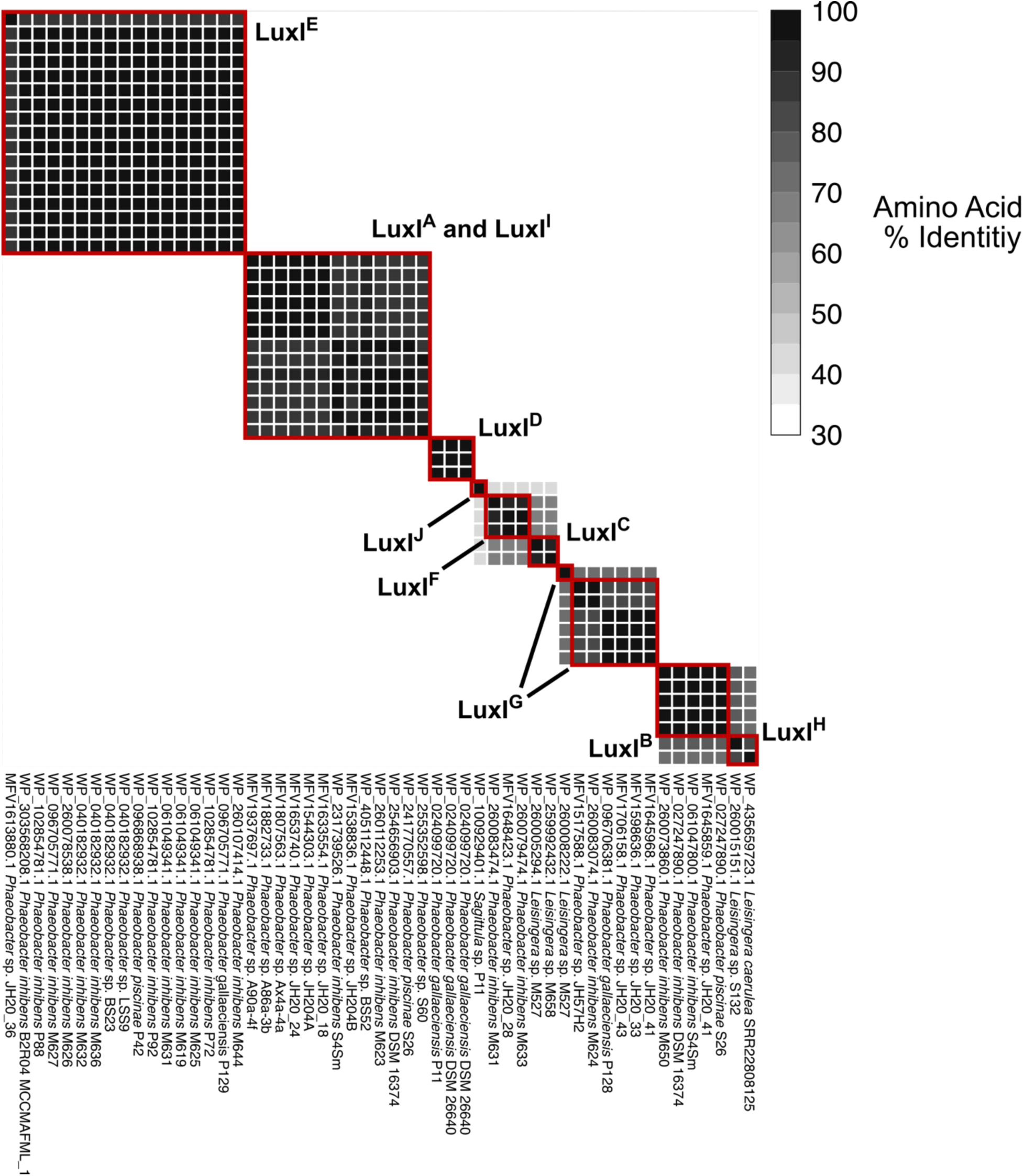
LuxR proteins encoded by phage *xre-luxR-luxI* operons cluster similarly to LuxI proteins with some exceptions. Pairwise heatmap of amino acid identities between the phage-encoded LuxR proteins. Protein identifiers, host species, and strain name are provided below each column. LuxR clusters determined with an 80% amino acid identity cutoff are indicated with red squares. Labels indicate the LuxI types associated with each LuxR cluster.

**Figure S10.**
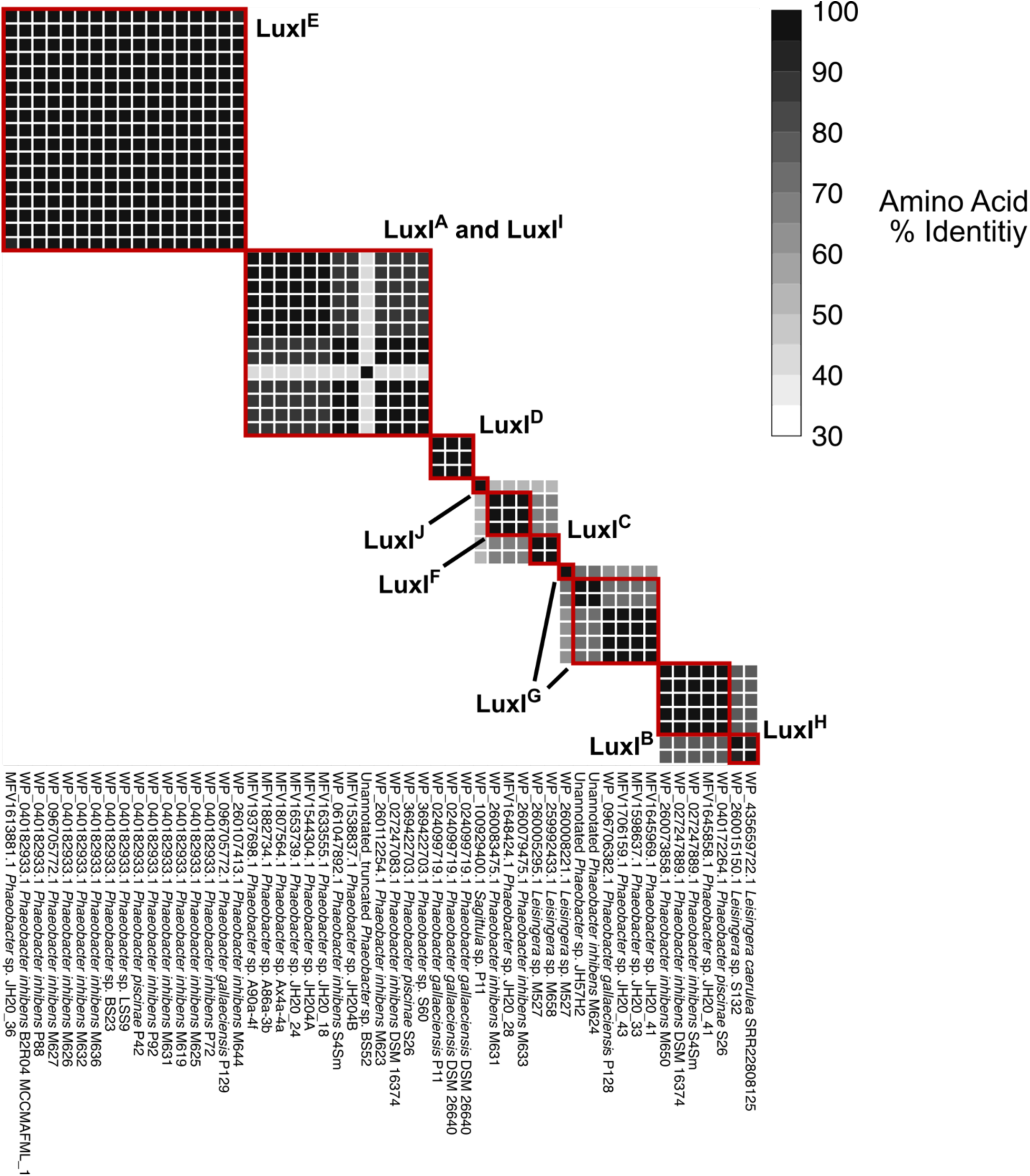
XRE proteins encoded by phage *xre-luxR-luxI* operons cluster similarly to LuxI proteins with some exceptions. Pairwise heatmap of amino acid identities between the phage-encoded XRE proteins. Protein identifiers, host species, and strain name are provided below each column. XRE clusters determined with an 80% amino acid identity cutoff are indicated with red squares. Labels indicate the LuxI types associated with each XRE cluster. ‘Unannotated’ indicates manually annotated proteins without identifiers. ‘Truncated’ indicates a frameshift with a premature STOP.

**Figure S11.**
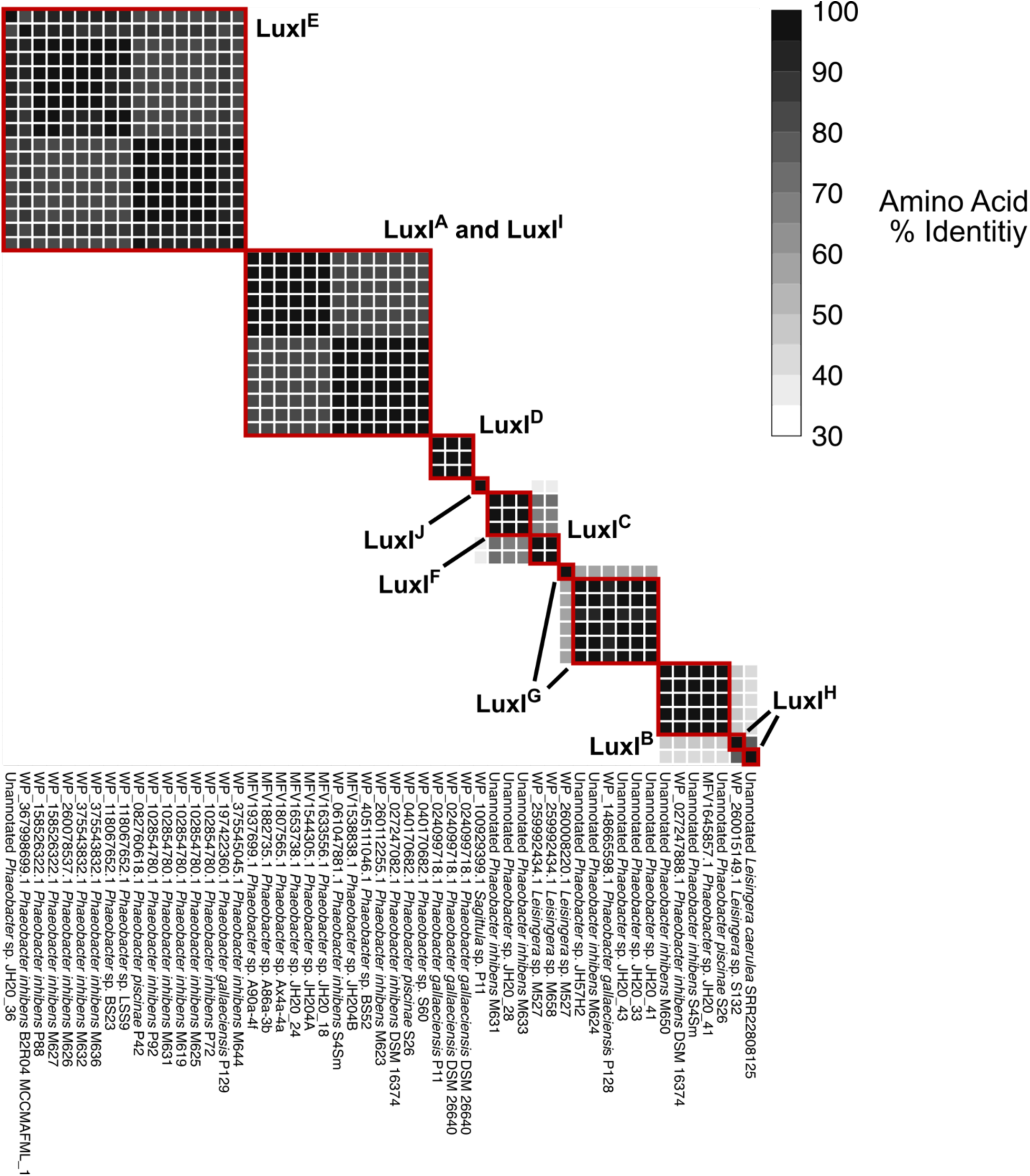
Hth proteins encoded by phage *xre-luxR-luxI* operons cluster similarly to LuxI proteins with some exceptions. Pairwise heatmap of amino acid identities between the phage-encoded Hth proteins. Protein identifiers, host species, and strain name are provided below each column. Hth clusters determined with an 80% amino acid identity cutoff are indicated with red squares. Labels indicate the LuxI types associated with each Hth cluster. ‘Unannotated’ indicates manually annotated proteins without identifiers.

**Figure S12.**
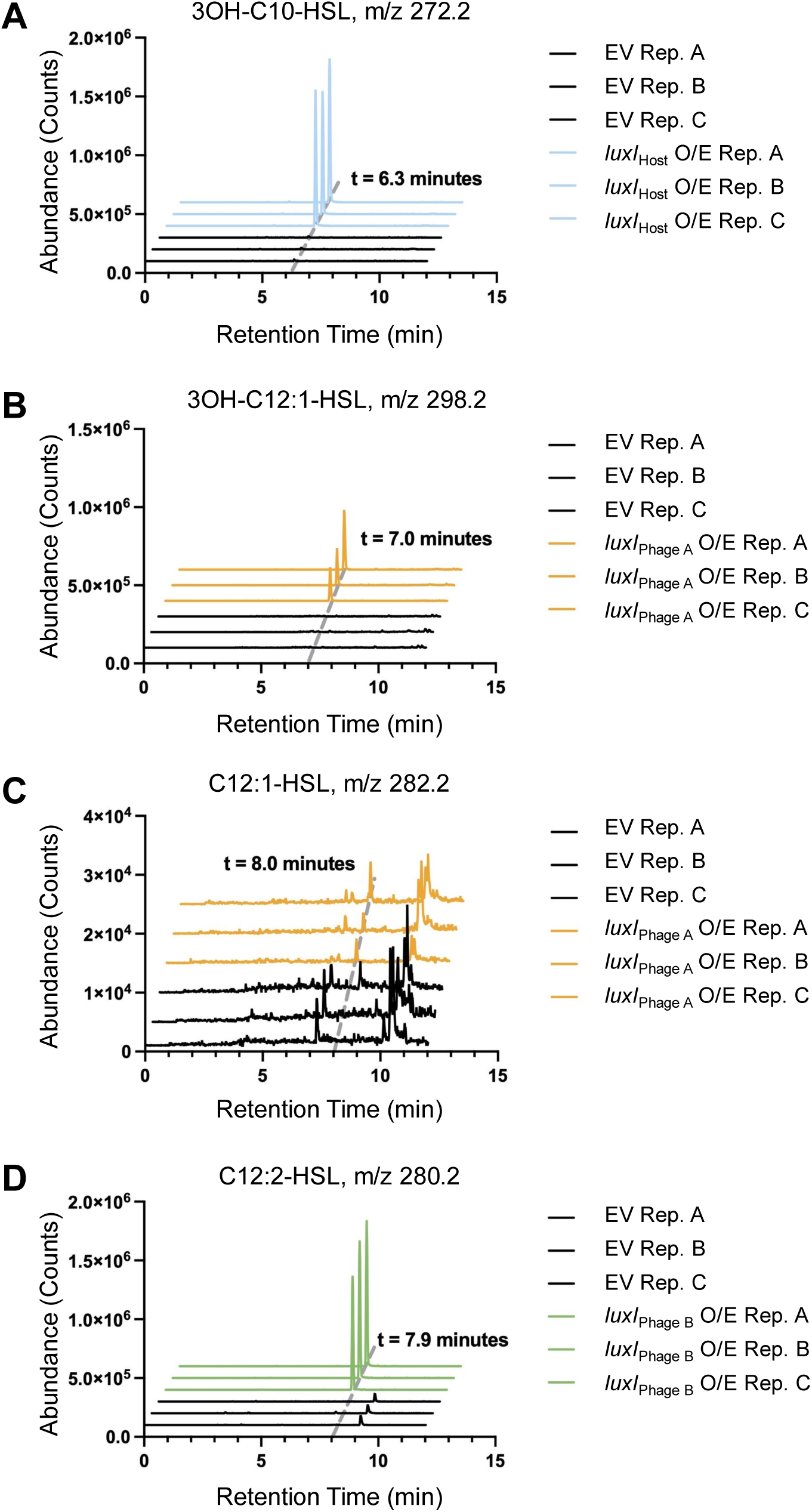
Identification of autoinducers produced by the three LuxI synthases in *P. inhibens* T5^T^. Extracted ion chromatographs (EIC) from preparations of *P. inhibens* DSM17395 carrying an empty vector (EV) or a plasmid overexpressing one of the three *luxI* genes studied here. (**A**) EIC for 3OH-C10-HSL (*m/z* 272.2). (**B**) EIC for 3OH-C12:1-HSL (*m/z* 298.2). (**C**) EIC for C12:1-HSL (*m/z* 282.2). (**D**) EIC for C12:2-HSL (*m/z* 280.2). Rep. designates biological replicates and O/E designates overexpression.

**Figure S13.**
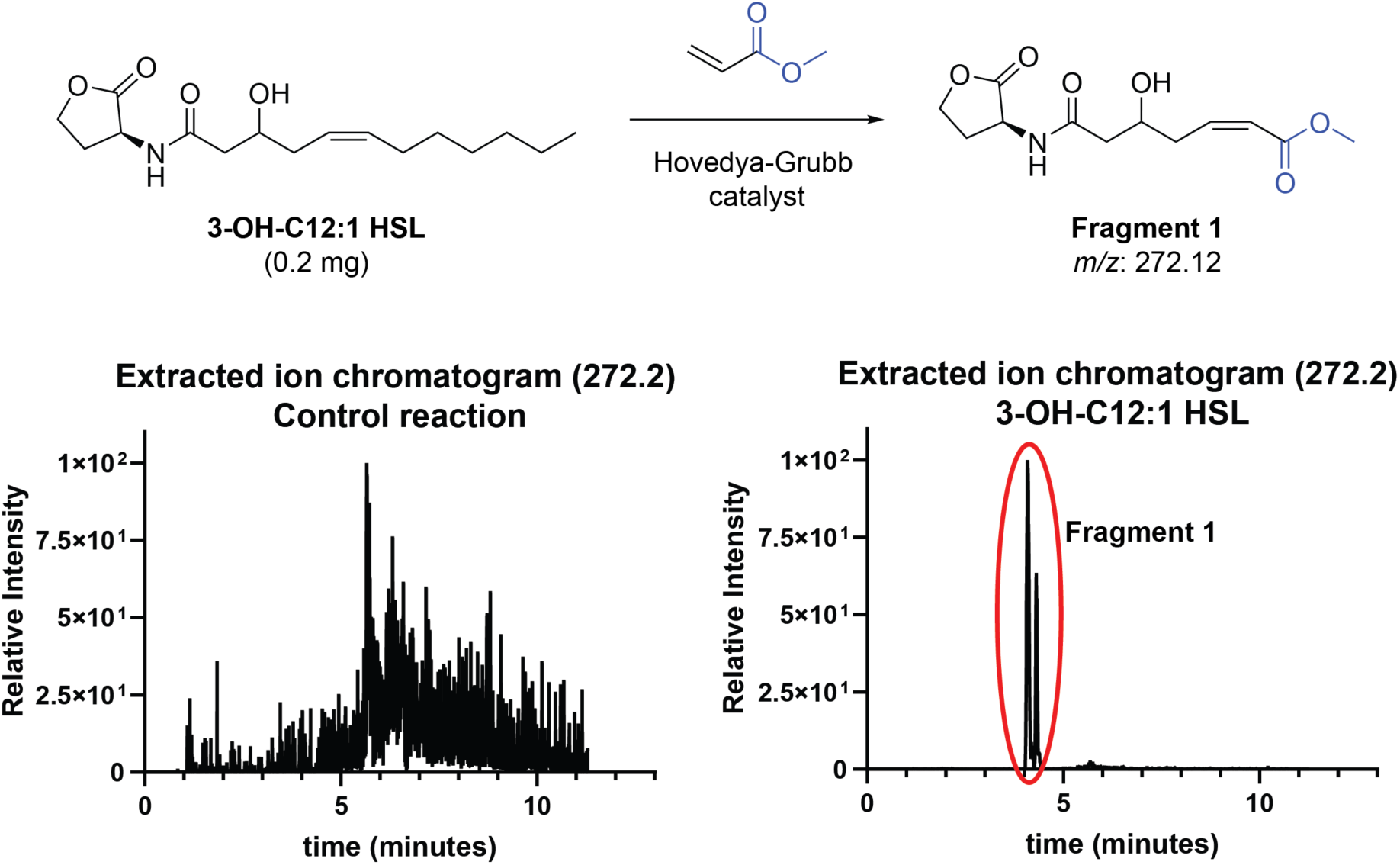
Olefin cross-metathesis of 3-OH-C12:1-HSL supporting acyl chain double bond location determination.

**Figure S14.**
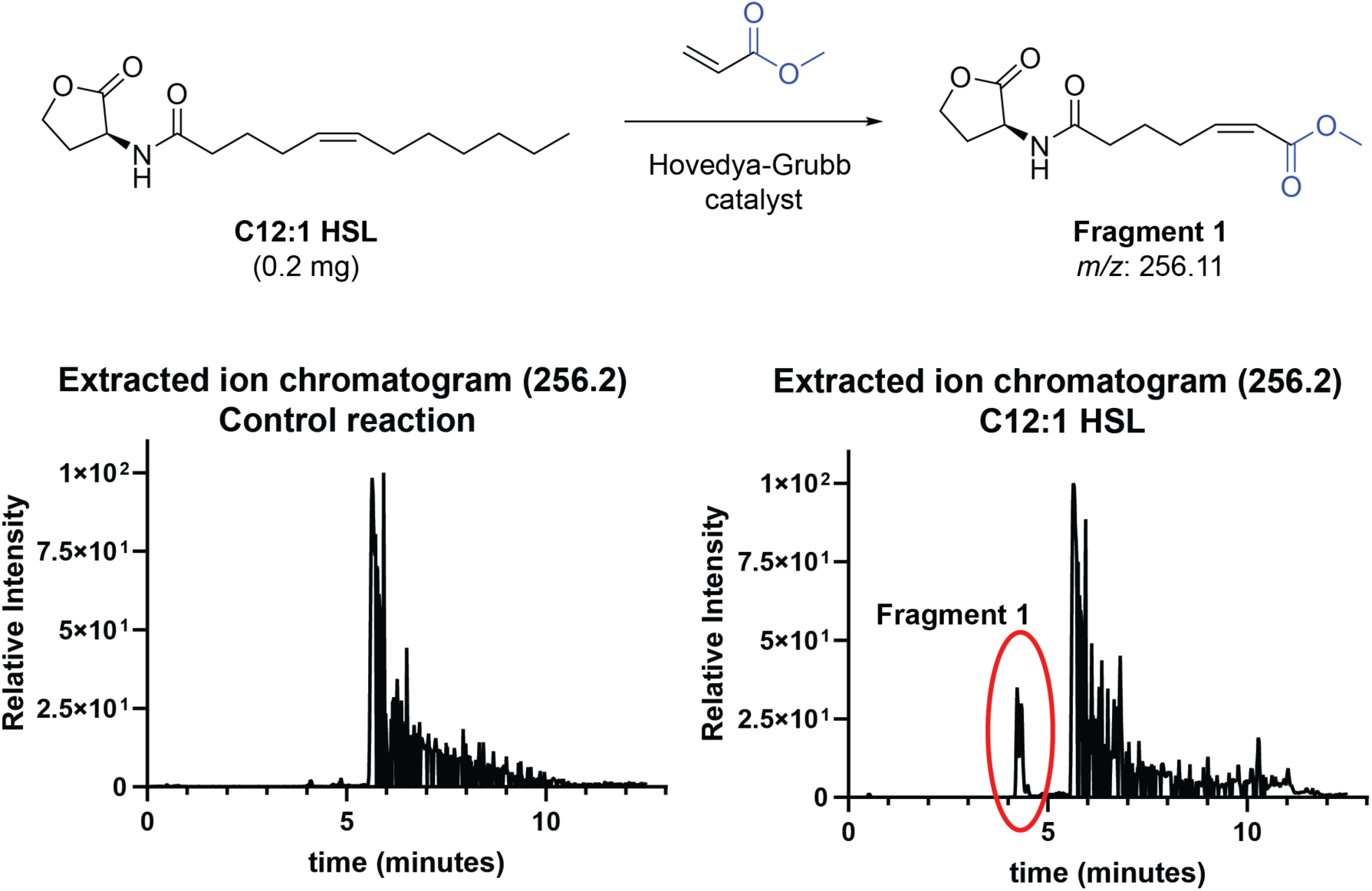
Olefin cross-metathesis of C12:1-HSL supporting acyl chain double bond location determination.

**Figure S15.**
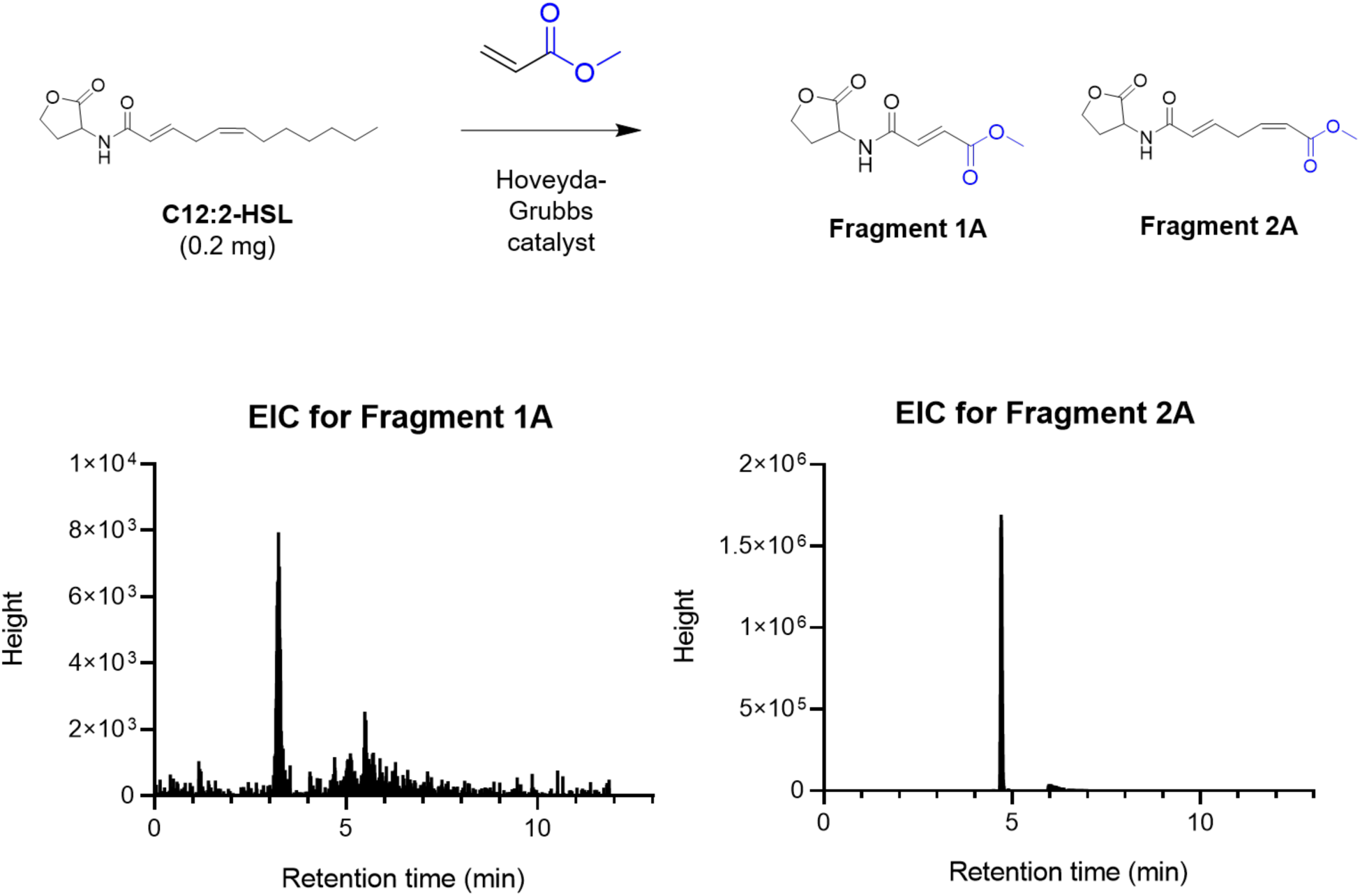
Olefin cross-metathesis of C12:2-HSL supporting acyl chain double bond location determination.

**Figure S16.**
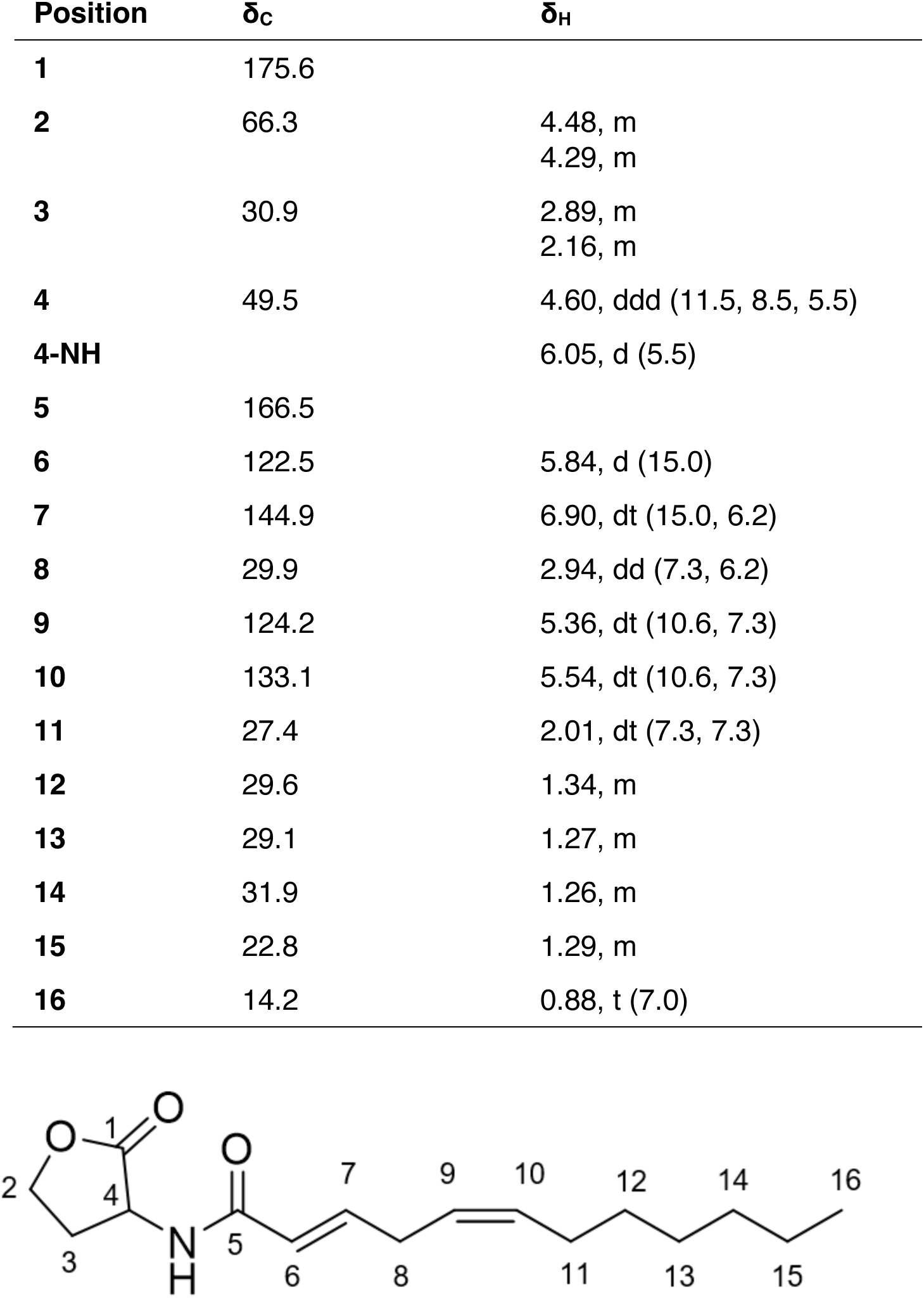
^1^H [ppm, mult. (*J* in Hz)] and ^13^C NMR data of C12:2-HSL in CDCl_3_.

**Figure S17.**
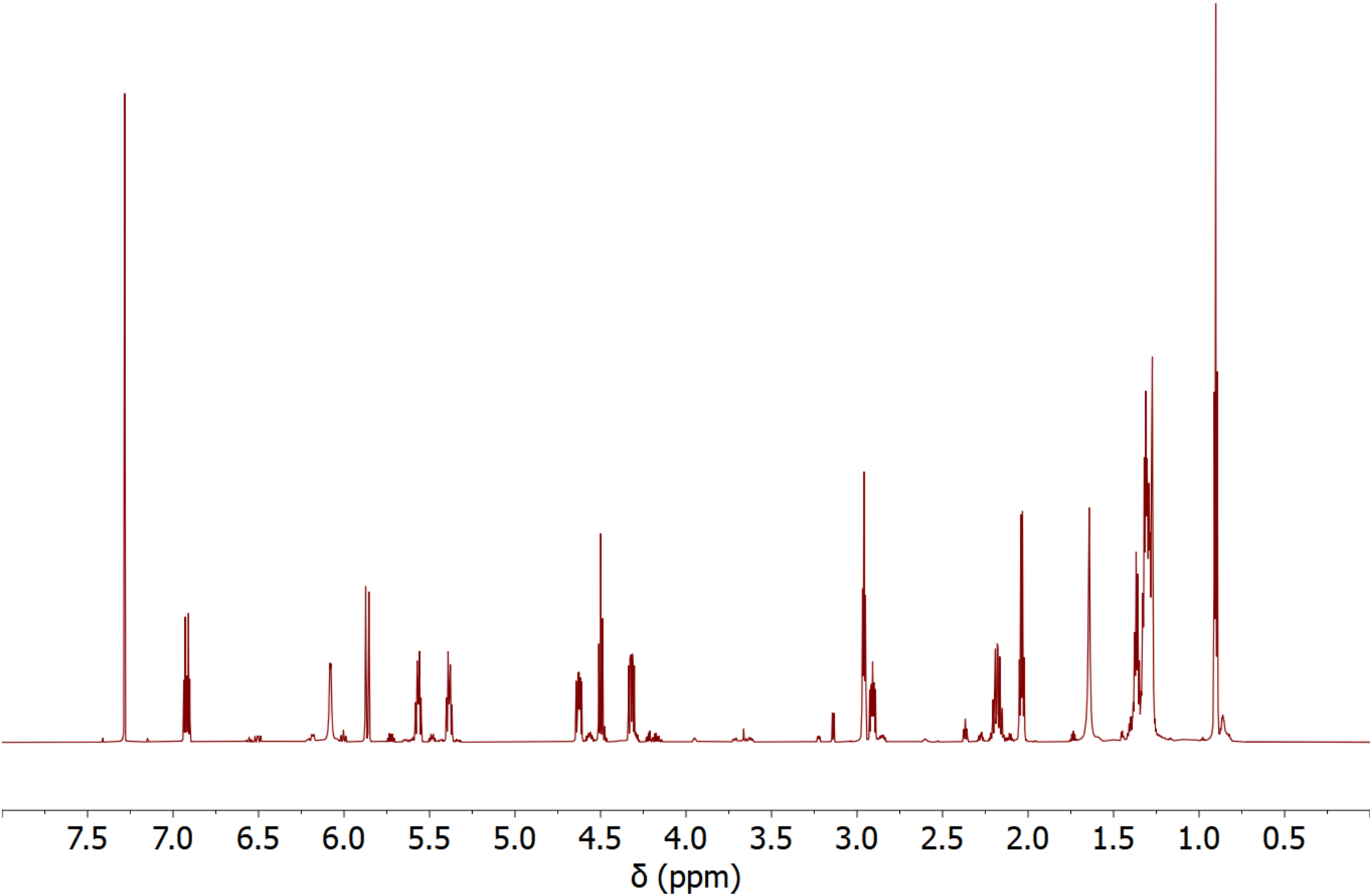
^1^H NMR spectrum of C12:2-HSL in CDCl_3_.

**Figure S18.**
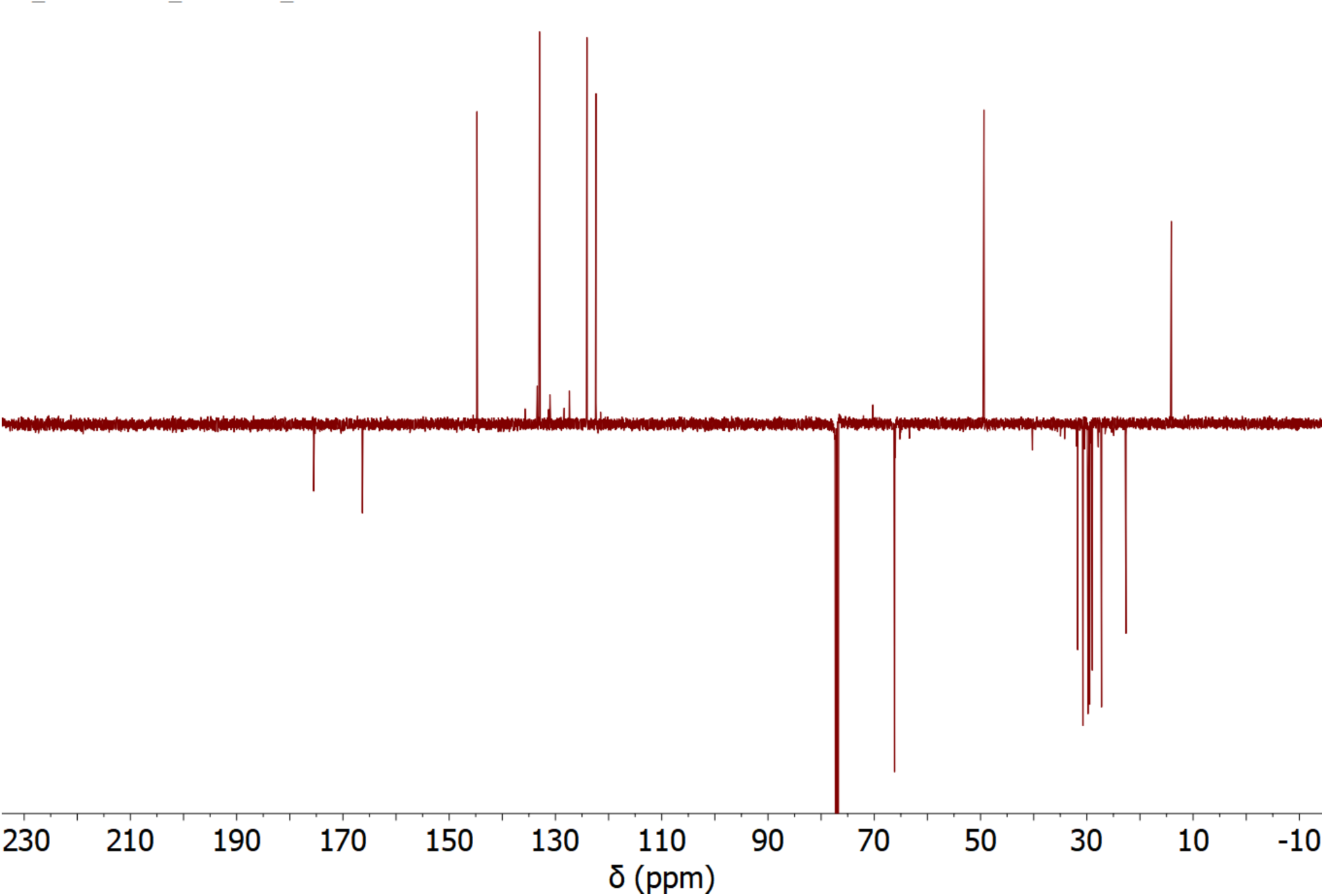
^13^C NMR spectrum of C12:2-HSL in CDCl_3_.

**Figure S19.**
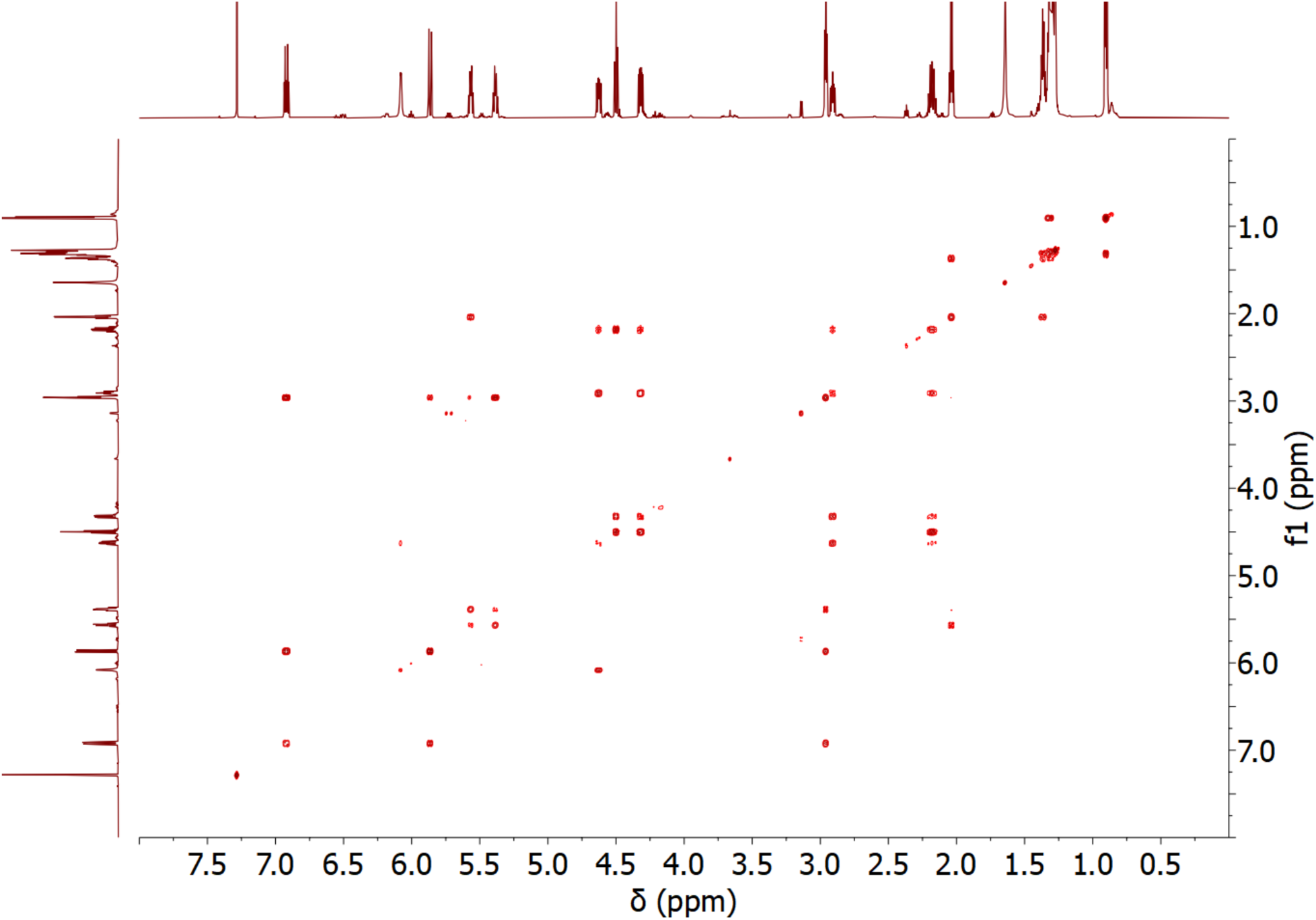
^1^H-^1^H COSY spectrum of C12:2-HSL in CDCl_3_.

**Figure S20.**
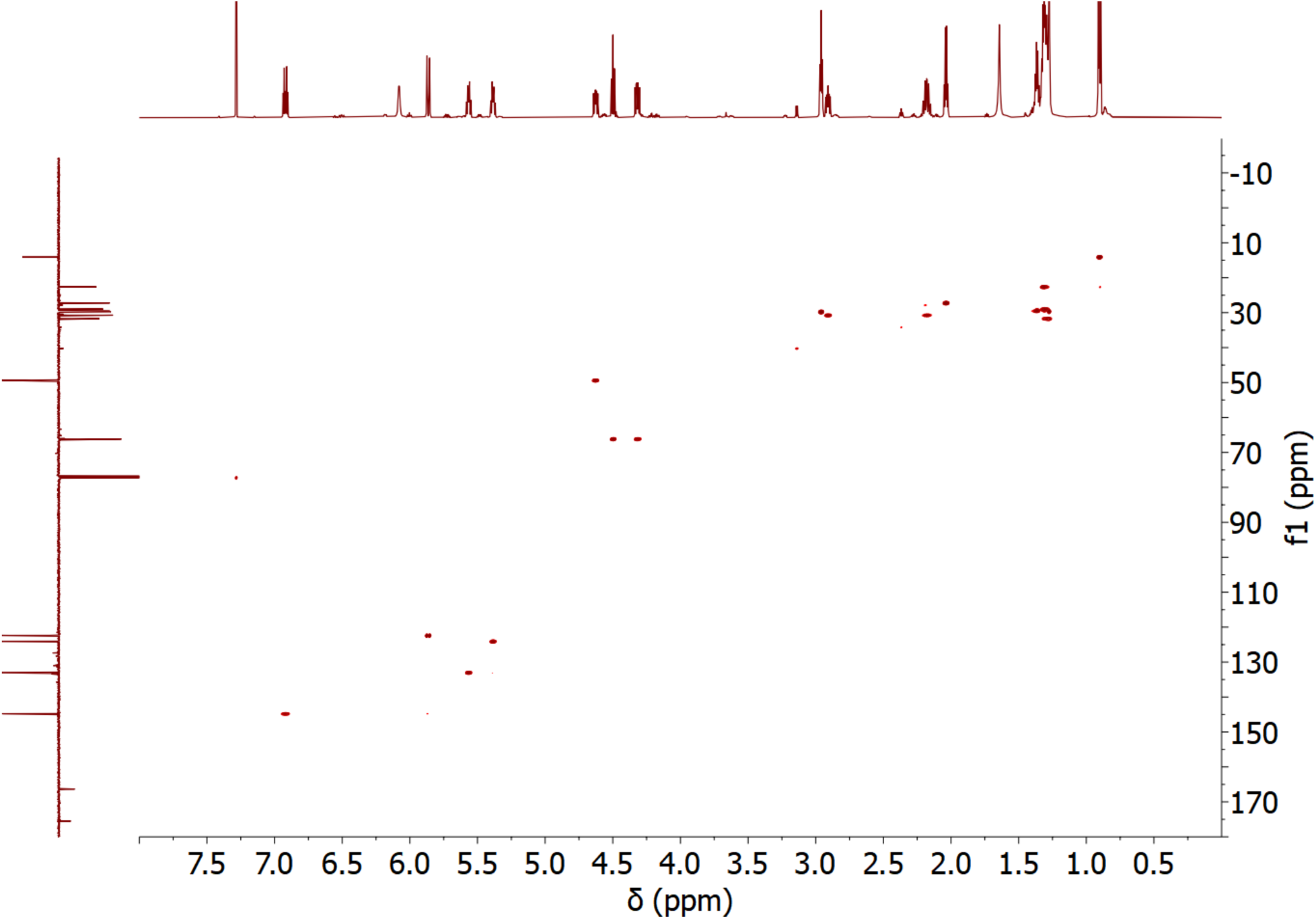
^1^H-^13^C HSQC spectrum of C12:2-HSL in CDCl_3_.

**Figure S21.**
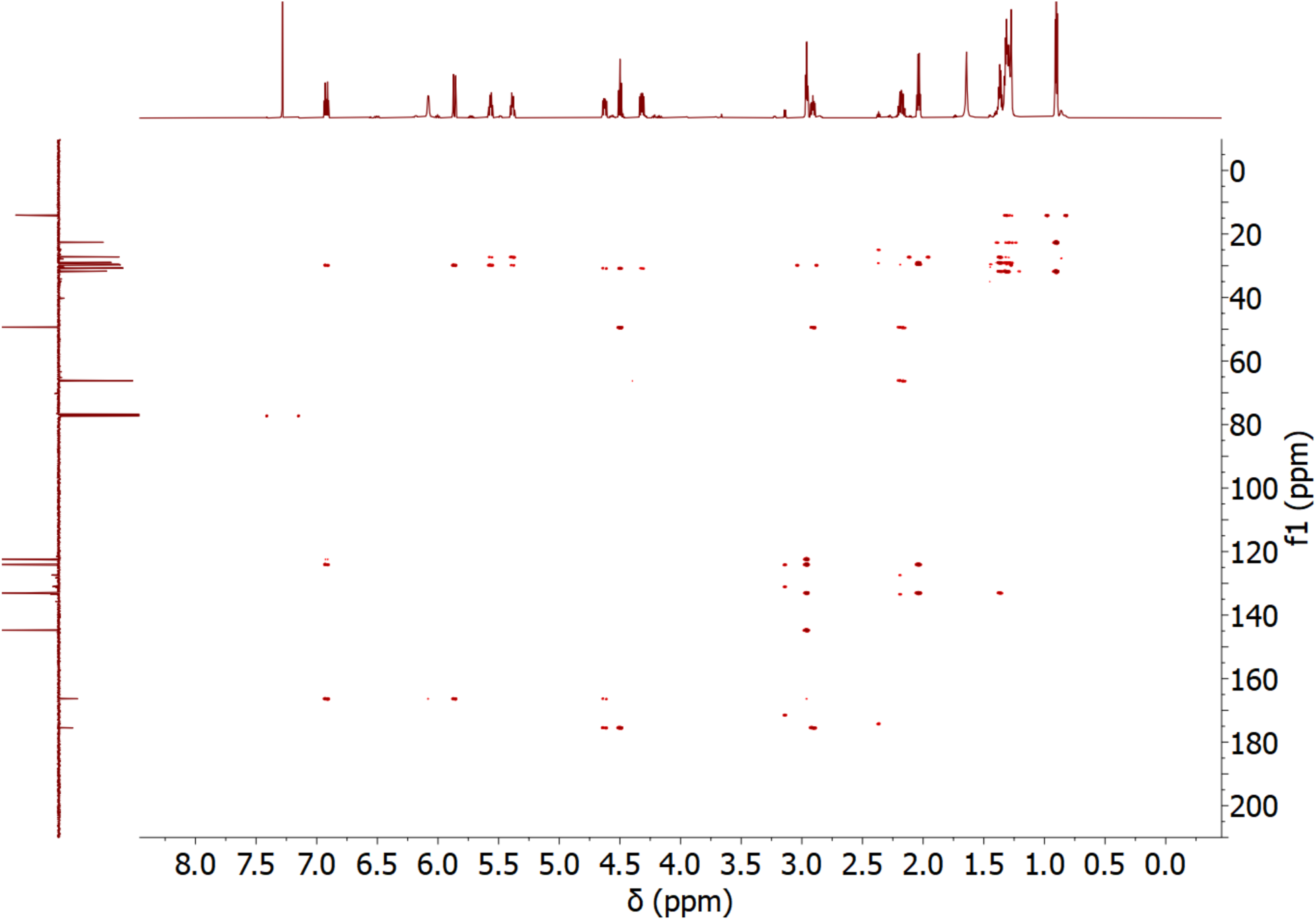
^1^H-^13^C HMBC spectrum of C12:2-HSL in CDCl_3_.

**Figure S22.**
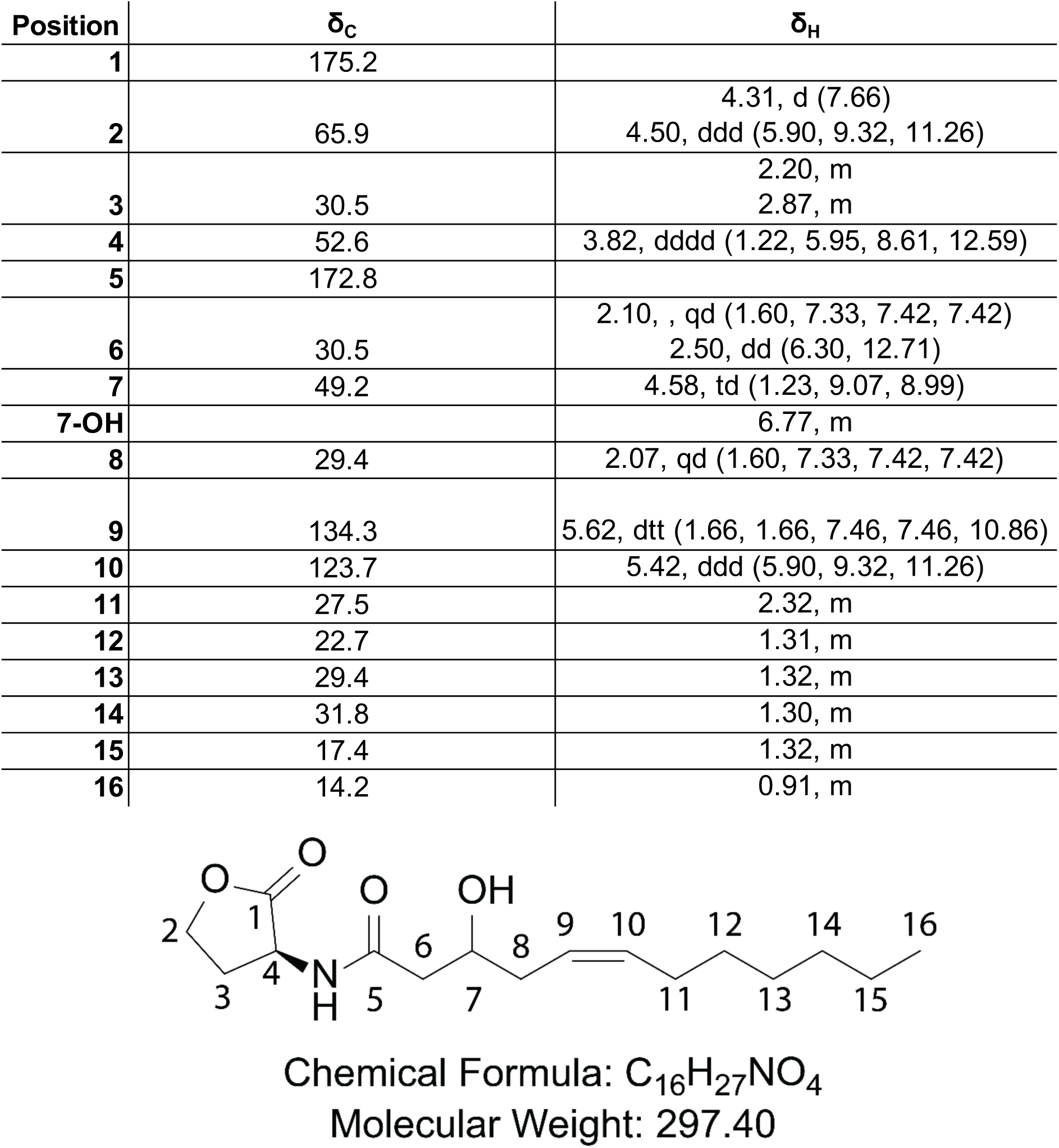
^1^H [ppm, mult. (*J* in Hz)] and ^13^C NMR data of 3-OH-C12:1-HSL in CDCl_3_.

**Figure S23.**
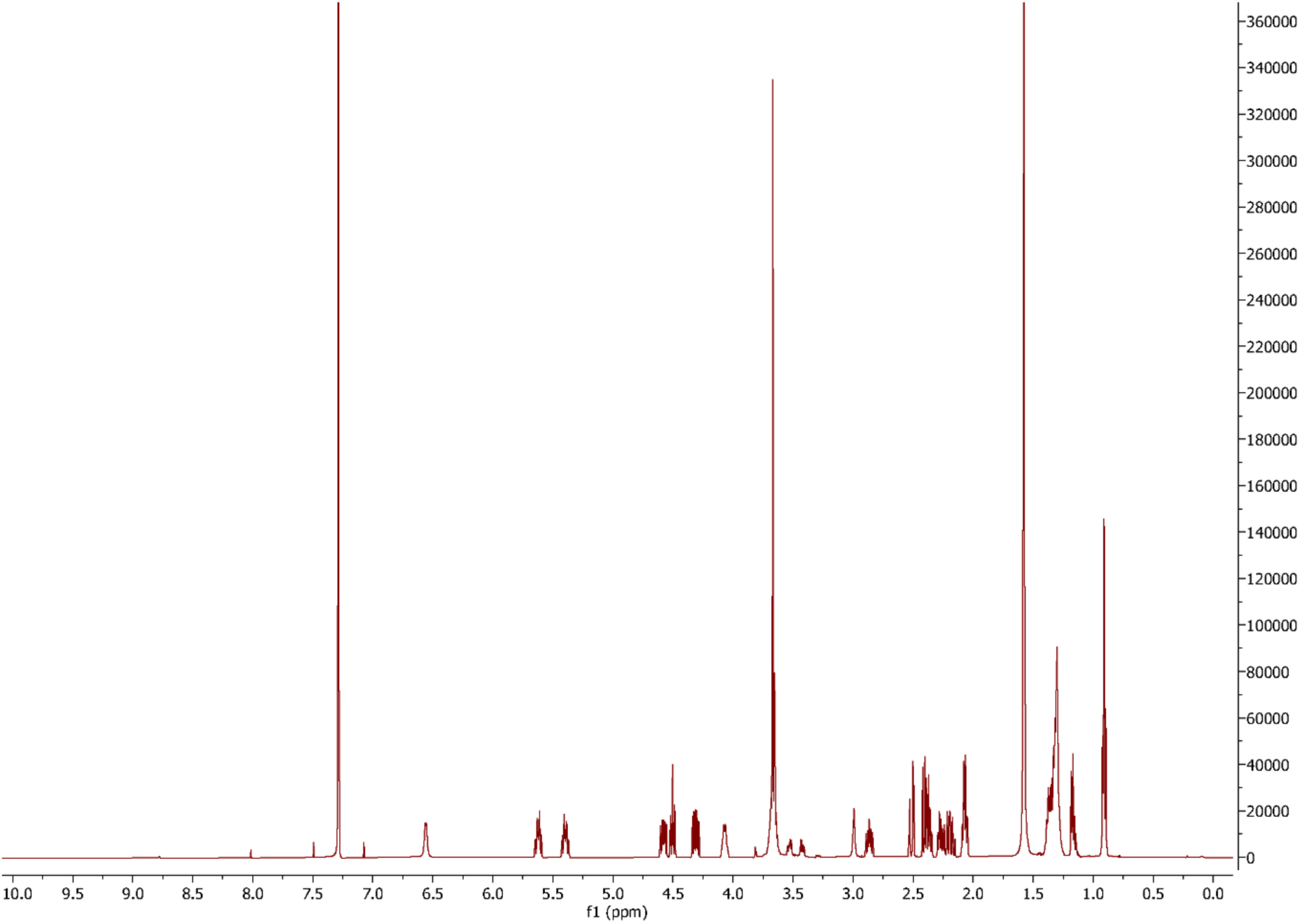
^1^H NMR spectrum of 3-OH-C12:1-HSL in CDCl_3_.

**Figure S24.**
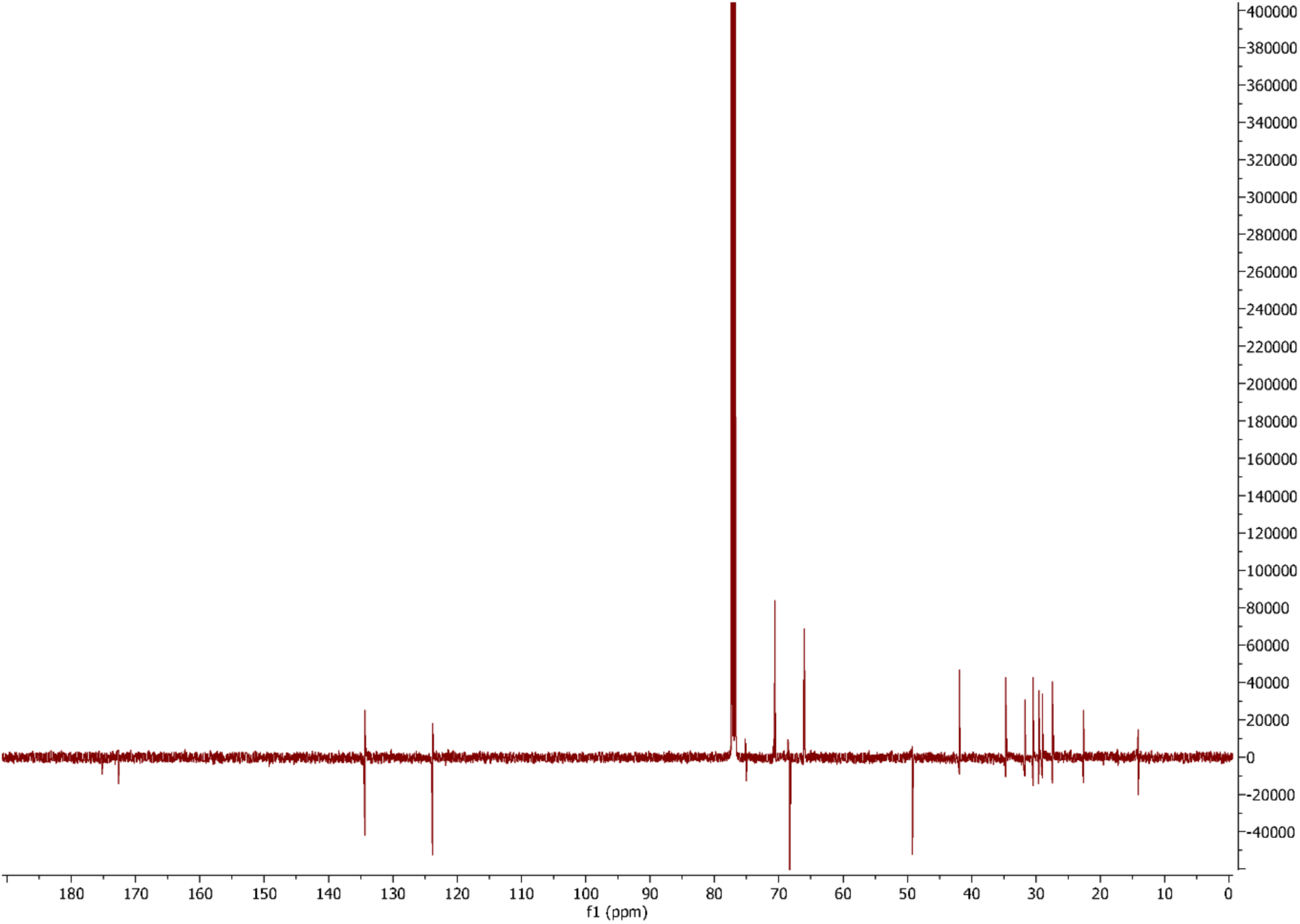
^13^C NMR spectrum of 3-OH-C12:1-HSL in CDCl_3_.

**Figure S25.**
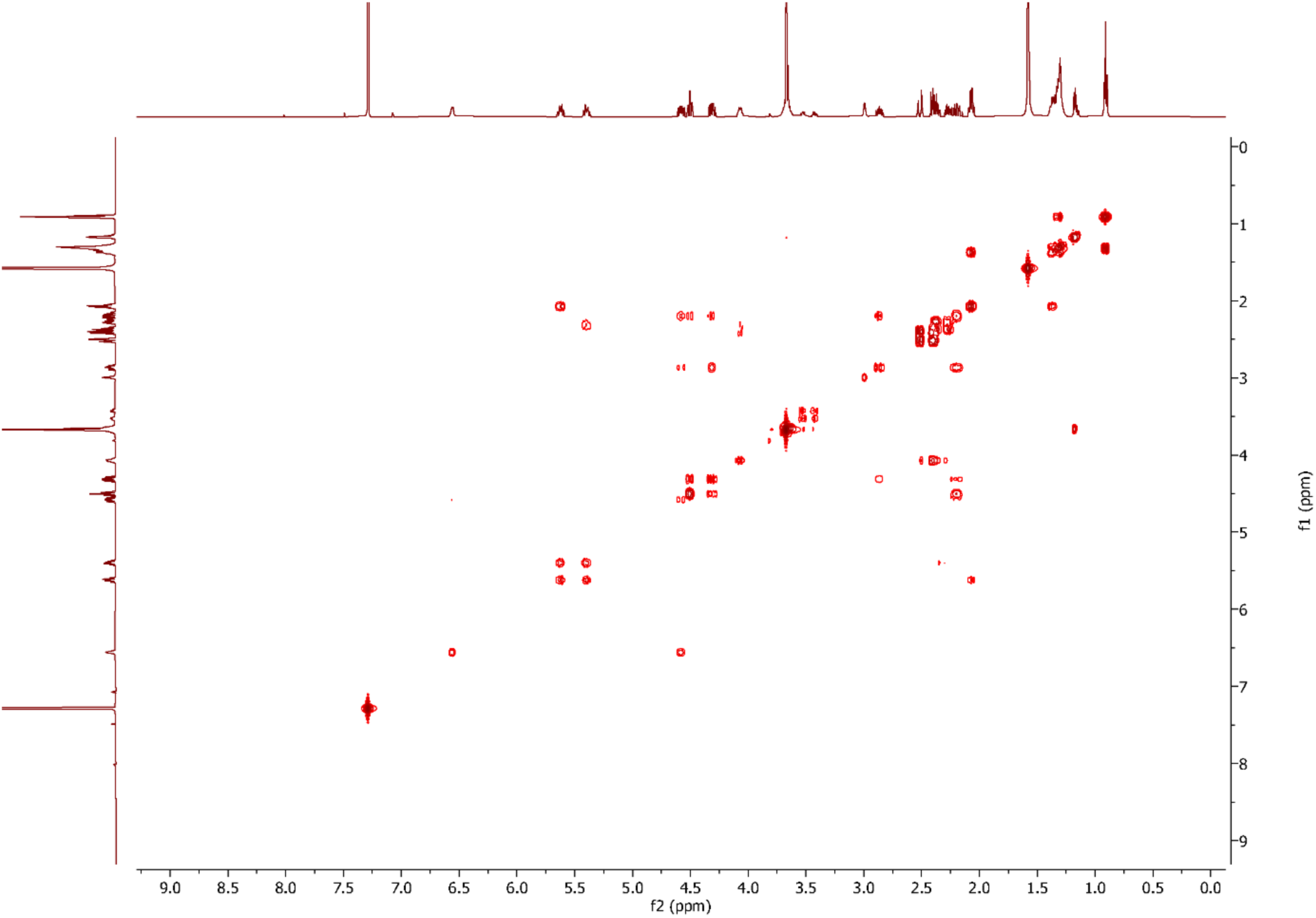
^1^H-^1^H COSY spectrum of 3-OH-C12:1-HSL in CDCl_3_.

**Figure S26.**
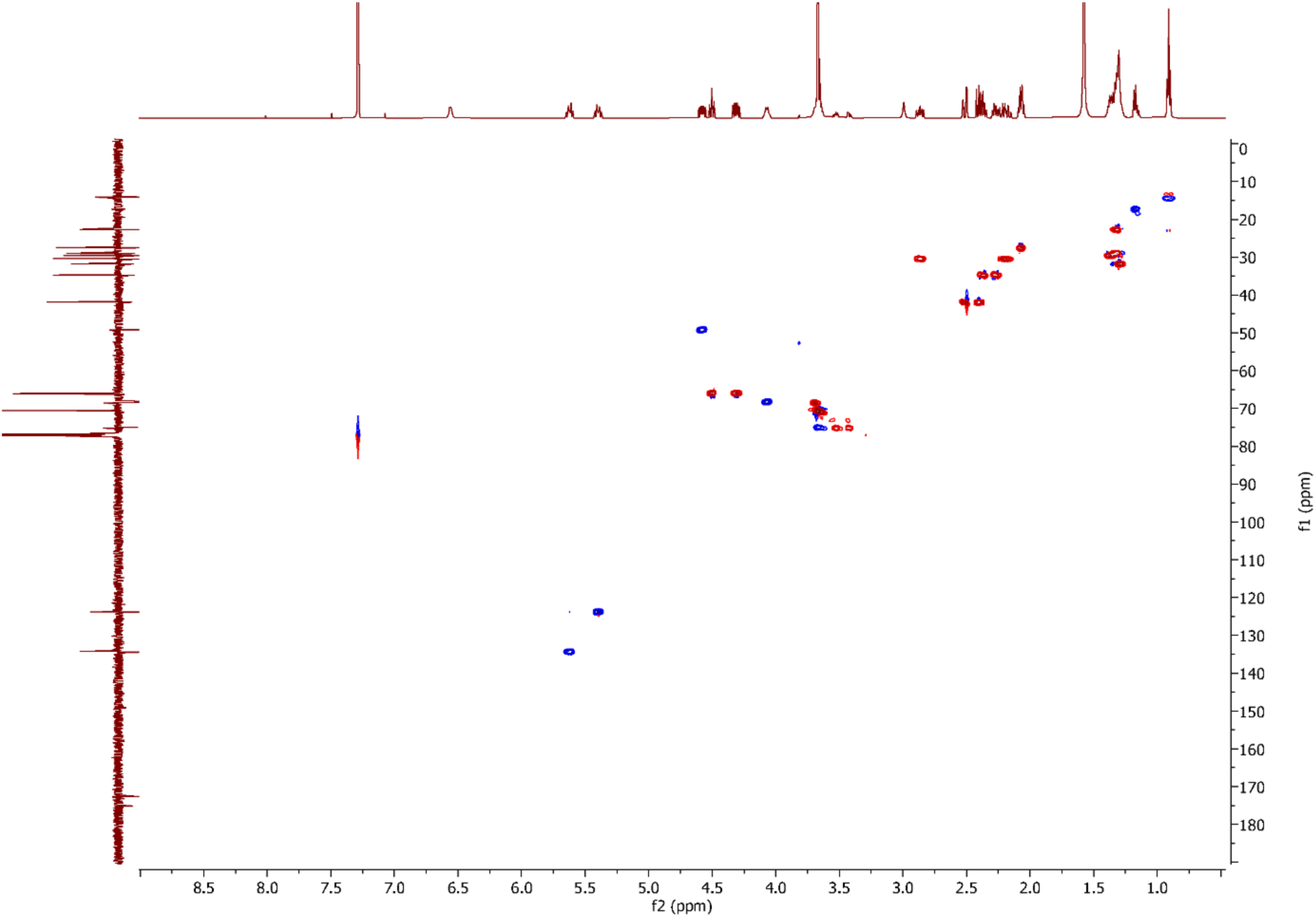
^1^H-^13^C HSQC spectrum of 3-OH-C12:1-HSL in CDCl_3_.

**Figure S27.**
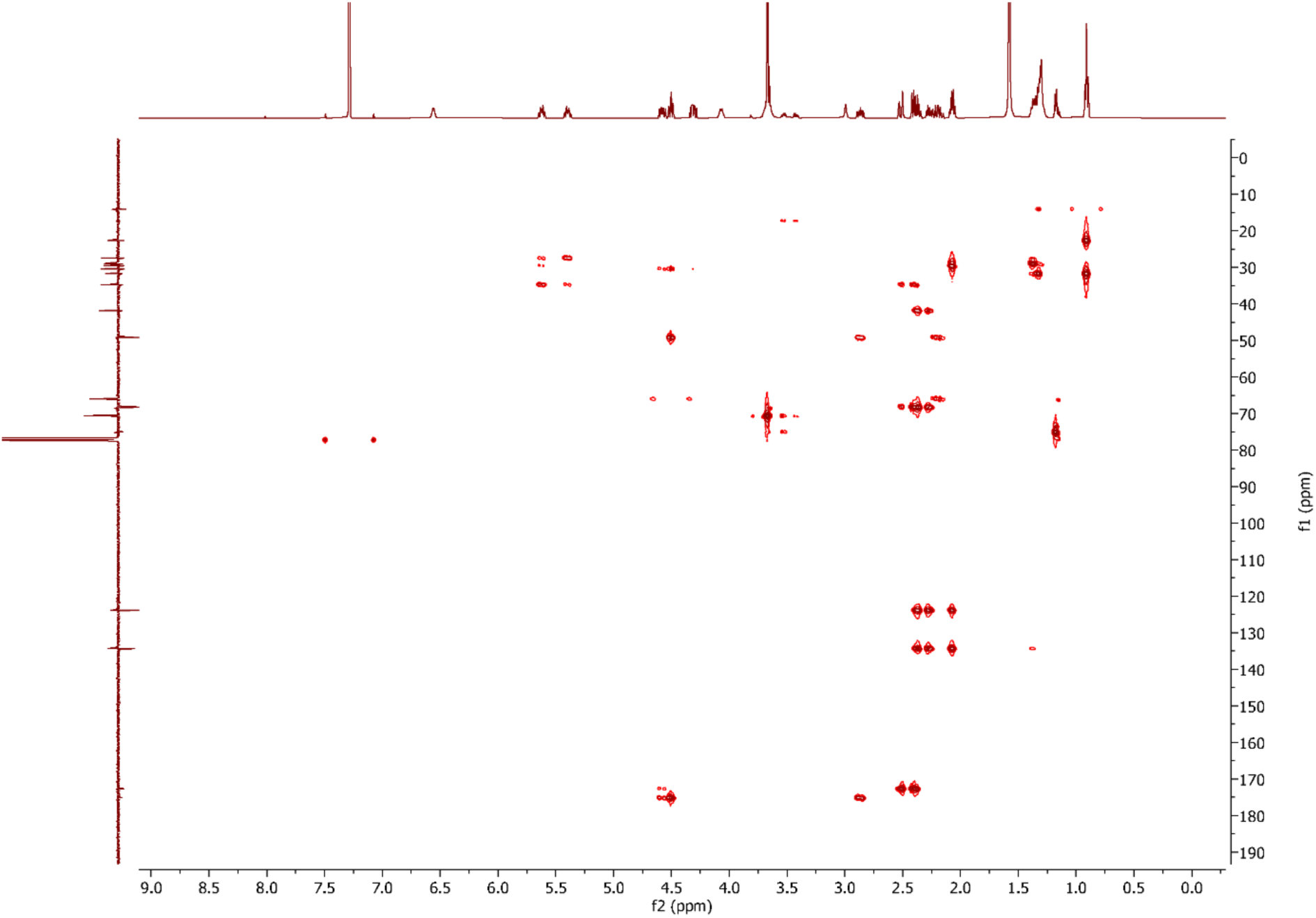
^1^H-^13^C HMBC spectrum of 3-OH-C12:1-HSL in CDCl_3_.

**Figure S28.**
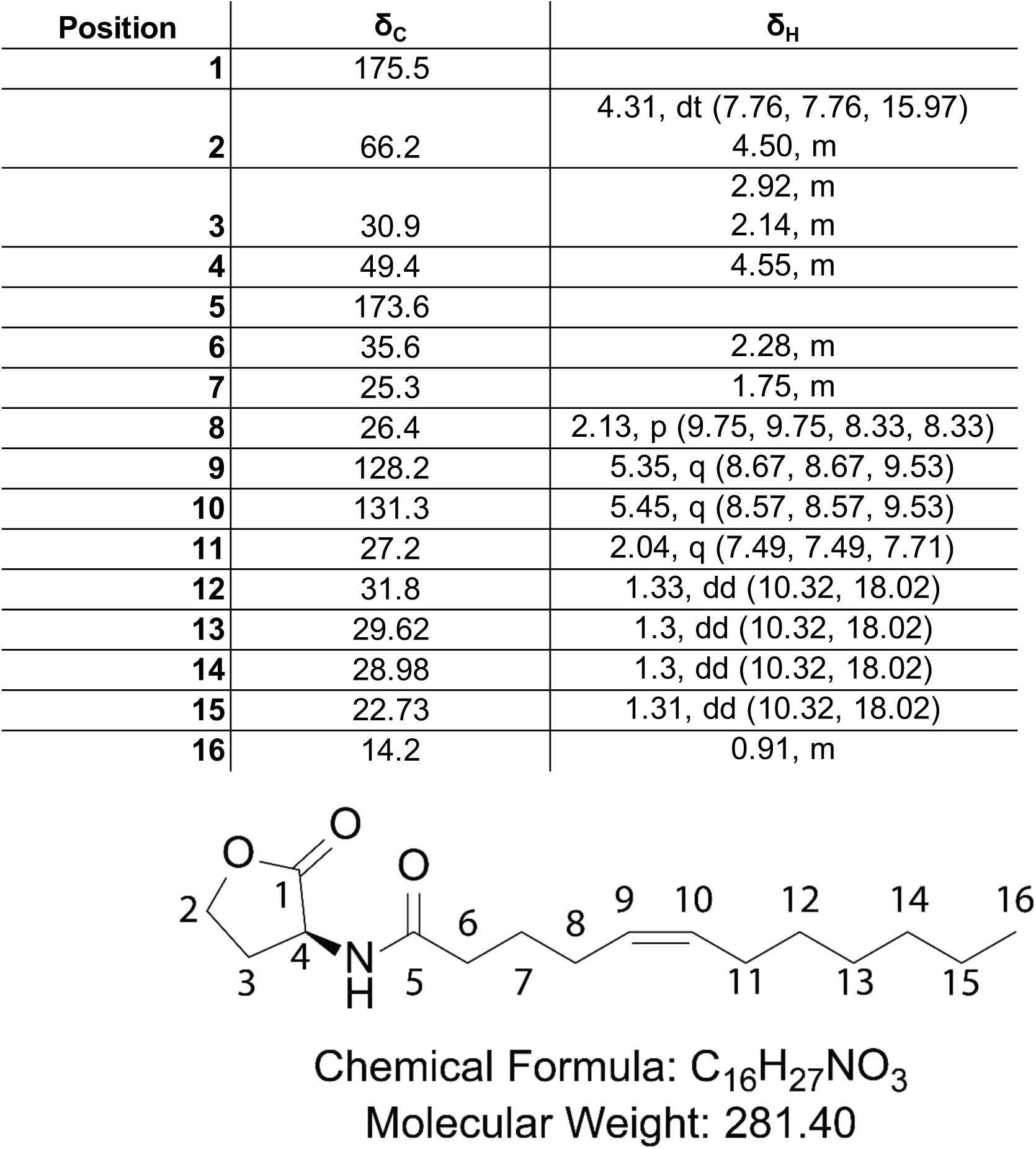
^1^H [ppm, mult. (*J* in Hz)] and ^13^C NMR data of C12:1-HSL in CDCl_3_.

**Figure S29.**
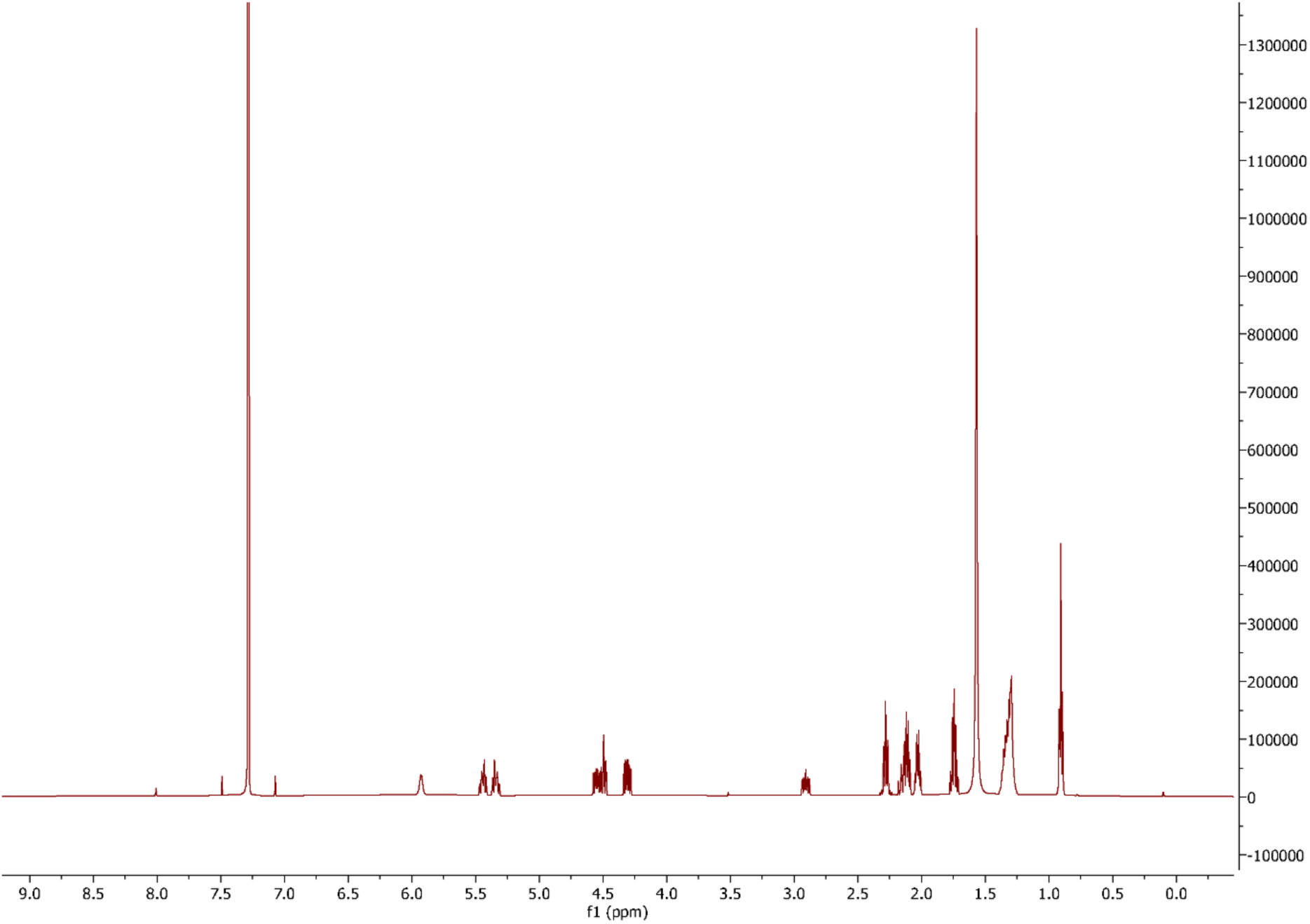
^1^H NMR spectrum of C12:1-HSL in CDCl_3_.

**Figure S30.**
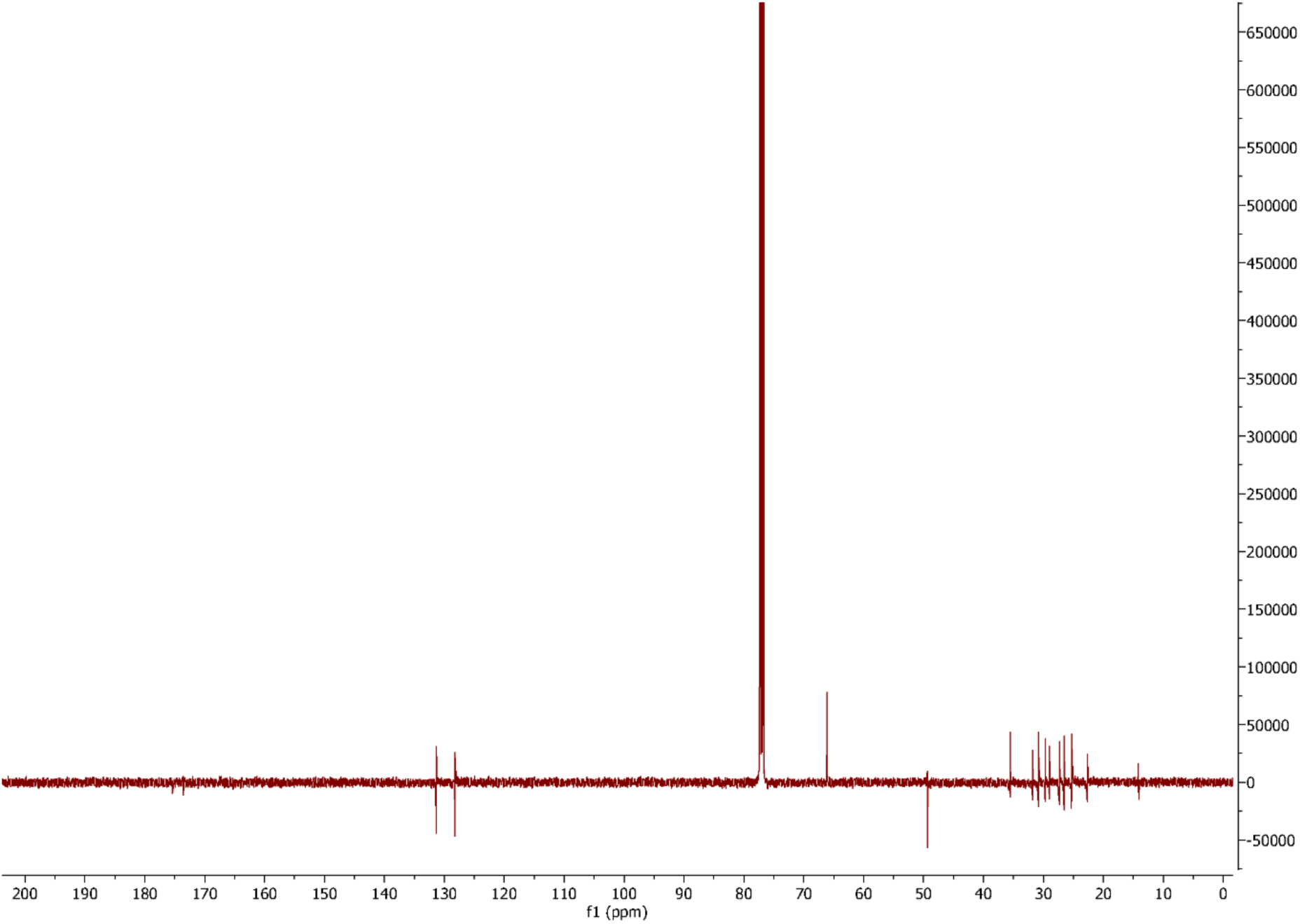
^13^C NMR spectrum of C12:1-HSL in CDCl_3_.

**Figure S31.**
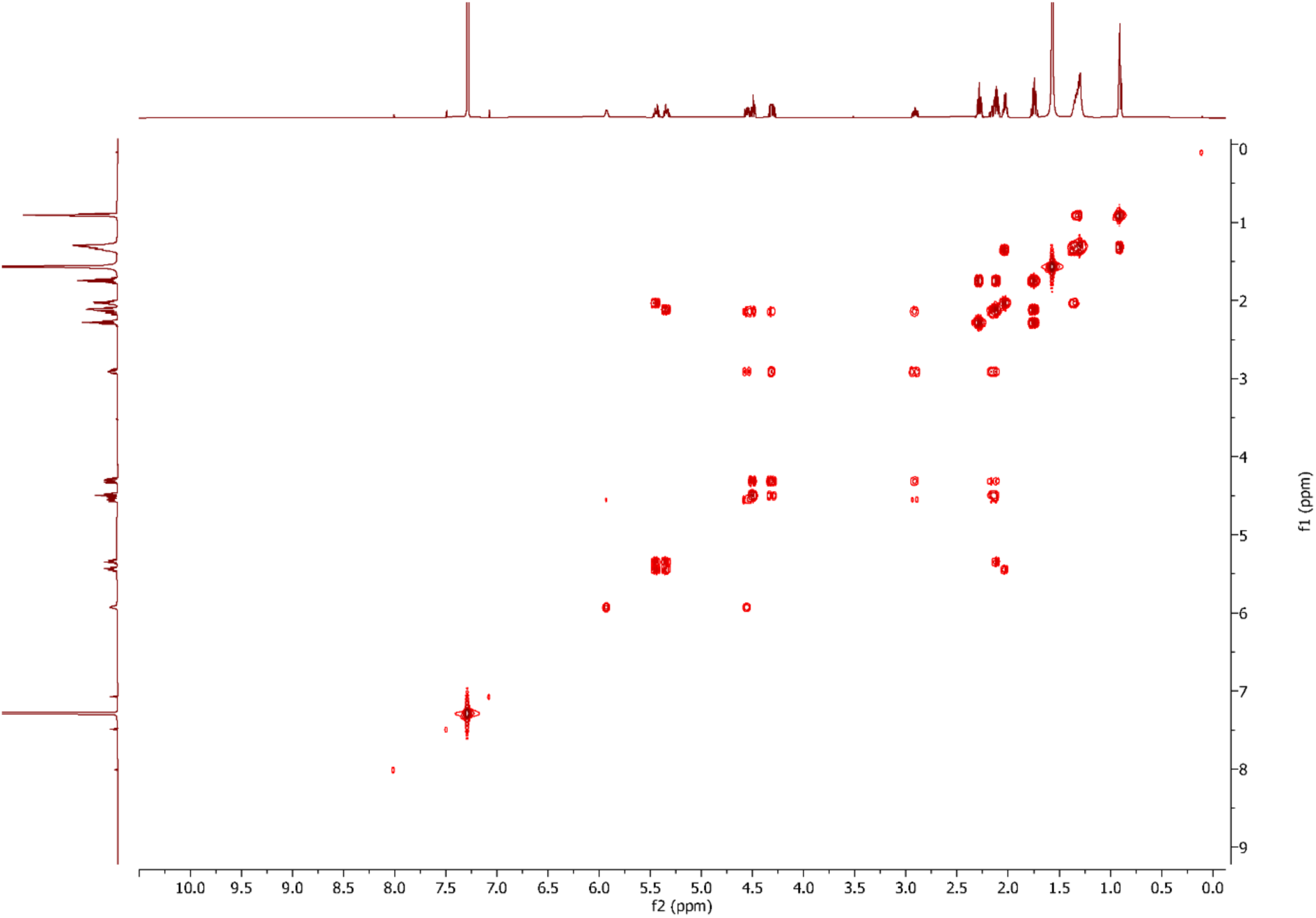
^1^H-^1^H COSY spectrum of C12:1-HSL in CDCl_3_.

**Figure S32.**
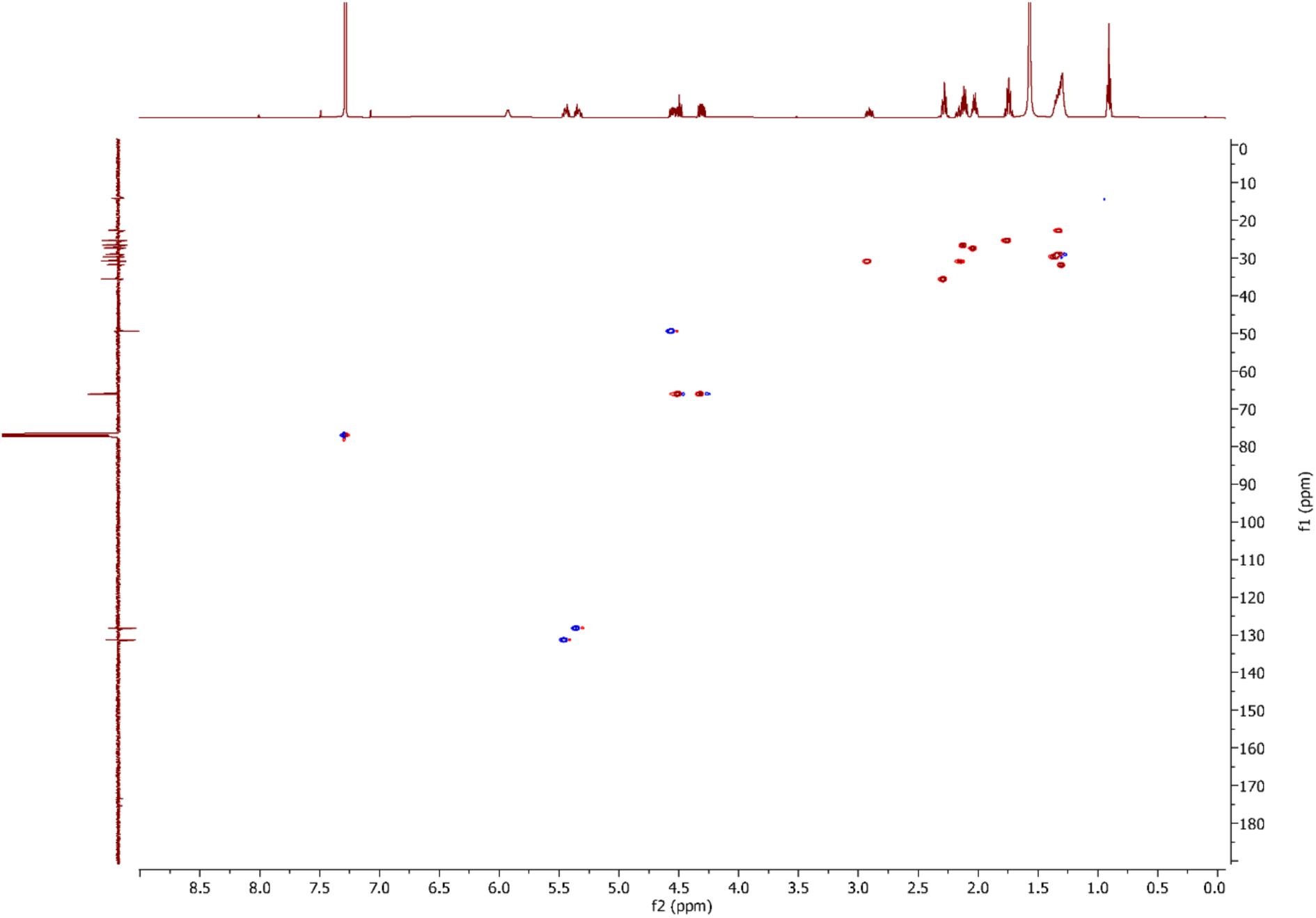
^1^H-^13^C HSQC spectrum of C12:1-HSL in CDCl_3_.

**Figure S33.**
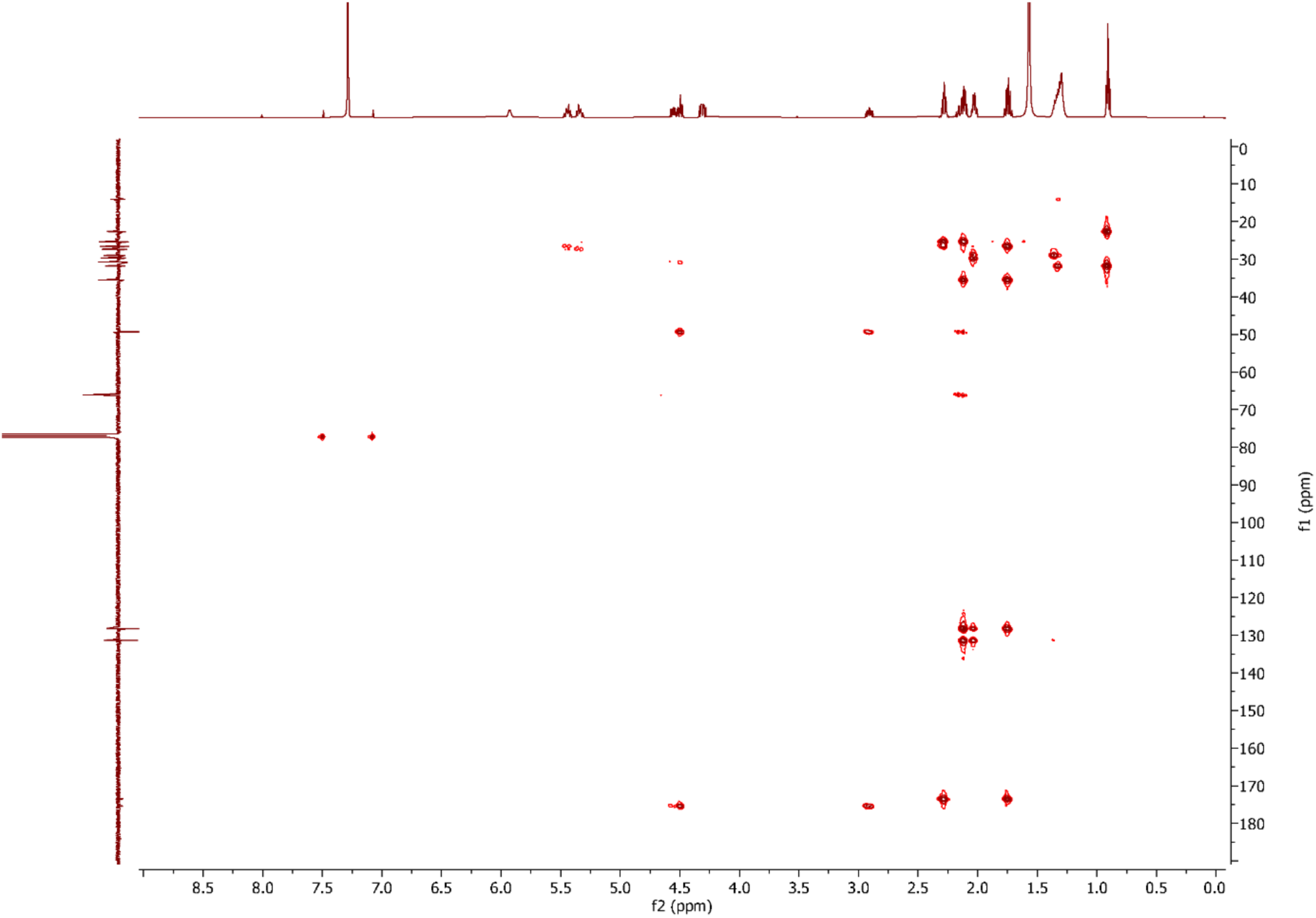
^1^H-^13^C HMBC spectrum of C12:1-HSL in CDCl_3_.

**Figure S34.**
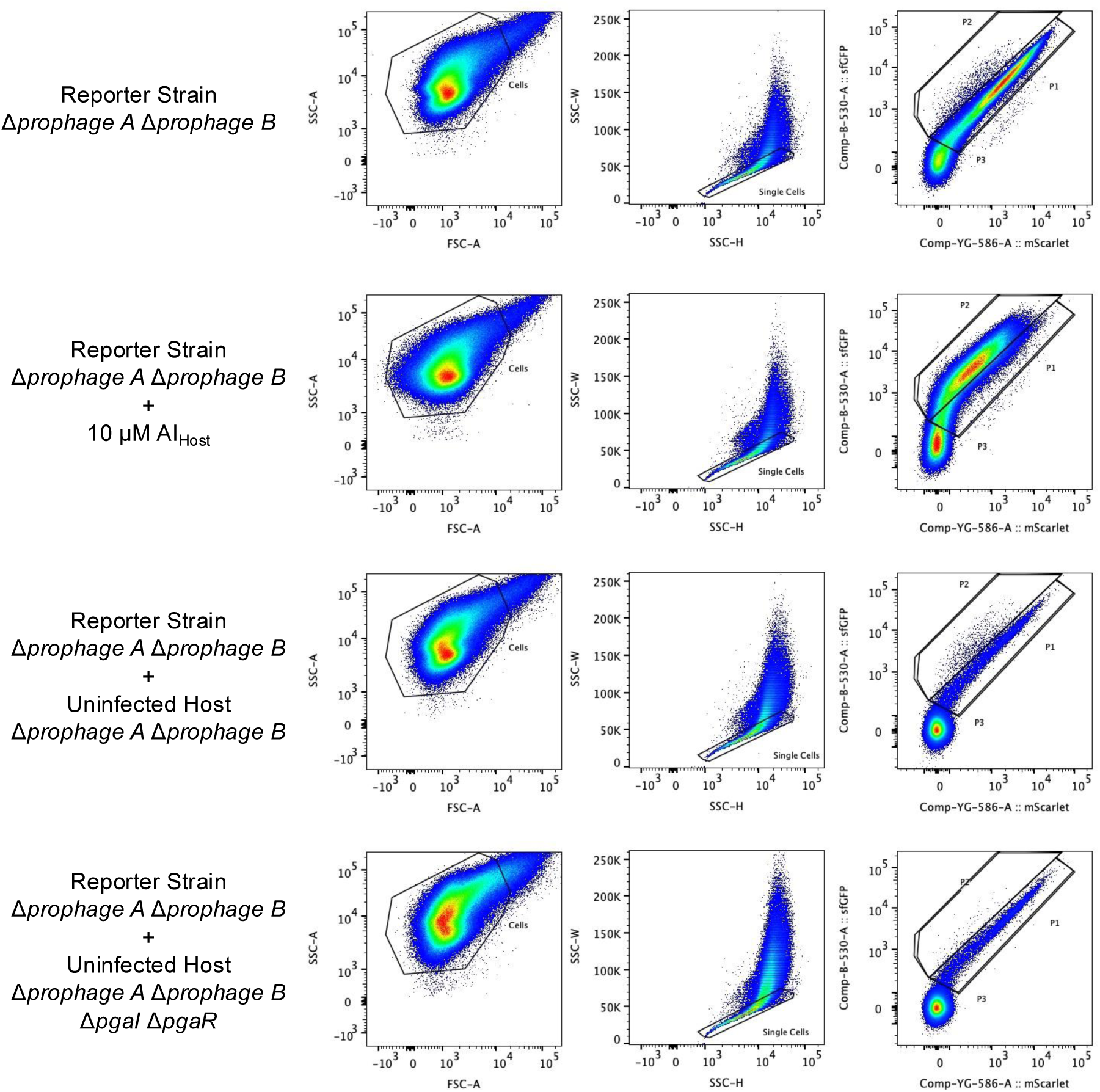
Flow cytometry gating strategy for quantifying reporter responses in co-culture experiments. Representative flow cytometry plots showing the sequential gating strategy used to quantify reporter activation. The Reporter Strain (Δ*prophage A* Δ*prophage B*) carrying the dual-fluorescence reporter was analyzed either alone, with 10 μM AI_Host_, or following co-culture with the indicated Uninfected Host strains. Cells were first gated based on forward scatter (FSC-A) and side scatter (SSC-A) (left panels), followed by gating on SSC-H vs. SSC-W (middle panels). Single cells were then displayed according to mScarlet-I fluorescence vs. sfGFP fluorescence (right panels). Reporter Strain cells were classified as non-responding (Gate P1) or responding (Gate P2) based on mScarlet-I fluorescence.

**Supplementary Table 1. Predicted Roseobacter prophages and associated QS modules.**

**Supplementary Table 2. Strains used in this study.**

**Supplementary Table 3. Oligonucleotides used in this study.**

**Supplementary Table 4. Plasmids used in this study.**

**Supplementary Table 5. List of Alphaproteobacterial genomes containing multiple *luxI* genes.**

**Supplementary File 1. Custom code used in this study.**

## Materials and Methods

### Strains and culture conditions

*P. inhibens* T5^T^ (DSM16374) and *P. inhibens* DSM17395 were obtained from the DSMZ strain collection. *P. inhibens* strains were grown in filter-sterilized (0.2 μm) (Sigma) Marine Broth (MB) (Fisher) at 28°C. *E. coli* TOP10 and TOP10 *pir*^+^ were used for cloning. The DAP auxotrophic, *pir*^+^ *E. coli* strain JKE201 was used as the donor for plasmid conjugation into *P. inhibens*. All *E. coli* strains were cultured in LB-Miller broth at 37°C. Antibiotics were used at the following concentrations in *E. coli*: ampicillin (Amp, 100 μg mL⁻¹), gentamicin (Gm, 20 μg mL⁻¹), chloramphenicol (Cm, 20 μg mL⁻¹), and kanamycin (Kan, 50 μg mL⁻¹). Gm (25 μg mL⁻¹) and Kan (50 μg mL⁻¹) were used in *P. inhibens*. Strains used in this study are listed in Supplementary Table 2.

For co-culture experiments, overnight cultures of *P. inhibens* strains were grown at 28°C to saturation with shaking. The OD₆₀₀ of each culture was measured, and a volume of cells corresponding to a 1 OD unit was pelleted by centrifugation and resuspended in 1 mL of fresh ½ MB. Co-cultures were established in 3 mL volumes of ½ MB supplemented with Gm by inoculating the medium with 5 μL of the appropriate reporter strain. Uninfected Host cells were then added at volumes of 2.5, 5, 25, 50, 100, or 150 μL to achieve Uninfected Host to Reporter Strain ratios of 0.5:1, 1:1, 5:1, 10:1, 20:1, and 30:1. Controls containing the Reporter Strain or the Uninfectd Host alone were prepared in parallel, and a positive control supplemented with 10 μM 3OH-C10-HSL was also included. Cultures were grown overnight at 28°C. The following morning, samples were prepared for flow cytometry by 1:10 dilution into ½ MB. Cells were distinguished from debris using DAPI staining (Invitrogen) which was added to a final concentration of 5 μg mL^-1^. Samples lacking DAPI were diluted into ½ MB.

For prophage induction experiments, overnight cultures of reporter strains were back-diluted 1:50 into ½ MB and grown for 2 h at 28°C. DMSO, 10 mM 3OH-C10-HSL, or 100 mM mBTL were added to produce a 1:1000 dilution of the stock. The cultures were grown for an additional 4-5 h at 28°C.

### Plasmid construction and transfer into *P. inhibens*

Plasmid constructs for gene expression in *P. inhibens* were based on pBBR1-MCS5. The suicide vector pRE112 with a counter-selectable *sacB* gene was employed to engineer *P. inhibens* chromosomal modifications. PCR with Phusion polymerase (NEB) was used to generate insert and backbone DNA fragments. DNA fragment assembly was carried out according to standard cloning techniques. Primers and plasmids are listed in Supplementary Tables 3 and 4, respectively.

For flow cytometry experiments, a dual-fluorescence reporter plasmid was constructed using a modified pBBR1-MCS5 backbone (pMLB358). The reporter plasmid harbored *sfGFP* inserted immediately downstream of a Gm resistance gene, preceded by a strong ribosome binding site. The plasmid also encoded *mScarlet-I* under the control of the Phage A P_Lysogeny_ promoter.

To conjugate plasmids into *P. inhibens*, an overnight culture of *E. coli* JKE201 carrying the plasmid of interest was grown to saturation in LB supplemented with 0.3 mM DAP and the appropriate antibiotic. The recipient *P. inhibens* strain was grown to saturation in ½ MB. Cells from 1 mL of each culture were pelleted by centrifugation. The *P. inhibens* recipient was resuspended in 1 mL of ½ MB and the *E. coli* donor was resuspended in 100 μL of ½ MB. A 50 μL aliquot of the donor suspension was combined with 5 μL of the recipient suspension and spread onto a ½ MB agar plate supplemented with 0.3 mM DAP. The plate was incubated for at least 12 h at 30°C, after which cells were recovered and plated on selective medium.

Deletion constructs were assembled by flanking a kanamycin resistance gene (Kan) with ∼1.5 kb of DNA encoding the chromosomal regions adjacent to the target gene. The DNA encoding the Kan resistance cassette and accompanying tandem FTR sites was amplified from the pUC18R6K-mini-Tn*7* plasmid. Chromosomal DNA from *P. inhibens* T5^T^ was used as the PCR template. Suicide vectors were introduced into *P. inhibens* strains by conjugation followed by inoculation into 5 mL of ½ MB supplemented with Kan. Cultures were grown to saturation to select merodiploids. To obtain double-crossover events, 100 μL of the saturated culture was transferred to 5 mL of ½ MB supplemented with 10% sucrose and Kan, and the culture was again grown to saturation. A 10 μL aliquot of this culture was plated to single colonies on ½ MB agar supplemented with 10% sucrose and Kan. Colonies were screened by PCR to confirm deletions. To eliminate the Kan resistance cassette, plasmid pMLB320 was introduced by conjugation. pMLB320 harbors the gene encoding the FLP recombinase from pCP20 under control of the λ cI(TS) promoter and it also carries *sacB*. To induce *FLP* expression, exconjugants were back-diluted into ½ MB and grown at 42°C for 30 min, followed by growth to saturation at 28°C. Cells were plated on ½ MB agar supplemented with 10% sucrose, and individual colonies were screened by PCR to identify those that had lost the Kan resistance cassette.

### Isolation and identification of host and phage autoinducers

Plasmids overexpressing host and prophage *luxI* genes were constructed by cloning each *luxI* gene under the *Pc* promoter into pBBR1-MCS5. The plasmids were individually introduced into *P. inhibens* DSM17395. To identify the autoinducers produced by the LuxI enzymes, strains were inoculated into a 100 mL culture of ½ MB supplemented with Gm and incubated for three days at 28°C. The cultures were extracted with ethyl acetate (Sigma) at 1:1 (v/v). The organic layer was isolated and dried *in vacuo* (Genevac HT6 S3i Evaporator). The dried extract was dissolved in 50 μL of MeOH and injected onto an Agilent Eclipse C18 1.8 μm (2.1 x 50 mm) column, using an Agilent MSD iQ LC-MS system coupled to an Agilent 1290 Infinity II HPLC. The mobile phase was a water-acetonitrile (MeCN, Sigma) gradient containing 0.1% formic acid. The flow rate was 0.3 mL min^-1^. Chromatography was performed as follows: 0-2 min 5% MeCN, 2-10 min 5-95% MeCN, 10-12 min 95% MeCN. The LC-MS iQ was carried out in positive mode scanning between m/z 100-1000 with the following source parameters: gas temperature 325°C, gas flow 11 (L min^-1^), capillary voltage 3500 V. Extracted ion chromatograms for ions of interest were quantified using area under the curve (Fig. S12).

To purify each autoinducer, a 10 mL aliquot of a saturated culture of each strain was distributed across ten 2 L flasks containing 1 L of ½ MB supplemented with Gm and 20 g of Amberlite XAD-16 resin (ThermoFisher). The Amberlite XAD-16 resin was conditioned prior to use as follows: beads were subjected to autoclave followed by soaking in excess sterile water (∼1% w/v) for 30 min. The water was removed by aspiration. The beads were washed twice with excess MeOH (30 min per wash), with decanting between washes. Residual MeOH was removed by repeated washes with sterile water and aspiration. Beads thus conditioned were resuspended in ½ MB supplemented with Gm and distributed to flasks immediately prior to inoculation with cells. The cultures containing the beads were incubated for 3 days with agitation at 28°C. The beads were collected in a chromatography column fitted to a vacuum manifold. The resin was washed with water to remove residual culture medium. Autoinducers were eluted into 10 column volumes of dichloromethane (Sigma). The eluate was dried over anhydrous MgSO_4_ (Sigma). The dichloromethane was evaporated *in vacuo*.

Fractionation of the above autoinducer preparations was performed on an Agilent Bond Elut C18 Cartridge, 10 g bed mass, 60 mL column size. The column cartridge was first conditioned with 10 mL MeCN followed by 10 mL water. The crude extract was dissolved in 2 mL of MeOH, applied to the column, and eluted with a stepwise gradient consisting of 10 mL fractions of 5%, 20%, 40%, 60%, 80% MeCN/water, followed by a final wash of 40 mL of 100% MeCN. All fractions were dried *in vacuo* and monitored for the ion of interest (C12:2-HSL m/z 280.2; C12:1-HSL m/z 282.2; and 3-OH-C12:1-HSL m/z 298.2) using an Agilent MSD iQ System coupled to an Agilent 1290 Infinity II HPLC. For this analysis, dried fractions were dissolved in 1 mL MeOH, diluted 1:10 in MeOH, and 2 µL were injected onto an Agilent Eclipse C18 1.8 μm (2.1 x 50 mm) column and chromatography was carried out identically to that described above. Regarding the phage B C12:2-HSL, the 60% and 80% MeCN fractions contained the ion of interest. In the case of the phage A C12:1-HSL, the 60% and 80% MeCN fractions contained C12:1-HSL along with a minor amount of C12:2-HSL, and the 80% and 100% MeCN fractions contained 3-OH-C12:1-HSL.

Fractions containing the ion(s) of interest were pooled, dried *in vacuo*, dissolved in 1 mL of MeOH, and further purified on an Agilent 1290 Infinity II HPLC coupled to an Agilent 1260 Infinity II Analytical Fraction Collector. Samples were injected iteratively in 20 µL volumes onto a Phenomenex Luna 10 μm C18 (250 x 10 mm) column. The mobile phase consisted of a water-MeCN gradient containing 0.1% formic acid. The flow rate was 4.5 mL min^-1^. Chromatography was performed as follows: 0-5 min 5-50% MeCN, 5-25 min 50-65% MeCN, 25-27 min 65-95% MeCN, 27-30 min 95% MeCN. Fraction collection was triggered by monitoring the 230 nm wavelength. For C12:1-HSL, an additional separation step was undertaken to resolve it from the co-eluting C12:2-HSL using a modified gradient: 0-5 min 5-51% MeCN, 5-15 min 51% MeCN, 15-17 min 51-95% MeCN, 17-20 min 95% MeCN. The sample was iteratively injected until the material was pure. The final yields were: Phage A C12:1-HSL: 3.7 mg; 3-OH-C12:1-HSL: 7.4 mg and a minor product, C12:2-HSL: 2.5 mg. The Phage B yield was C12:2-HSL: 1.2 mg.

### Olefin cross metathesis for HSLs

Approximately 0.2 mg of each pure, isolated HSL was dissolved in 455 μL of dichloromethane, 45 μL of methyl acrylate (Sigma-Aldrich), and 50 μg of 2nd generation Hoveyda-Grubbs catalyst (Sigma-Aldrich) (Fig. S13-S15). The reactions were stirred for 3 h at room temperature. A 50 μL aliquot of the reaction mixture was concentrated *in vacuo*, redissolved in 100 μL of MeOH, and analyzed by HRMS (C12:2-HSL) or iQMS (3-OH-C12:1-HSL and C12:1-HSL).

### HRMS and NMR procedures

HRMS was acquired on an Agilent 6546 LC-QTOF 1290 LC system using an Agilent Eclipse C18 1.8 μm (2.1 x 50 mm) column. The mobile phase was a water-MeCN gradient containing 0.1% formic acid. The flow rate was 0.3 mL min^-1^. The injection volume was 2 µL. Chromatography was performed as follows: 0-2 min 5% MeCN, 2-10 min 5-95% MeCN, 10-12 min 95% MeCN. Source parameters were as follows: gas temperature 275°C, gas flow 12 (L min^-1^), capillary voltage 3500 V. MS1 acquisition was carried out in positive mode scanning from 100-1000 m/z. Data were processed using Agilent MassHunter Qualitative Analysis 10.0. Peaks were extracted by m/z within a 10 ppm error window to generate extracted ion chromatogram (EIC) spectra.

NMR spectral data were acquired at the Princeton University Department of Chemistry NMR Facility. 1D (^1^H, ^13^C) and 2D (COSY, HSQC, and HMBC) NMR spectra were collected using a Bruker Avance III HD 800-MHz NMR spectrometer equipped with a triple resonance cryoprobe. ^1^H NMR spectra were tabulated as follows: chemical shift, multiplicity (s = singlet, d = doublet, t = triplet, dd = doublet of doublets, ddd = doublet of doublets of doublets, dt = doublet of triplets, m = multiplet), coupling constant (Hz), and number of protons. ^13^C NMR spectra were tabulated by observed peak, and no special nomenclature is used for equivalent carbons. For C12:1-HSL and 3-OH-C12:1-HSL, NMR data were collected using a Bruker 500 MHz NMR spectrometer equipped with a DCH double resonance cryoprobe. All NMR data were analyzed with MestReNova software, and the chemical shifts were recorded as δ values (ppm) referenced to solvent residual signals. Spectra are provided in Figs. S16-S33.

### Quantitation of HSLs produced by *P. inhibens* T5^T^

Cultures of *P. inhibens* T5^T^ and DSM 17395 were grown ½ MB for 16 h at 28°C, with shaking at 200 rpm before 3 mL aliquots were supplemented with an internal standard (C10-HSL, 2.5 μM) and extracted with two volumes of ethyl acetate, twice. The organic phase was removed and dried *in vacuo*. The dried extracts were resuspended in 200 μL of MeOH (Supelco) and analyzed by UPLC-MS (Agilent Technologies 1290 Infinity II UPLC system) coupled to an Agilent Technologies iQ single-quadrupole mass spectrometer. Separation was performed on an Agilent Eclipse C18 1.8 μm (2.1 x 50 mm) column using methods identical to those used above.

### Reporter assays

Overnight cultures of reporter strains were back-diluted 1:50 into ½ MB supplemented with appropriate antibiotics and grown for 2 h at 28°C. Autoinducer stocks (1000x in DMSO) were diluted into ½ MB to prepare 2x working solutions. 200 μL of the working stocks or of a DMSO control were combined with 200 μL of the back-diluted culture in wells of a 96-well deep well plate (USA Scientific). Plates were incubated for 14 h at 28°C. Strains containing the P*_paaZ2_*-*mScarlet-I* reporter, were grown 42 h at 28°C to compensate for slower growth compared to other strains. Subsequently, the cultures were mixed by repeatedly drawing up and down with a pipetman followed by transfer to a 96-well flat clear-bottom black polystyrene microplate (Corning Costar). OD₆₀₀ and mScarlet fluorescence (excitation 569 nm, emission 594 nm) were measured using a BioTek Synergy Neo2 plate reader. Relative *mScarlet-I* expression was calculated by dividing the mScarlet-I fluorescence reading by the OD₆₀₀ measurement for each well.

### Microscopy

Overnight cultures of *P. inhibens* strains were back-diluted 1:50 into fresh ½ MB medium supplemented with appropriate antibiotics and grown for 2–3 h at 28°C. Cultures were gently mixed using a pipette to break up aggregates. A 2 μL aliquot of each culture was dispensed onto a 24×60 mm cover glass no. 1.5 (Fisherbrand). A 1.5% agarose pad (made using fresh ½ MB medium supplemented with 0.1% DMSO, 10 μM 3OH-C10-HSL, and/or 10 μM 3OH-C12:1-HSL) was placed on top of the droplet, and a 22×22 mm cover glass no. 1.5 (Globe Scientific) was placed on the agarose pad^32^. The sample was incubated at 30°C using a stage-top incubator (Tokai Hit) and imaging was performed using an Eclipse Ti2 inverted confocal microscope (Nikon) equipped with a CSU-W1 SoRa scanning unit (Yokogawa), an ORCA-Fusion BT sCMOS camera (Hamamatsu Photonics), and a CFI Plan Apo DM Lambda oil-immersion objective lens (100×, NA 1.45, Nikon). Isolated cells were identified at different xy-positions, and phase-contrast images of the same set of xy-positions were acquired at a frequency of once every 10 min for 12 h. To account for focus drift during imaging, at each time point, for each xy-position, a z-stack of multiple focal planes (at least 300 nm apart, centering on the best-focused z-slice and spanning at least 3 μm) was acquired. Image acquisition was controlled using NIS-Elements Advanced Research software (Nikon). Image processing and preparation of the movies were performed using NIS-Elements Advanced Research and ImageJ2/FIJI software^33^.

### qRT-PCR and qPCR

Cells used to prepare RNA were collected by centrifugation and flash frozen in liquid nitrogen. RNA was extracted and DNA was removed using the RNeasy Mini Kit (Qiagen). cDNA was produced using SuperScript VILO MasterMix (Thermo Fisher). DNA was extracted from identically collected and flash frozen cells using the MasterPureTM (Epicenter) kit followed by resuspension in ddH_2_O to a concentration of 100 ng μL^-1^. To isolate phage particles, cell-free culture fluids were clarified by pelleting followed by passage of the recovered supernatants through Spin-X centrifuge tube filters (Corning). These samples were subjected to treatment with the Promega DNaseI kit. DNA and cDNA were quantified using PerfeCTa SYBR Green FastMix Low ROX (Quanta BioSciences). Relative quantity represents ΔΔCt compared to the reference, 16S RNA, and DMSO treatment.

### Flow Cytometry

Flow cytometry data were acquired using a FACSymphony A3 flow cytometer (BD Biosciences, San Jose, CA) with a 488 nm laser for GFP excitation and a 561 nm laser for mScarlet excitation. Emitted fluorescence was captured through a 530/30 nm and a 586/15 nm bandpass filter for GFP and mScarlet, respectively. Cells were gated on FCS-A vs. SSC-A and SSC-H vs. SSC-W to identify single cells. Fluorescent populations were identified on an mScarlet YG-586-A vs. sfGFP B-530-A plot (Fig. S34). The abundance of Uninfected Host cells was determined by dividing the number of GFP-negative cells by the total number of cells. The percentage of responding Reporter Strain cells was determined by dividing the number of cells within the P2 fluorescence gate by the total number of cells within both the P1 and P2 fluorescence gates (Fig. S34).

### Identification of Alphaproteobacteria strains with multiple *luxI* genes and LuxI protein clustering

These and the following computational analyses, unless stated otherwise, were performed with custom Python (v3.14.2) or R (v.4.5.2) scripts. All complete RefSeq genomes corresponding to Alphaproteobacteria (Taxonomy ID: 28211) were obtained from the NCBI nt database with Bio.Entrez^34^. Genomes harboring *luxI* genes were identified with a profile HMM search with HMMER3 (v.3.3.2) for domain PF00765 (Pfam v.37) against a database consisting of all proteins from the queried genomes^35,36^. Any genome with two or more LuxI hits was determined to harbor multiple *luxI* genes (multi-*luxI*). LuxI proteins from multi-*luxI* genomes were extracted and clustered using CD-HIT (v.4.8.1) with an 80% amino acid identity cutoff (parameters: “-c 0.80 -n 5”)^37^.

### Core genome phylogeny and identification of horizontally-transferred *luxI* genes

Genbank files for all multi-*luxI* Alphaproteobacteria genomes were retrieved by submitting their respective RefSeq Accession IDs (Supplementary Table 5) to NCBI Datasets^38^. A set of core proteins (those encoded in 100% of the input genomes) present across the 523 Alphaproteobacteria genomes was extracted from the Genbank files using usearch with an amino acid identity cutoff of 30%, as implemented by the BPGA pipeline (v1.3), and aligned using MUSCLE^39,40,41^. Five genomes were excluded due to incomplete file formatting. Due to the high level of species diversity in the dataset, only 22 core proteins were identified. The resulting alignment of the core proteins, corresponding to 7,194 amino acid positions, was used to construct a phylogenetic tree with RAxML-NG (v.2.0.0) (parameters: “--model LG+I+G4 --bs-trees autoMRE --bs-metric tbe --seed 52143”)^42^. Branch support was calculated using 500 bootstrap replicates. The substitution model was determined with ModelTest-NG (v.0.1.7)^43^. LuxI protein clusters were appended to their corresponding genomes in the core genome tree. LuxI protein clusters that did not group according to the core genome phylogeny were considered horizontally transferred. All LuxI protein clusters displaying a pattern of vertical transmission, representing orphan *luxI* genes, representing known *traI/R* genes, or representing known *rhiABC* genes were excluded from our analyses. Phylogenetic trees with associated LuxI protein clusters were generated with the Interactive Tree of Life (iTOL v.7.6) web server^44^.

### Phage identification, phage LuxI typing, viral taxonomy, and alignments

A 200 kb window around each horizontally-transferred *luxI* gene in its respective genome was extracted and submitted to VIBRANT (v.1.2.0) to predict potential prophage sequences^22^. Identified prophages were manually curated to determine which prophage-borne *luxI* genes resided in *xre-luxR-luxI* operons. Roseobacter clade prophages harboring *xre-luxR-luxI* operons were extracted and upstream and downstream flanking host genes were identified. The corresponding regions of these upstream to downstream genes in prophage-naïve genomes were used to search NCBI nt and a longitudinal *Phaeobacter* dataset for prophages that our analysis may have missed due to the strictness of our parameters^45^. LuxI proteins from all extracted prophages, including these newly identified prophages, were clustered with CD-HIT (v.4.8.1)^37^. The resulting 10 clusters were assigned a type from A to J, with A and B corresponding to the Phage A and Phage B *xre-luxR-luxI* operons, respectively. For visualization, protein alignments were performed in Geneious Prime (v.2026.0.2) using MUSCLE^41^. Heatmaps were generated in R with pheatmap (v.1.0.13) using percent identities calculated in Geneious Prime. The acquired prophage genomes from the Roseobacter clade were dereplicated with CD-HIT-EST (v.4.8.1) (parameters: “-c 0.99 - aS 1.0 -g 1 -d 0”)^37^. The unique prophage genomes were submitted to vConTACT3 (v.3.2.4) with the Prokaryotic Viral RefSeq database (v.211) to determine phage taxonomy^23,46^. Genome synteny was calculated and visualized with clinker (v.0.0.31)^47^.

