## Supplementary File 1 for "Competing Inter-Domain Quorum-Sensing Systems Control Prophage Lysis-Lysogeny Decisions": 1_LuxI_HMMsearch.py

```
#!/usr/bin/env python3
'''
Created on December 20, 2025

@author: Francis J Santoriello (Polished with Claude Opus 4.7 on June 30, 2026)

This script accepts an NCBI Taxonomy ID and HMM Profiles. It then:
    1. Retrieves all RefSeq genomes associated with the input Taxonomy ID in GenBank format.
    2. Extracts all CDS features from each genome and writes them to a combined protein
       fasta file.
    3. Runs hmmsearch for every HMM profile against the custom protein database.
    4. Assesses protein co-occurrence per assembly.

This script returns:
    1. all_prots.faa             -- combined CDS translations for every RefSeq assembly fetched
    2. <HMM>_hits                -- hmmsearch tabular output
    3. <HMM>_hits.aln            -- hmmsearch Stockholm alignment of hits
    4. <HMM>_hmm.faa             -- FASTA of hmmsearch hits (with alignment gaps)
    5. protein_co-occurence.csv  -- assembly x HMM hit-count matrix
    6. multi-hit_strains.csv     -- assemblies with >=2 LuxI hits
    7. LuxI_multi.faa            -- LuxI proteins from multi-hit assemblies
'''

import os
import argparse
import textwrap
import subprocess
import glob
import pandas as pd
from Bio import Entrez
from Bio import SeqIO

cwd = os.getcwd()  # current working directory


### User Input ###
class MyFormatter(argparse.ArgumentDefaultsHelpFormatter,
                  argparse.RawDescriptionHelpFormatter):
    pass

parser = argparse.ArgumentParser(
                prog='1_LuxI_HMMsearch.py',
                formatter_class=MyFormatter,
                description=textwrap.dedent('''\
                    This script accepts an NCBI Taxonomy ID and HMM Profiles. It then:
                        1. Retrieves all RefSeq genomes associated with the input Taxonomy ID in GenBank format.
                        2. Extracts all CDS features from each genome and writes them to a combined protein
                           fasta file.
                        3. Runs hmmsearch for every HMM profile against the custom protein database.
                        4. Assesses protein co-occurrence per assembly.

                    This script returns:
                        1. all_prots.faa             -- combined CDS translations for every RefSeq assembly fetched
                        2. <HMM>_hits                -- hmmsearch tabular output
                        3. <HMM>_hits.aln            -- hmmsearch Stockholm alignment of hits
                        4. <HMM>_hmm.faa             -- FASTA of hmmsearch hits (with alignment gaps)
                        5. protein_co-occurence.csv  -- assembly x HMM hit-count matrix
                        6. multi-hit_strains.csv     -- assemblies with >=2 LuxI hits
                        7. LuxI_multi.faa            -- LuxI proteins from multi-hit assemblies
                    '''),
                usage='%(prog)s [options] --ncbi_email <email> <taxonomy_id> <HMM_profiles>\n\n '
                      'Dependencies: python 3, biopython, EntrezDirect, hmmer, cd-hit, muscle\n')
parser.add_argument('taxonomy_id',
                    help='NCBI Taxonomy Browser ID for fetching genomes. Ex: txidXXXXX')
parser.add_argument('hmms', nargs='*',
                    help='HMM profile files for co-occurence matrix')
parser.add_argument('--ncbi_email', type=str, required=True,
                    help='Contact email for NCBI E-utilities (required by NCBI policy).')
parser.add_argument('--ncbi_api_key', type=str, default=None,
                    help='NCBI E-utilities API key (optional; raises per-second rate limit).')
parser.add_argument('--assembly_level', type=str, default='complete genome',
                    help='NCBI assembly level filter (e.g. "complete genome", "chromosome")')
parser.add_argument('-e', '--hmm_eval', type=float, default=0.00001,
                    help='E-value threshold for hmmsearch')
parser.add_argument('-o', '--output', type=str, default='outdir',
                    help='Name of directory for output files')

args = parser.parse_args()


### Variables ###
taxonomy = args.taxonomy_id
assembly = args.assembly_level
hmm_profiles = args.hmms
threshold = args.hmm_eval
outdir = args.output

# NCBI credentials come from the CLI so nothing identifying is checked into source.
#   --ncbi_email   required by NCBI's E-utilities policy
#   --ncbi_api_key optional; raises the per-second rate limit
Entrez.email = args.ncbi_email
if args.ncbi_api_key:
    Entrez.api_key = args.ncbi_api_key


def search_and_fetch_genomes(tax_id, assmbl_lvl, outpath):
    '''
    Fetch every RefSeq nucleotide accession under `tax_id` matching `assmbl_lvl`,
    write each as a per-accession .gb in `outpath`, and append every usable CDS
    translation to `outpath/all_prots.faa`.

    "Usable" CDS = has protein_id, product, translation (without '?'), and the
    record has an Assembly: dbxref so we can tag the FASTA header with the GCF id.
    '''
    search_term = f'{tax_id}[ORGN] AND {assmbl_lvl}[ALL] AND refseq[filter]'
    print(f"Searching assembly for: {search_term}")

    # ESearch returns UIDs for the matching nucleotide records.
    # retmax is set high so we get them all in one call.
    handle = Entrez.esearch(db='nucleotide', term=search_term, retmax=1000000)
    record = Entrez.read(handle)
    handle.close()

    nuc_ids = record["IdList"]
    count = len(nuc_ids)
    print(f"Found {count} accessions.")

    if count == 0:
        print("No records found. Exiting.")
        return

    print(f'Retrieved {len(set(nuc_ids))} unique nucleotide accessions.')

    # Resume support: any *.gb already in outpath was fetched on a prior run --
    # skip those records when we see them come back from efetch.
    fetched_gbs = glob.glob(os.path.join(outpath, "*.gb"))
    fetched = {os.path.splitext(os.path.basename(p))[0] for p in fetched_gbs}

    with open(os.path.join(outpath, "all_prots.faa"), 'w') as outfile:
        # Efetch is happier with smaller batches; 200 is a safe chunk size.
        chunked_nuc_ids = [nuc_ids[i:i + 200] for i in range(0, len(nuc_ids), 200)]

        for i, chunk in enumerate(chunked_nuc_ids):
            handle = Entrez.efetch(id=chunk, db='nuccore', rettype='gbwithparts',
                                   retmode='text')
            counter = 0
            total_seqs = len(chunk)
            print(f'Total sequences in Chunk {i+1}: {total_seqs}')

            for record in SeqIO.parse(handle, 'genbank'):
                # Skip non-RefSeq (e.g. INSDC mirrors) and anything already fetched.
                if 'RefSeq' not in record.annotations['keywords']:
                    print('Non-RefSeq sequence passed.')
                    total_seqs -= 1
                    continue
                if record.id in fetched:
                    print(f"{record.id} fetched in a previous run. Removed.")
                    total_seqs -= 1
                    continue

                # Persist the per-accession GenBank for resumability / downstream use.
                with open(os.path.join(outpath, f"{record.id}.gb"), 'w') as out_gb:
                    SeqIO.write(record, out_gb, 'genbank')

                # Pull the Assembly (GCF) dbxref so we can tag every CDS header
                # with its source assembly -- needed by prot_cooccur().
                dblinks = record.dbxrefs
                assembly_link = next((s for s in dblinks if s.startswith('Assembly:')), None)
                if assembly_link is None:
                    print(f"{record.id} passed for missing Assembly db link.")
                    total_seqs -= 1
                    continue
                assembly_id = assembly_link[len('Assembly:'):]

                feat_total = 0
                feat_count = 0
                for feat in record.features:
                    feat_total += 1
                    if feat.type != 'CDS':
                        continue
                    q = feat.qualifiers
                    # Require protein_id, product, and a clean translation
                    # ('?' indicates ambiguous/unknown residues we don't want).
                    if ('protein_id' not in q
                            or 'product' not in q
                            or 'translation' not in q
                            or '?' in q['translation'][0]):
                        continue

                    header = (f">{q['protein_id'][0]} "
                              f"{q['product'][0]} "
                              f"[{record.description}|{assembly_id}]")
                    protein_seq = q['translation'][0]
                    outfile.write(f"{header}\n{protein_seq}\n")
                    feat_count += 1

                counter += 1
                total_seqs -= 1
                print(f"Chunk {i+1}: Appended {feat_count} CDS of {feat_total} "
                      f"features from accession {counter} out of {total_seqs} "
                      f"remaining accessions.")

        print('Protein fasta file complete.')


def prot_cooccur(hmmsearch_faas, outpath):
    '''
    Build an assembly x HMM count matrix.

    For every record in all_prots.faa, extract the GCF assembly id and strain
    name from the header. For every hmmsearch-hit FASTA in `hmmsearch_faas`,
    count how many hits each assembly produced. Write the matrix to
    `outpath/protein_co-occurence.csv`.
    '''
    # asmbl_dict: GCF_id -> {"Strain": name, "<hmm>": count, ...}
    asmbl_dict = {}
    with open(os.path.join(outpath, "all_prots.faa"), 'r') as prot_db:
        for record in SeqIO.parse(prot_db, 'fasta'):
            # Header format: ">prot_id product [<description>|<GCF_id>]"
            asmbl_start = record.description.rfind('GCF')
            asmbl_end = record.description.rfind(']')
            asmbl_id = record.description[asmbl_start:asmbl_end]

            strain_start = record.description.rfind('[')
            strain_end = record.description.rfind('|')
            strain_name = record.description[strain_start + 1:strain_end]

            asmbl_dict[asmbl_id] = {"Strain": strain_name}

    # hmm_dict: hmm_short_name -> [assembly_ids hit by this HMM, with repeats]
    hmm_dict = {}
    for faa in hmmsearch_faas:
        # Derive a short hmm name from the filename (strip dir + extension).
        hmm = os.path.splitext(os.path.basename(faa))[0]
        hmm_dict[hmm] = []
        with open(faa, 'r') as hmm_hits:
            for record in SeqIO.parse(hmm_hits, 'fasta'):
                asmbl_start = record.description.find('GCF')
                asmbl_end = record.description.rfind(']')
                asmbl_id = record.description[asmbl_start:asmbl_end]
                hmm_dict[hmm].append(asmbl_id)

    # Fill in per-HMM hit counts for every assembly (0 if no hits).
    for asmbl in asmbl_dict:
        for hmm, hits in hmm_dict.items():
            asmbl_dict[asmbl][hmm] = hits.count(asmbl)

    prot_mat = pd.DataFrame.from_dict(asmbl_dict, orient='index')
    prot_mat.to_csv(os.path.join(outpath, "protein_co-occurence.csv"))


### Main ###

if __name__ == '__main__':

    # Create output directory (no-op if it already exists)
    os.makedirs(outdir, exist_ok=True)

    # Fetch genomes from NCBI and build the combined protein FASTA
    search_and_fetch_genomes(taxonomy, assembly, outdir)

    # Run hmmsearch for each supplied HMM profile against all_prots.faa.
    # argv-list calls (no shell) so paths/thresholds can't be reinterpreted as shell syntax.
    print(hmm_profiles)
    for hmm in hmm_profiles:
        hmm_short = os.path.splitext(os.path.basename(hmm))[0]
        hits_table = os.path.join(outdir, f"{hmm_short}_hits")
        hits_aln = os.path.join(outdir, f"{hmm_short}_hits.aln")
        hits_faa = os.path.join(outdir, f"{hmm_short}_hmm.faa")

        print('Running hmmsearch...')
        subprocess.run(
            ["hmmsearch",
             "--tblout", hits_table,
             "-A", hits_aln,
             "-E", str(threshold),
             hmm, os.path.join(outdir, "all_prots.faa")],
            check=True,
        )

        # Convert the Stockholm alignment of hits to a plain FASTA.
        # `esl-reformat` writes to stdout; redirect to the output file via stdout=.
        with open(hits_faa, "w") as faa_out:
            subprocess.run(
                ["esl-reformat", "fasta", hits_aln],
                stdout=faa_out,
                check=True,
            )

    # Build the assembly x HMM co-occurrence matrix
    hmm_hit_faas = glob.glob(os.path.join(outdir, "*_hmm.faa"))
    print('Calculating protein co-occurence...')
    prot_cooccur(hmm_hit_faas, outdir)

    # --- Hardcoded LuxI-specific multi-hit extraction ---
    # Filter the co-occurrence table for assemblies with >=2 LuxI HMM hits
    hit_df = pd.read_csv(os.path.join(outdir, "protein_co-occurence.csv"))
    filtered_hit_df = hit_df[hit_df['LuxI_hmm'] >= 2]
    filtered_hit_df.to_csv(os.path.join(outdir, "multi-hit_strains.csv"))

    # iloc[:, 0] = the first column of the CSV (the assembly id index that
    # to_csv dumped without a header). Use a set for O(1) membership checks.
    multihit_assemblies = set(filtered_hit_df.iloc[:, 0].tolist())

    # Read the LuxI hmmsearch tabular output to get the set of protein_ids
    # that passed the E-value threshold. --tblout format: whitespace-delimited,
    # comments start with '#', target (protein_id) is column 0.
    luxi_hits_tbl = os.path.join(outdir, "LuxI_hits")
    luxi_hit_prot_ids = set()
    with open(luxi_hits_tbl, 'r') as tbl:
        for line in tbl:
            if line.startswith('#') or not line.strip():
                continue
            luxi_hit_prot_ids.add(line.split()[0])

    # Pull the LuxI hits from multi-hit assemblies out of all_prots.faa.
    # Sourcing from all_prots.faa (instead of LuxI_hmm.faa) gives clean unaligned
    # sequences without the alignment gaps introduced by hmmsearch -A / esl-reformat.
    with open(os.path.join(outdir, "all_prots.faa"), 'r') as infile:
        matching_records = []
        for record in SeqIO.parse(infile, 'fasta'):
            if record.id not in luxi_hit_prot_ids:
                continue
            asmbl_start = record.description.rfind('GCF')
            asmbl_end = record.description.rfind(']')
            asmbl_id = record.description[asmbl_start:asmbl_end]
            if asmbl_id in multihit_assemblies:
                matching_records.append(record)

        if matching_records:
            with open(os.path.join(outdir, "LuxI_multi.faa"), 'w') as outfile:
                SeqIO.write(matching_records, outfile, "fasta")
        else:
            print("No records matched the condition. Output file not created.")

    print('Process completed.')


else:
    print('1_LuxI_HMMsearch.py is being imported by another module.')
```
