## Supplementary File 1 for "Competing Inter-Domain Quorum-Sensing Systems Control Prophage Lysis-Lysogeny Decisions": 2_batch_NCBI_genome_query.py

```
#!/usr/bin/env python3
'''
Created on April 3, 2026

@author: Francis J Santoriello (Polished with Claude Opus 4.7 on June 30, 2026)

This script accepts a CSV file of RefSeq Accession IDs. It then:
    1. Downloads the GBFF file for each accession from NCBI Datasets.
    2. Pulls per-accession assembly metadata (organism, biosample, ...).
    3. Merges the per-accession metadata TSVs into a single summary table.

This script returns:
    1. One GBFF (.gbk) per Accession ID in the output directory.
    2. A merged assembly_metadata.tsv summarizing all accessions.
'''

import os
import argparse
import textwrap
import subprocess
import csv
import glob
import shutil

cwd = os.getcwd()  # current working directory


### User Input ###
class MyFormatter(argparse.ArgumentDefaultsHelpFormatter,
                  argparse.RawDescriptionHelpFormatter):
    pass

parser = argparse.ArgumentParser(
                prog='batch_NCBI_genome_query.py',
                formatter_class=MyFormatter,
                description=textwrap.dedent('''\
                    Created on April 3, 2026

                    @author: Francis J Santoriello

                    This script accepts a CSV file of RefSeq Accession IDs. It then:
    					1. Downloads the GBFF file for each accession from NCBI Datasets.
    					2. Pulls per-accession assembly metadata (organism, biosample, ...).
    					3. Merges the per-accession metadata TSVs into a single summary table.

					This script returns:
    					1. One GBFF (.gbk) per Accession ID in the output directory.
    					2. A merged assembly_metadata.tsv summarizing all accessions.

                    '''),
                usage='%(prog)s [options] <assembly_ids> \n\n '
                      'Dependencies: python 3, NCBI datasets\n')
parser.add_argument('assembly_ids',
                    help='CSV file of NCBI assembly IDs (one ID per row, first column).')
parser.add_argument('-o', '--outdir', type=str, default='output_gbffs',
                    help='Name of output directory for gbffs.')
parser.add_argument('-m', '--metadata-only', action='store_true',
                    help='Only retrieve and merge assembly metadata; skip GBFF downloads.')

args = parser.parse_args()


### Variables ###
id_file = args.assembly_ids
outdir = args.outdir
metadata_only = args.metadata_only


### Modules ###
def fetch_gbff(asmbl_id, outdir):
    """Download and stage the GBFF for one accession into outdir/{asmbl_id}.gbk.

    Uses argv-list subprocess calls (no shell) so accession strings can't be
    interpreted as shell metacharacters.
    """
    zip_path = f"{asmbl_id}.zip"

    # Download the zipped GBFF bundle for this accession
    subprocess.run(
        ["datasets", "download", "genome", "accession", asmbl_id,
         "--filename", zip_path, "--include", "gbff"],
        check=True,
    )

    # Unpack; -o overwrites silently so a partial prior run can't deadlock on a prompt
    subprocess.run(["unzip", "-o", zip_path], check=True)
    os.remove(zip_path)

    # NCBI Datasets nests the GBFF under ncbi_dataset/data/<accession>/*.gbff
    src_glob = os.path.join("ncbi_dataset", "data", asmbl_id, "*.gbff")
    matches = glob.glob(src_glob)
    if not matches:
        print(f"Warning: no GBFF found for {asmbl_id} at {src_glob}; skipping move.")
    else:
        shutil.move(matches[0], os.path.join(outdir, f"{asmbl_id}.gbk"))

    # Clean up the staging artifacts that `unzip` drops in cwd
    shutil.rmtree("ncbi_dataset", ignore_errors=True)
    for stray in ("md5sum.txt", "README.md"):
        try:
            os.remove(stray)
        except FileNotFoundError:
            pass


def fetch_metadata(asmbl_id, out_path):
    """Pipe `datasets summary ... | dataformat tsv ...` into out_path.

    Implemented as two chained Popens rather than a shell pipeline so neither
    asmbl_id nor out_path is interpreted by a shell.
    """
    with open(out_path, "w") as out_fh:
        summary = subprocess.Popen(
            ["datasets", "summary", "genome", "accession", asmbl_id,
             "--as-json-lines"],
            stdout=subprocess.PIPE,
        )
        formatter = subprocess.Popen(
            ["dataformat", "tsv", "genome",
             "--fields", "accession,organism-name,assminfo-biosample-*"],
            stdin=summary.stdout,
            stdout=out_fh,
        )
        # Allow `summary` to receive SIGPIPE if `formatter` exits early
        summary.stdout.close()
        formatter.wait()
        summary.wait()


### Main ###

if __name__ == '__main__':

    # Read assembly IDs from the first column of the input CSV
    with open(id_file, 'r', encoding='utf-8') as infile:
        reader = csv.reader(infile)
        asmbl_ids = [row[0] for row in reader]
        print(f"Extracted assembly ids: {asmbl_ids}")

    # Deduplicate so we don't redownload the same accession multiple times
    unique_asmbl_ids = list(set(asmbl_ids))

    # Create the output directory (no-op if it already exists)
    os.makedirs(outdir, exist_ok=True)

    # For each accession: download the GBFF (unless metadata-only) + per-accession metadata TSV
    for asmbl_id in unique_asmbl_ids:
        if not metadata_only:
            fetch_gbff(asmbl_id, outdir)
        fetch_metadata(asmbl_id, os.path.join(outdir, f"{asmbl_id}_metadata.tsv"))

    # Merge the per-accession metadata TSVs into a single summary table
    output_filename = os.path.join(outdir, "assembly_metadata.tsv")
    input_files = glob.glob(os.path.join(outdir, "*.tsv"))

    # Make sure the merged file isn't included as one of its own inputs
    if output_filename in input_files:
        input_files.remove(output_filename)

    # Sort for deterministic merge order
    input_files.sort()

    with open(output_filename, 'w') as outfile:
        first_file = True
        for file_path in input_files:
            with open(file_path, 'r') as infile:
                lines = infile.readlines()

            if not lines:
                continue  # Skip empty metadata files

            if first_file:
                # Keep the header from the first non-empty file
                outfile.writelines(lines)
                first_file = False
            else:
                # Drop the header on subsequent files
                outfile.writelines(lines[1:])

            # Remove the per-accession TSV now that it's been merged
            os.remove(file_path)

    print(f"Successfully merged {len(input_files)} files into {output_filename}")


else:
    print('batch_NCBI_genome_query.py is being imported by another module.')
```
