## Supplementary File 1 for "Competing Inter-Domain Quorum-Sensing Systems Control Prophage Lysis-Lysogeny Decisions": 3_LuxI_phage_association.py

```
#!/usr/bin/env python3
'''
Created on January 04, 2026

@author: Francis J Santoriello (Polished with Claude Opus 4.7 on June 30, 2026)

This script accepts a list of GenBank files and a protein FASTA file. It then:
    1. Searches each GenBank for CDS features whose protein_id matches a record
       in the input FASTA.
    2. Extracts the nucleotide sequence around each match (CDS +/- flank window)
       and writes it to a single nucleotide FASTA.

Expected input FASTA description format:
    "><protein_id> ..."

This script returns:
    1. output.fna (or whatever -o is set to) -- one record per matched CDS,
       containing `[CDS - flank, CDS + flank]` clipped to the contig boundaries.
'''

import os
import argparse
import textwrap
from Bio import SeqIO

cwd = os.getcwd()  # current working directory


### User Input ###
class MyFormatter(argparse.ArgumentDefaultsHelpFormatter,
                  argparse.RawDescriptionHelpFormatter):
    pass

parser = argparse.ArgumentParser(
                prog='LuxI_phage_association.py',
                formatter_class=MyFormatter,
                description=textwrap.dedent('''\
                    This script accepts a list of GenBank files and a protein FASTA file. It then:
                        1. Searches each GenBank for CDS features whose protein_id matches a record
                           in the input FASTA.
                        2. Extracts the nucleotide sequence around each match (CDS +/- flank window)
                           and writes it to a single nucleotide FASTA.

                    Expected input FASTA description format:
                        "><protein_id> ..."

                    This script returns:
                        1. output.fna (or whatever -o is set to) -- one record per matched CDS,
                           containing `[CDS - flank, CDS + flank]` clipped to the contig boundaries.
                    '''),
                usage='%(prog)s [options] <protein_faa> <gbs>\n\n '
                      'Dependencies: python 3, biopython\n')
parser.add_argument('protein_faa',
                    help='Fasta file of LuxI protein sequences')
parser.add_argument('gbs', nargs='*',
                    help='GenBank files for all alphaproteobacteria')
parser.add_argument('-w', '--window', type=int, default=20000,
                    help='Length (bp) of upstream and downstream flank around each CDS')
parser.add_argument('-o', '--outfile', type=str, default='output.fna',
                    help='Name of output file')

args = parser.parse_args()


### Variables ###
protfile = args.protein_faa
genbanks = args.gbs
window = args.window
outfile = args.outfile


### Modules ###
def fetch_flanks(infile, genomes, flank, output):
    '''
    For every protein id in `infile`, scan each GenBank in `genomes` for a CDS
    feature with a matching protein_id and write the nucleotide region
    `[CDS_start - flank, CDS_end + flank]` (clipped to the contig) into `output`.

    Header format for output records:
        >{protein_id} [{record description} -- {record id}|{assembly id}]
    '''
    # Collect target protein ids from the input FASTA descriptions.
    # See module docstring for the expected '...|<protein_id>/...' format.
    prot_ids = set()
    with open(infile, 'r') as prots:
        for record in SeqIO.parse(prots, 'fasta'):
            prot_ids.add(record.id)

    # Walk each GenBank once; emit a FASTA record for every matching CDS.
    with open(output, 'w') as out_fh:
        for gb_path in genomes:
            print(gb_path)
            for record in SeqIO.parse(gb_path, 'genbank'):
                print(record.id)

                # Pull the Assembly (GCF) dbxref once per record so we can tag
                # every emitted CDS header with its source assembly.
                assembly_link = next(
                    (s for s in record.dbxrefs if s.startswith('Assembly:')),
                    None,
                )
                assembly_id = (
                    assembly_link[len('Assembly:'):] if assembly_link else 'NA'
                )

                seq_len = len(record.seq)

                for feat in record.features:
                    if feat.type != 'CDS':
                        continue
                    if 'protein_id' not in feat.qualifiers:
                        continue
                    protein_id = feat.qualifiers['protein_id'][0]
                    if protein_id not in prot_ids:
                        continue

                    # Biopython locations are 0-based start, half-open end.
                    start = int(feat.location.start)
                    end = int(feat.location.end)

                    # Clip the flank window to the contig boundaries so we
                    # don't slice off the edge of the sequence.
                    upstream_start = max(0, start - flank)
                    downstream_end = min(seq_len, end + flank)

                    flank_seq = record.seq[upstream_start:downstream_end]

                    header = (f">{protein_id} "
                              f"[{record.description} -- {record.id}|{assembly_id}]")
                    out_fh.write(f"{header}\n{str(flank_seq)}\n")

                    print(f"Wrote {len(flank_seq)} bp sequence for "
                          f"{protein_id} from {record.id}.")

    print('Flank fasta file complete.')


### Main ###

if __name__ == '__main__':

    # Fetch flanking genomic sequences for each protein id
    fetch_flanks(protfile, genbanks, window, outfile)

    print('Process completed. <output.fna> file prepared for use with VIBRANT.')


else:
    print('LuxI_phage_association.py is being imported by another module.')
```
