## Supplementary File 1 for "Competing Inter-Domain Quorum-Sensing Systems Control Prophage Lysis-Lysogeny Decisions": 4_core_genome_MSA_from_BPGAout.py

```
#!/usr/bin/env python3
'''
Created on March 11, 2026

@author: Francis J Santoriello (Polished with Claude Opus 4.7 on June 30, 2026)

This script accepts the following files from BPGA version 1.3:
    1. list
    2. core.txt
    3. u_clusters.txt
    4. INPUT_all.faa

It then:
    1. Extracts all core genes for each genome from INPUT_all.faa
    2. Concatenates the core genes from each genome into a single string
    3. Aligns the concatenated core "genomes" with MUSCLE

This script returns:
    1. core_genome.faa  -- concatenated core proteins, one record per genome
    2. core_genome.aln  -- MUSCLE alignment of those concatenations
'''

import os
import argparse
import textwrap
import subprocess
import csv

from Bio import SeqIO

cwd = os.getcwd()  # current working directory


### User Input ###
class MyFormatter(argparse.ArgumentDefaultsHelpFormatter,
                  argparse.RawDescriptionHelpFormatter):
    pass

parser = argparse.ArgumentParser(
                prog='core_genome_MSA_from_BPGAout.py',
                formatter_class=MyFormatter,
                description=textwrap.dedent('''\
                    Created on March 11, 2026

                    @author: Francis J Santoriello

                    This script accepts the following files from BPGA version 1.3:
                        1. list
                        2. core.txt
                        3. u_clusters.txt
                        4. INPUT_all.faa

                    It then:
                        1. Extracts all core genes for each genome from INPUT_all.faa
                        2. Concatenates the core genes from each genome into a single string
                        3. Aligns the core genome strings

                    This script returns:
                        1. core_genome.faa
                        2. core_genome.aln
                    '''),
                usage='%(prog)s [options] <list_file> <core> <u_clust> <input_faa>\n\n '
                      'Dependencies: python 3, biopython, muscle\n')
parser.add_argument('list_file',
                    help='list file output by BPGA')
parser.add_argument('core',
                    help='core.txt file output by BPGA')
parser.add_argument('u_clust',
                    help='u_clusters.txt file output by BPGA')
parser.add_argument('input_faa',
                    help='INPUT_all.faa file output by BPGA')
parser.add_argument('-o', '--output', type=str, default='outdir',
                    help='Name of directory for output files')

args = parser.parse_args()


### Variables ###
def read_tsv(path):
    """Load a tab-delimited file as a list of row lists."""
    with open(path, newline='', encoding='utf-8') as fh:
        return list(csv.reader(fh, delimiter='\t'))


# BPGA's `list` file: one row per genome, columns identify the genome
genomes = read_tsv(args.list_file)

# BPGA's `core.txt`: one row per core cluster (column 0 is the cluster id)
cores = read_tsv(args.core)

# BPGA's `u_clusters.txt`: CD-HIT-style cluster table. Header rows start with
# 'C' in column 0; member rows have the cluster id in column 1 and the member
# protein id in column 8 (formatted as <genome>|<locus>).
clusters = read_tsv(args.u_clust)

input_faa = args.input_faa
outdir = args.output


### Modules ###
def extract_and_concat_core(genome_list, core_clusters, clust_file, INPUT_prot, outpath):
    '''
    Build, for each genome, the concatenation of its core-cluster proteins and
    write the result as a FASTA at `outpath/core_genome.faa`.

    Steps:
        1. For every (genome, core cluster) pair, find the member protein id
           in `clust_file` (BPGA's u_clusters.txt).
        2. Walk INPUT_all.faa once and replace each stored protein id with
           the actual amino-acid sequence.
        3. Emit one FASTA record per genome whose sequence is the
           concatenation of its core protein sequences, in core_clusters order.
    '''
    # core_prot_dict[genome_key][core_cluster_id] = prot_id (later: sequence)
    # Using a dict-of-dicts preserves a stable per-cluster ordering so the
    # concatenation order matches across genomes.
    core_prot_dict = {}

    # Step 1: locate the member protein id for each (genome, core cluster) pair
    for genome in genome_list:
        genome_key = f"{genome[0]}|{genome[1]}"
        core_prot_dict[genome_key] = {}

        print(f"Parsing {genome_key}...")

        for core_id in core_clusters:
            print(f"Searching for cluster {core_id[0]}...")

            for clust in clust_file:
                # Skip CD-HIT cluster-header rows (col 0 == 'C'); match on
                # genome prefix (col 8 before '|') AND cluster id (col 1).
                if (clust[0] != 'C'
                        and clust[8].split('|')[0] == genome[0]
                        and clust[1] == core_id[0]):
                    prot_id = clust[8]
                    print(f"Found {prot_id}.")
                    core_prot_dict[genome_key][core_id[0]] = prot_id
                    break  # one member per (genome, cluster) is enough

    # Step 2: invert the dict so we can resolve each FASTA record in O(1)
    # rather than rescanning core_prot_dict for every sequence in INPUT_all.faa.
    prot_id_to_location = {
        prot_id: (genome_key, core_id)
        for genome_key, cluster_map in core_prot_dict.items()
        for core_id, prot_id in cluster_map.items()
    }

    with open(INPUT_prot, 'r') as prot_db:
        for record in SeqIO.parse(prot_db, 'fasta'):
            location = prot_id_to_location.get(record.id)
            if location is None:
                continue
            genome_key, core_id = location
            core_prot_dict[genome_key][core_id] = str(record.seq)
            print(f"Replacing {core_id} value {record.id} with amino acid sequence...")

    # Step 3: write one FASTA record per genome -- header is the genome key,
    # sequence is the concatenation of its core protein sequences.
    with open(os.path.join(outpath, "core_genome.faa"), 'w') as outfile:
        print("Writing concatenated core genome sequences to output file...")
        for genome_key, cluster_map in core_prot_dict.items():
            header = f">{genome_key}"
            seq = "".join(cluster_map.values())
            outfile.write(f"{header}\n{seq}\n")


### Main ###

if __name__ == '__main__':

    # Create output directory (no-op if it already exists)
    os.makedirs(outdir, exist_ok=True)

    # Extract and concatenate core genes for each genome
    extract_and_concat_core(genomes, cores, clusters, input_faa, outdir)

    # Align the per-genome core concatenations with MUSCLE.
    # argv-list call (no shell) so paths can't be reinterpreted as shell syntax.
    print('Aligning core genome sequences with muscle...')
    subprocess.run(
        ["muscle",
         "-align", os.path.join(outdir, "core_genome.faa"),
         "-output", os.path.join(outdir, "core_genome.aln")],
        check=True,
    )

    print('Process completed.')


else:
    print('core_genome_MSA_from_BPGAout.py is being imported by another module.')
```
