## Supplementary File 1 for "Competing Inter-Domain Quorum-Sensing Systems Control Prophage Lysis-Lysogeny Decisions": prot_clustering.Rmd

````
---
title: "20260105 Alphaproteobacteria phage Hth-Xre/LuxI/LuxR Module Clustering"
author: "Frank Santoriello"
date: "2026 APR 09"
output: 
  html_notebook: default
  html_document: default
  pdf_document: default
---
```{r setup, include=FALSE}
knitr::opts_chunk$set(echo = TRUE)

#Set working directory to the location of the data.
knitr::opts_knit$set(root.dir = "~/")
```

```{r}
getwd()
```

This is an [R Markdown](http://rmarkdown.rstudio.com) Notebook. When you execute code within the notebook, the results appear beneath the code. 

Try executing this chunk by clicking the *Run* button within the chunk or by placing your cursor inside it and pressing *Cmd+Shift+Enter*. 

LOAD PACKAGES
```{r}
library(tidyverse)
library(pheatmap)
```

IMPORT AND FORMAT DATA
```{r}
# Import and format distance matrices created in Geneious

luxI_dist_mat <- read.csv("phage_luxI_AAident.csv")
luxR_dist_mat <- read.csv("phage_luxR_AAident.csv")
xre_dist_mat <- read.csv("phage_XRE_AAident.csv")
hth_dist_mat <- read.csv("phage_HTH_AAident.csv")


luxI_dist_mat <- as_tibble(luxI_dist_mat)
luxR_dist_mat <- as_tibble(luxR_dist_mat)
xre_dist_mat <- as_tibble(xre_dist_mat)
hth_dist_mat <- as_tibble(hth_dist_mat)

luxI_dist_mat %>%
  mutate(across(where(is.numeric), ~replace_na(.x, 100))) -> luxI_dist_mat

luxR_dist_mat %>%
  mutate(across(where(is.numeric), ~replace_na(.x, 100))) -> luxR_dist_mat

xre_dist_mat %>%
  mutate(across(where(is.numeric), ~replace_na(.x, 100))) -> xre_dist_mat

hth_dist_mat %>%
  mutate(across(where(is.numeric), ~replace_na(.x, 100))) -> hth_dist_mat


```

Generate heatmap
```{r}

# Build heatmap

thresholdValue = 30
breaksList = seq(30, 100, by = 5)

luxI_dist_mat %>%
  select(!X) %>%
  mutate(across(everything(), ~if_else(. < thresholdValue, 0, .))) %>%
  pheatmap(.,color = colorRampPalette(c('white','black'))(length(breaksList)),
        breaks = breaksList,
        cellwidth = 5, cellheight = 5,
        cluster_rows = FALSE, cluster_cols = FALSE,
        show_rownames = FALSE,
        border_color = 'white',
        fontsize_col = 5)

luxR_dist_mat %>%
  select(!X) %>%
  mutate(across(everything(), ~if_else(. < thresholdValue, 0, .))) %>%
  pheatmap(.,color = colorRampPalette(c('white','black'))(length(breaksList)),
        breaks = breaksList,
        cellwidth = 5, cellheight = 5,
        cluster_rows = FALSE, cluster_cols = FALSE,
        show_rownames = FALSE,
        border_color = 'white',
        fontsize_col = 5)

xre_dist_mat %>%
  select(!X) %>%
  mutate(across(everything(), ~if_else(. < thresholdValue, 0, .))) %>%
  pheatmap(.,color = colorRampPalette(c('white','black'))(length(breaksList)),
        breaks = breaksList,
        cellwidth = 5, cellheight = 5,
        cluster_rows = FALSE, cluster_cols = FALSE,
        show_rownames = FALSE,
        border_color = 'white',
        fontsize_col = 5)

hth_dist_mat %>%
  select(!X) %>%
  mutate(across(everything(), ~if_else(. < thresholdValue, 0, .))) %>%
  pheatmap(.,color = colorRampPalette(c('white','black'))(length(breaksList)),
        breaks = breaksList,
        cellwidth = 5, cellheight = 5,
        cluster_rows = FALSE, cluster_cols = FALSE,
        show_rownames = FALSE,
        border_color = 'white',
        fontsize_col = 5)


```

Add a new chunk by clicking the *Insert Chunk* button on the toolbar or by pressing *Cmd+Option+I*.

When you save the notebook, an HTML file containing the code and output will be saved alongside it (click the *Preview* button or press *Cmd+Shift+K* to preview the HTML file). 

The preview shows you a rendered HTML copy of the contents of the editor. Consequently, unlike *Knit*, *Preview* does not run any R code chunks. Instead, the output of the chunk when it was last run in the editor is displayed.
````
