## Supplementary material for "Competing Inter-Domain Quorum-Sensing Systems Control Prophage Lysis-Lysogeny Decisions": Notice of HHMI Publication Policy

Howard Hughes Medical Institute

### NOTICE OF PRE-EXISTING CONDITIONS, REQUIREMENTS AND LICENSES FOR ARTICLE SUBMISSION

The article submitted together with this notice is subject to the Immediate Access to Research policy of the Howard Hughes Medical Institute ("HHMI"). In accordance with this policy: (i) a preprint of this article either has been, or will be, deposited on a preprint server under a Creative Commons Attribution 4.0 International (CC BY 4.0) license and (ii) an additional author-published revised version of this article incorporating peer review feedback and/or new results or analysis either has been, or prior to journal publication will be, deposited on a preprint server under a CC BY 4.0 license. In addition, a non-exclusive CC BY 4.0 license to this article has been granted to the public and HHMI has a sublicensable, non-exclusive license to this article.

**THIS ARTICLE IS SUBMITTED FOR REVIEW AND ACCEPTANCE SUBJECT TO THESE PRE-EXISTING CONDITIONS, REQUIREMENTS AND LICENSES.**

If you have any concerns with any of these pre-existing conditions, requirements or licenses, please contact the corresponding author immediately.
